# Driving and suppressing the human language network using large language models

**DOI:** 10.1101/2023.04.16.537080

**Authors:** Greta Tuckute, Aalok Sathe, Shashank Srikant, Maya Taliaferro, Mingye Wang, Martin Schrimpf, Kendrick Kay, Evelina Fedorenko

**Author notes:** Corresponding authors: Greta Tuckute and Evelina Fedorenko.

## Abstract

Transformer models such as GPT generate human-like language and are highly predictive of human brain responses to language. Here, using fMRI-measured brain responses to 1,000 diverse sentences, we first show that a GPT-based encoding model can predict the magnitude of brain response associated with each sentence. Then, we use the model to identify new sentences that are predicted to drive or suppress responses in the human language network. We show that these model-selected novel sentences indeed strongly drive and suppress activity of human language areas in new individuals. A systematic analysis of the model-selected sentences reveals that surprisal and well-formedness of linguistic input are key determinants of response strength in the language network. These results establish the ability of neural network models to not only mimic human language but also noninvasively control neural activity in higher-level cortical areas, like the language network.

## Introduction

Reading and understanding this sentence engages a set of left-lateralized frontal and temporal brain regions. These interconnected areas, or the ‘language network’ (e.g., ^1–3^), a) support both comprehension and production of spoken, written, and signed linguistic utterances (e.g., ^4,2,5–10^) across diverse languages ^11,12^; b) are highly selective for language relative to diverse non-linguistic inputs (e.g., ^13–17^; see ^18^ for a review); c) are sensitive to linguistic structure at many levels (e.g., ^2,14,6,19–21,18,10^); and d) are causally important for language such that their damage leads to linguistic deficits ^22–28^. However, many aspects of the representations and algorithms that support language comprehension remain unknown.

Over the last few years, artificial neural networks for language have emerged as in-silico models of language processing. These large language models (LLMs) can generate coherent text, answer questions, translate between languages, and perform sophisticated language comprehension tasks (e.g., ^29–35^). Strikingly, despite the fact that the LLMs were not developed with the goal of modeling human language processing, some of these models (especially the unidirectional Transformer architectures ^36^) have a remarkable capacity to mimic human language behavior (e.g., ^37–40^) and predict brain activity during language processing (e.g., ^41–53^). However, despite LLMs being today’s most quantitatively accurate models of language processing, there has been no attempt to test whether LLMs can *causally control* language responses in the brain (e.g., ^41–53^). By ‘causal control’ we mean the use of models to make quantitative predictions about a neural target (a cell or a brain area/network) and subsequently using those predictions to successfully modulate neural activity in the target in a ‘closed-loop’ manner.

Recent work in visual neuroscience has shown that artificial neural network models of image recognition can causally intervene on the non-human primate visual system by generating visual stimuli that modulate activity in different regions of the ventral visual pathway ^54–57^. In this work, we ask whether similar model-based control is feasible for the higher-level cognitive domain of language: can we leverage the predictive power of LLMs to identify new stimuli to maximally drive or suppress brain responses in the language network of new individuals? This question taps into two key aspects of the generalization ability of LLMs: i) do LLMs capture features of language representations that *generalize across humans*? and ii) do LLMs have the capacity to predict brain responses to model-selected stimuli that *extend beyond the distribution of naturally occurring linguistic input*? We demonstrate that model-selected stimuli drive and suppress brain responses in the language network of new individuals, establishing the ability of brain-aligned LLMs to non-invasively control areas implicated in higher-level cognition. We then leverage sentence-level brain responses to a broad distribution of linguistic input to ask what kinds of linguistic input the language network is most responsive to. In a large-scale behavioral experiment, we collect rating norms for ten sentence properties and use these norms to characterize the language network’s preferred stimuli.

## Results

### Model-selected sentences control language network responses

Our aim is to test whether current models of the human language network are capable of driving and suppressing brain responses in these higher-level cognitive brain areas. We developed an encoding model of the left hemisphere (LH) language network in the human brain with the goal of identifying new sentences that would activate the language network to a maximal or minimal extent. The model takes as input last-token sentence embeddings from GPT2-XL ^36^ (previously identified as the most brain-aligned language model ^43^; layer 22, see <u>SI 6A</u> for the cross-validated analysis that led to this choice) and was trained, via ridge regression, to predict the average LH language network’s (functionally defined ^2^) BOLD response (henceforth language network’s response; <u>Methods; Definition of ROIs</u>). The BOLD responses were acquired from 5 *train* participants who read a set of 1,000 diverse, corpus-extracted sentences (*baseline set*) (2 sessions each, n=10 sessions total; <u>Methods; Encoding model development</u>) (**Figure 1A**). The encoding model achieved a prediction performance of r=0.38 (noise ceiling is r=0.56; <u>SI 6</u>) when evaluated on held-out sentences within the *baseline set* (SE over five splits=0.16, all five p-values <.001; <u>SI 6A)</u>. To ensure that the encoding model performance did not hinge on specific experimental decisions, we confirmed that the model maintained high predictivity performance on held-out sentences when changing the procedure for obtaining sentence embeddings (the average of all tokens in the sentence; <u>SI 6B</u>) and even using sentence embeddings from a different LLM architecture (a bidirectional-attention Transformer model, BERT-large; <u>SI 6C</u>). Further, the encoding model also achieved relatively high predictive performance on anatomically, rather than functionally, defined language regions, although predictivity was lower (<u>SI 6D</u>).

**Figure 1.**
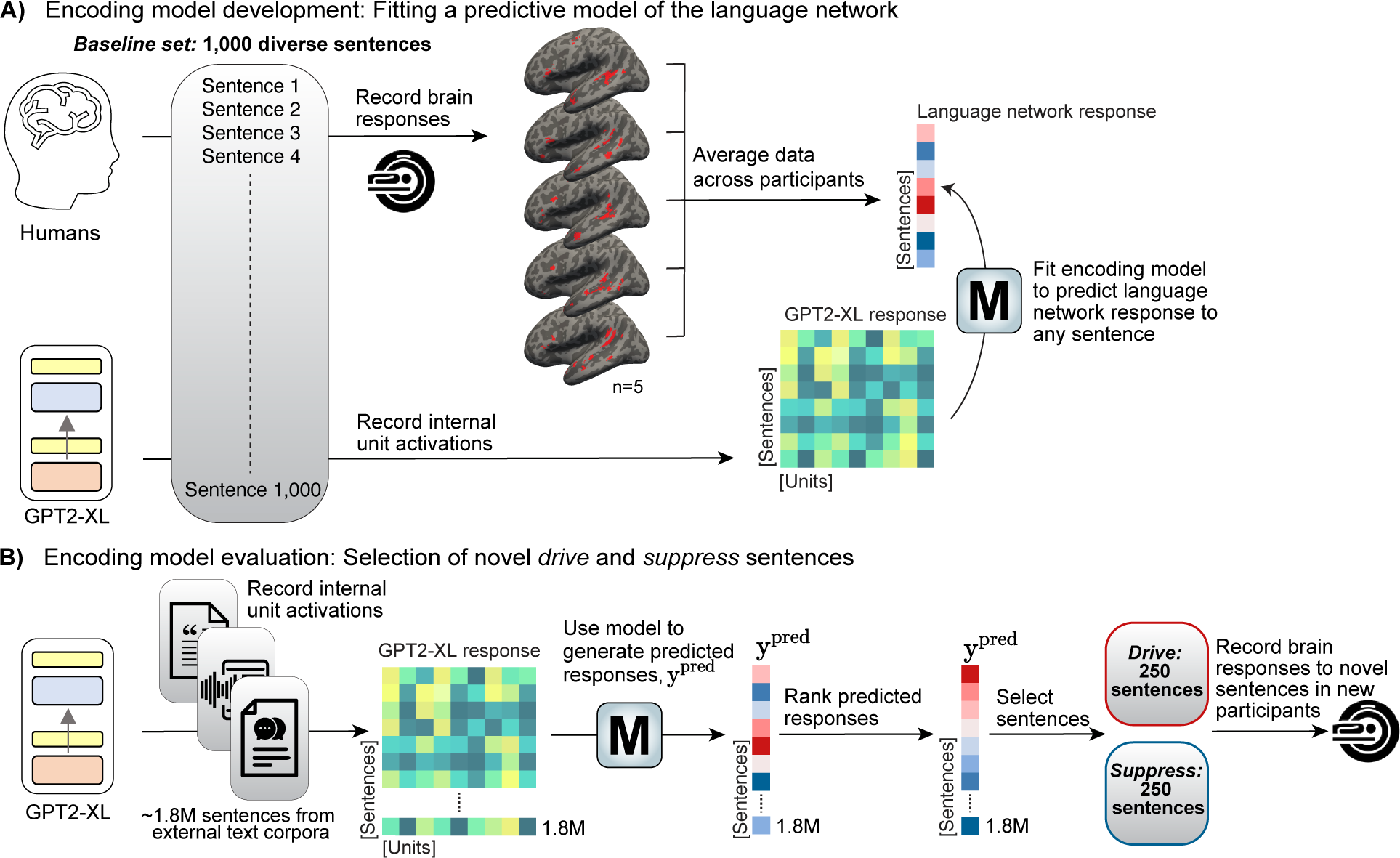
Overview of the procedure for encoding model development and stimulus selection for evaluation. **A)** We developed an encoding model (**M**) of the left hemisphere (LH) language network in the human brain with the goal of identifying novel sentences that activate the language network to a maximal or minimal extent (<u>Methods; Encoding model development</u>). Five participants (*train* participants) read a large sample (n=1,000) of 6-word corpus-extracted sentences, the *baseline set* (sampled to maximize linguistic diversity; <u>SI 1</u>), in a rapid, event-related design while their brain activity was recorded using fMRI. Blood-oxygen-level-dependent (BOLD) responses from voxels in the LH language network were averaged within each *train* participant and averaged across participants to yield an average language network response to each of the 1,000 *baseline set* sentences. We trained a ridge regression model from the representations of the unidirectional-attention Transformer language model, GPT2-XL (identified as the most brain-aligned language base model in Schrimpf et al. ^43^), to the 1,000 averaged fMRI responses. Given that GPT2-XL can generate a representation for any sentence, the encoding model (**M**) can predict the LH language network response for arbitrary sentences. To select the top-performing layer for our encoding model, we evaluated all 49 layers of GPT2-XL and selected the layer that had highest predictivity performance on brain responses to held-out *baseline set* sentences (layer 22; <u>SI 6A</u>). **B)** To evaluate the encoding model (**M**), we identified a set of sentences to activate the language network to a maximal extent (*drive* sentences) or minimal extent (*suppress* sentences) (<u>Methods; Encoding model</u> <u>evaluation</u>). To do so, we obtained GPT2-XL embeddings for ∼1.8 million sentences from diverse, large text corpora, generated predicted language network responses, and ranked these responses to select the sentences that are predicted to increase or decrease brain responses relative to the *baseline set*. Finally, we collected brain responses to these novel sentences in new participants (*evaluation* participants).

To identify sentences that would elicit a desired (high or low) level of activation in the language network, we searched across ∼1.8 million sentences from 9 diverse large-scale text corpora (**Figure 1B**). We identified a set of 250 sentences that were predicted to elicit maximally strong activity in the language network (*drive* sentences; e.g., “Turin loves me not, nor will.” or “People on Insta Be Like, “Gross!””) and 250 sentences that were predicted to elicit minimal activity in the language network (*suppress* sentences; e.g., “We were sitting on the couch.” or “Inside was a tiny silver sculpture.”). We evaluated our encoding model by recording brain responses to these new *drive* and *suppress* sentences in new participants (denoted as *evaluation* participants) (note the fully independent procedure using both new stimuli and participants; for evidence that the new *drive* and *suppress* sentences differ from the *baseline* sentences, see <u>SI</u> <u>10</u> and <u>11</u>).

We collected fMRI responses to the *drive* and *suppress* sentences in an event-related, single-trial paradigm in three new participants (3 sessions each, n=9 sessions total; <u>Methods;</u> <u>Encoding model evaluation</u>). The *drive* and *suppress* sentences were randomly interspersed among the 1,000 *baseline* sentences. **Figure 2B** shows the average responses for the n=3 *evaluation participants* for the *drive, suppress,* and *baseline* sentence conditions. The *drive* sentences yielded significantly higher responses than the *suppress* sentences (β=0.57, t=15.94, p<.001 using linear mixed effect modeling, see <u>Methods; Statistical analyses</u> and <u>SI 18</u>). The *drive* sentences also yielded significantly higher responses than the *baseline* sentences (β=0.27, t=9.72, p<.001) with the evoked BOLD signal being 85.7% higher for the *drive* condition relative to *baseline* (quantified using non-normalized BOLD responses; <u>SI 12A</u>). Finally, the *suppress* sentences yielded lower responses than the baseline sentences (β=-0.29, t=-10.44, p<.001) with the evoked BOLD responses being 97.5% lower for the *suppress* condition relative to *baseline* (<u>SI 12A</u>). In summary, we trained an encoding model to generate predictions about the magnitude of activation in the language network for a new set of sentences and then ‘closed the loop’ by collecting brain data for these new sentences in new participants to demonstrate that these sentences modulate brain responses as predicted. We note that although we trained the encoding model using the responses in the LH language network as a whole, the five individual LH language fROIs showed highly correlated responses across the *baseline* set (<u>SI 4</u>; and <u>Results; Language regions exhibit high stimulus-related</u> <u>activity</u>) and similar condition-level responses to the *drive*, *suppress*, and *baseline* sentences (<u>SI</u> <u>15F</u>) (see <u>SI 15G</u> for evidence that this pattern of responses to *drive, suppress*, and *baseline* sentences is not ubiquitously present across the brain). These inter-fROI similarities align well with past work showing similar modulation of the different language areas by diverse linguistic manipulations (e.g., ^10,11,14,15,17,21,58–60^).

**Figure 2.**
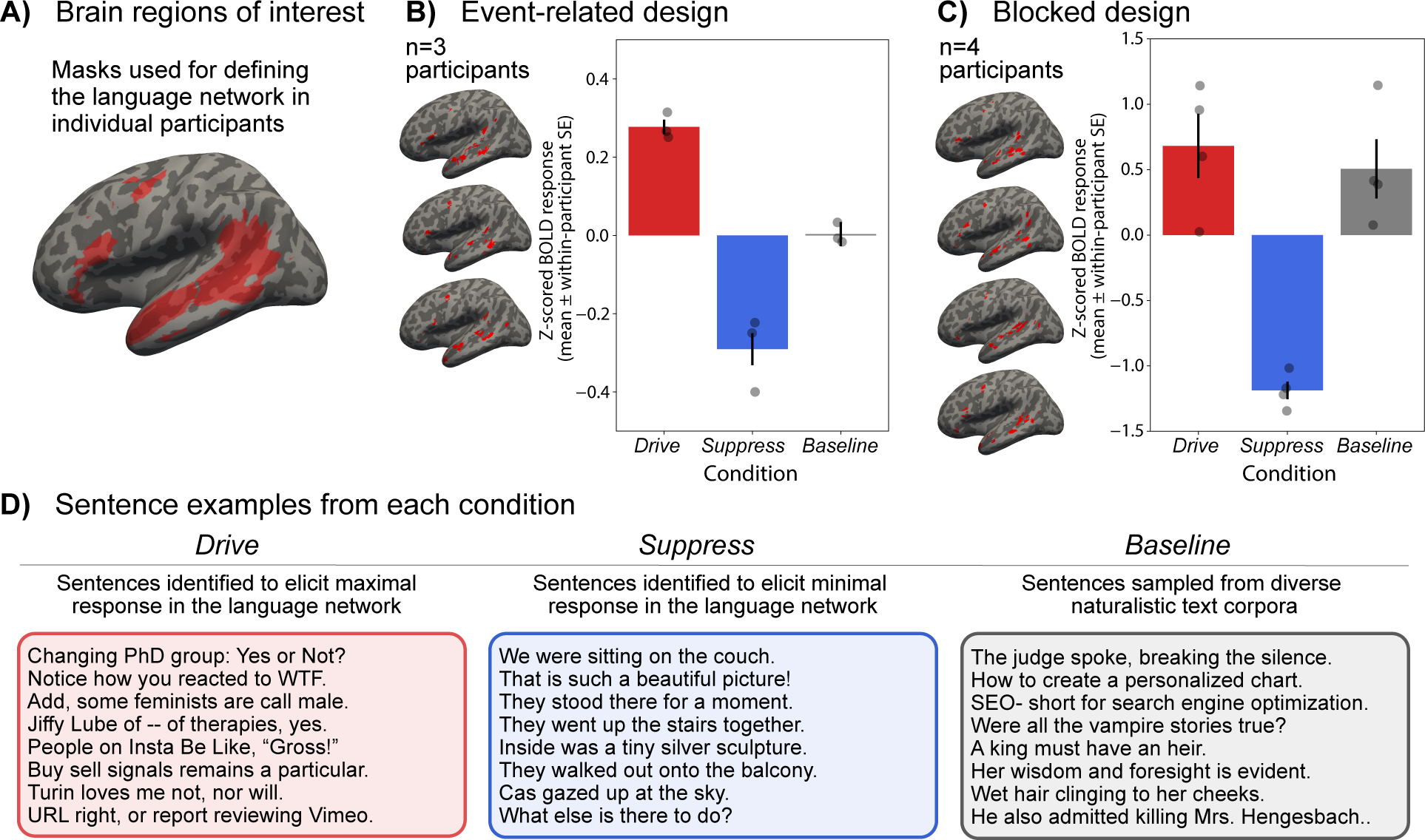
Model-selected sentences successfully drive and suppress responses in the language network. **A)** We used our encoding model to select sentences that would elicit maximal response (*drive* sentences) or minimal response (*suppress* sentences) in the functionally defined language network. To define functional ROIs, we used demarcations (‘language parcels’; shown on the surface-inflated MNI152 template brain) within which most or all individuals in prior studies showed activity for the language localizer contrast in large samples (e.g., ^2,3^). We defined the LH language network as regions within the borders of these five parcels that were activated (top 10%) in the functional localizer acquired for each participant (see brain visualizations in B and C). **B)** The average language network fMRI response across, respectively, 250 *drive,* 250 *suppress*, and 1,000 *baseline* sentences for n=3 *evaluation* participants, collected in an event-related, single-trial fMRI paradigm. In both B and C, individual points show the average of each condition per participant. fMRI responses were z-scored session-wise (see <u>SI 12A</u> for responses without normalization; no key patterns are affected). The evoked BOLD response was 85.7% higher for *drive* relative to *baseline* and 97.5% lower for *suppress* relative to *baseline* (<u>SI 12A)</u>. Error bars show within-participant standard error of the mean. The brain illustrations show the functionally defined language network in the participants of interest on the surface-inflated brain, visualized in Freeview. For surface projections, volumetric data (in MNI IXI549Space; SPM12 ^197^) were registered to FreeSurfer’s CVS35 (combined volumetric and surface-based (CVS)) in the MNI152 space using *mri_vol2surf* in FreeSurfer v7.3.2 ^198^ with a projection distance of 1.5mm and otherwise default parameters. **C)** The average language network fMRI response across, respectively, 240 *drive,* 240 *suppress*, and 240 *baseline* sentences (randomly sampled from the superset of 250/250 *drive/suppress* sentences and 1,000 *baseline* sentences) for n=4 *evaluation* participants, collected in a blocked fMRI paradigm. The evoked BOLD response was 12.9% higher for *drive* relative to *baseline* and 56.6% lower for *suppress* relative to *baseline* (<u>SI 12B)</u>. **D)** Example sentences from each condition.

To further validate the robustness of responses to the *drive* and *suppress* sentences, we collected brain data for a large subset of the *drive, suppress,* and *baseline* stimuli in a traditional blocked fMRI design, where *drive, suppress*, and *baseline* sentences are blocked into groups, in four additional participants (1 session each, n=4 sessions total; <u>Methods; fMRI experiments</u>). The results mirrored those from the event-related experiment: the *drive* sentences yielded the highest response followed by *baseline* sentences (the evoked BOLD response was 12.9% higher for *drive* relative to *baseline*; 56.6% lower for *suppress* relative to *baseline*; <u>SI 12B</u>). Hence, independent of experimental design (event-related vs. blocked) and b) modeling procedure (single-trial modeling vs. condition-level modeling), the brain responses to the *drive* sentences were high relative to the baseline sentences, and the responses to the *suppress* sentences were low relative to the baseline sentences.

For a final examination of model-guided stimulus selection, we explored an alternative approach to selecting *drive*/*suppress* sentences: the *modify* approach where, instead of searching within existing text corpora, we used gradient-based modifications to transform a random sentence into a novel sentence predicted to elicit high or low fMRI responses (<u>SI 16A</u>) and collected responses to these novel sentences in two participants (event-related design, n=6 sessions total). We found that this exploratory *modify* approach was able to drive responses by 57% relative to *baseline*, but failed to suppress responses, most likely because the resulting *modify* stimuli were often akin to word lists which the encoding model was not trained on (<u>SI 16B</u>).

### Model captures most explainable variance in new participants

In the previous section, we examined predictivity at the condition level (*drive* vs. *suppress* vs. *baseline*). Here, we sought to evaluate the accuracy of the predictions from the encoding model at the level of individual sentences. To do so, we turned to the event-related experiment (**Figure 2B**), which allows us to estimate sentence-level brain responses to 1,500 sentences for each of the three *evaluation* participants.

**Figure 3** shows the model-predicted versus observed brain responses in the language network (n=3 *evaluation* participants). These participants were not used to train the encoding model and hence allow us to estimate encoding model predictivity performance in held-out participants and held-out sentences. Across the full set of 1,500 *baseline, drive,* and *suppress* sentences, we obtained a Pearson correlation of 0.43 (dof=1498, p<.001, t=18.60, SE=0.02) between predicted and observed brain responses. Because the *drive* and *suppress* sentences were designed to elicit high or low brain responses respectively, one might expect that the correlation might be unduly driven by these two conditions. Therefore, we isolated the set of n=1,000 naturalistic, corpus-extracted *baseline* sentences and obtained a correlation of 0.30 (dof=998, p<.001, t=9.88, SE=0.03). Hence, the encoding model was able to predict a substantial and statistically significant amount of variance in brain responses in new participants to both naturalistic, sentences that fall within the distribution of the training (*baseline*) set and out-of-distribution sentences (*drive/suppress* set), for which encoding model predictions (x-axis in **Figure 3**) extend far beyond the training set distribution.

**Figure 3.**
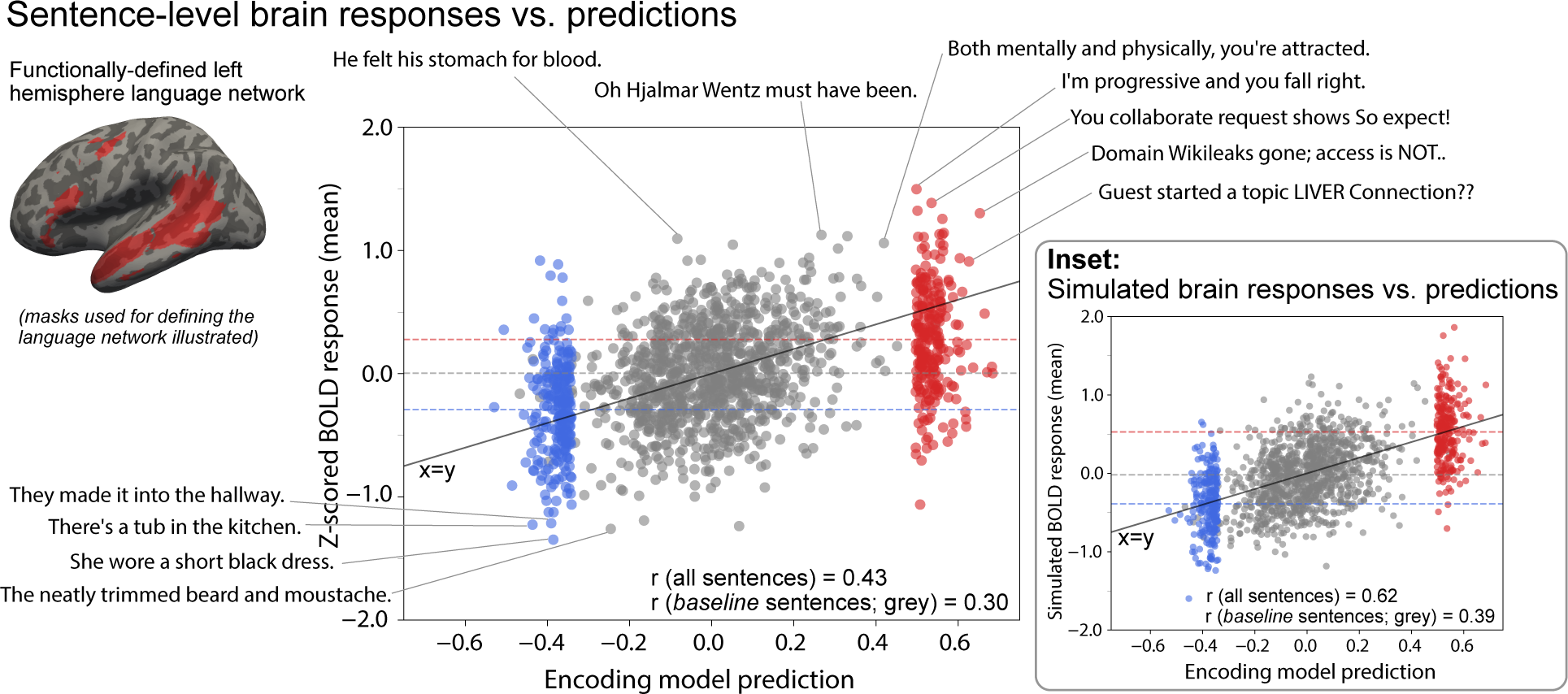
The encoding model maintains high predictive performance for brain responses from three new participants to out-of-distribution-sentences. Sentence-level brain responses as a function of the predicted responses along with sentence examples. Predicted brain responses were obtained from the encoding model (x-axis). The observed brain responses (y-axis) are the average of n=3 *evaluation* participants’ language network responses (illustrated for individual participants in <u>SI 13</u>). The blue points represent the *suppress* sentences, the grey points represent the *baseline* sentences, and the red points represent the *drive* sentences. The *suppress* and *drive sentences* were selected to yield respectively low or high brain responses and are therefore clustered on the low and high end of the prediction axis (x-axis). Dashed horizontal lines show the mean of each condition. **Inset:** Simulated sentence-level brain responses as a function of predicted responses. Predicted brain responses were obtained from the encoding model (x-axis). The simulated brain responses (y-axis) were obtained by sampling from a noise distribution representing the empirical inter-participant variability. This plot illustrates the maximum possible predictive performance, given inter-participant variability and fMRI measurement noise.

To better interpret the accuracy of sentence-level predictions, we quantified the maximal possible prediction performance by treating inter-participant variability as “noise” that cannot be predicted by a computational model. The goal here is to assess how well our model predicts brain activity at the group level, taking into account irreducible variance due to inter-participant variability and measurement noise. First, we computed the empirical variability in participants’ responses to the 1,500 sentences. Next, we simulated response noise for each participant using the empirical variability across participants (drawing samples from a Gaussian distribution with zero mean and the empirical inter-participant standard deviation). For each sentence, simulated response noise was added to the encoding model’s predicted response (x-axis in **Figure 3**) and responses were then averaged across participants. This simulation provides an estimate of the maximum possible encoding model prediction performance.

**Figure 3 inset** shows these simulated brain responses versus predicted responses. In these simulations, the Pearson correlation was 0.62 (dof=1498, p<.001, t=30.85, SE=0.02) across all 1,500 sentences (observed: r=0.43, i.e., 69.4% of the theoretically obtainable correlation), and 0.39 (dof=998, p<.001, t=13.32, SE=0.03) across the 1,000 *baseline* sentences (observed: r=0.30, i.e., 76.9% of the theoretically obtainable correlation). These results show that due to inter-participant variability in fMRI measurements, even a perfect model can achieve only r=0.62 predictive performance. Although our model is not perfect, the performance level suggests that the model successfully captures much of the neurally relevant variance in responses to individual sentences.

### Language regions exhibit high stimulus-related activity

Having established that model-selected stimuli could indeed drive and suppress brain responses in the language network of new individuals (**Figure 2, 3**), our next goal was to investigate what kinds of linguistic input the LH language network is most responsive to. Before delving into that investigation, however, we wanted to assess that the LH language regions show reliable responses to and track properties of linguistic stimuli. We also wanted to assess the similarity among the language fROIs in their fine-grained linguistic preferences in order to decide whether it may be worth to examine the fROIs separately in addition to examining the language network as a whole.

First, we quantified noise ceilings for the language regions along with a set of control brain regions (**Figure 4A**). A noise ceiling (NC) for a brain region is a measure of stimulus-related response reliability and is typically expressed in terms of the fraction of variance that can be attributed to the stimulus rather than to measurement noise. Standard approaches for NC estimation leverage repeated stimulus presentations, with the core assumption that repeated presentations should yield the same brain response (e.g., ^61–63^). Because in the current study, each sentence was presented only once to a given participant (for the motivation for this design choice and details of the procedure, see <u>Methods</u> and <u>Discussion</u>), we developed a procedure for NC estimation that makes use of the repeated presentations of the same sentence *across participants*, allowing for estimation of reliability in single-repetition paradigms (<u>SI 5</u>). Using this procedure, we computed NCs based on the brain responses to the 1,000 *baseline* sentences for the n=5 *train* participants in language regions and a set of control regions (**Figure 4A**). In particular, we examined two large-scale brain networks that have been linked to high-level cognitive processing—the multiple demand (MD) network ^64–68^ and the default mode network (DMN) ^69–73^—which we defined using independent functional localizers (see <u>SI 15</u> for details) (**Figure 4A**). For additional comparison, we examined a set of anatomical parcels ^74^ that cover a large fraction of the cortical surface (<u>SI 8</u>).

**Figure 4.**
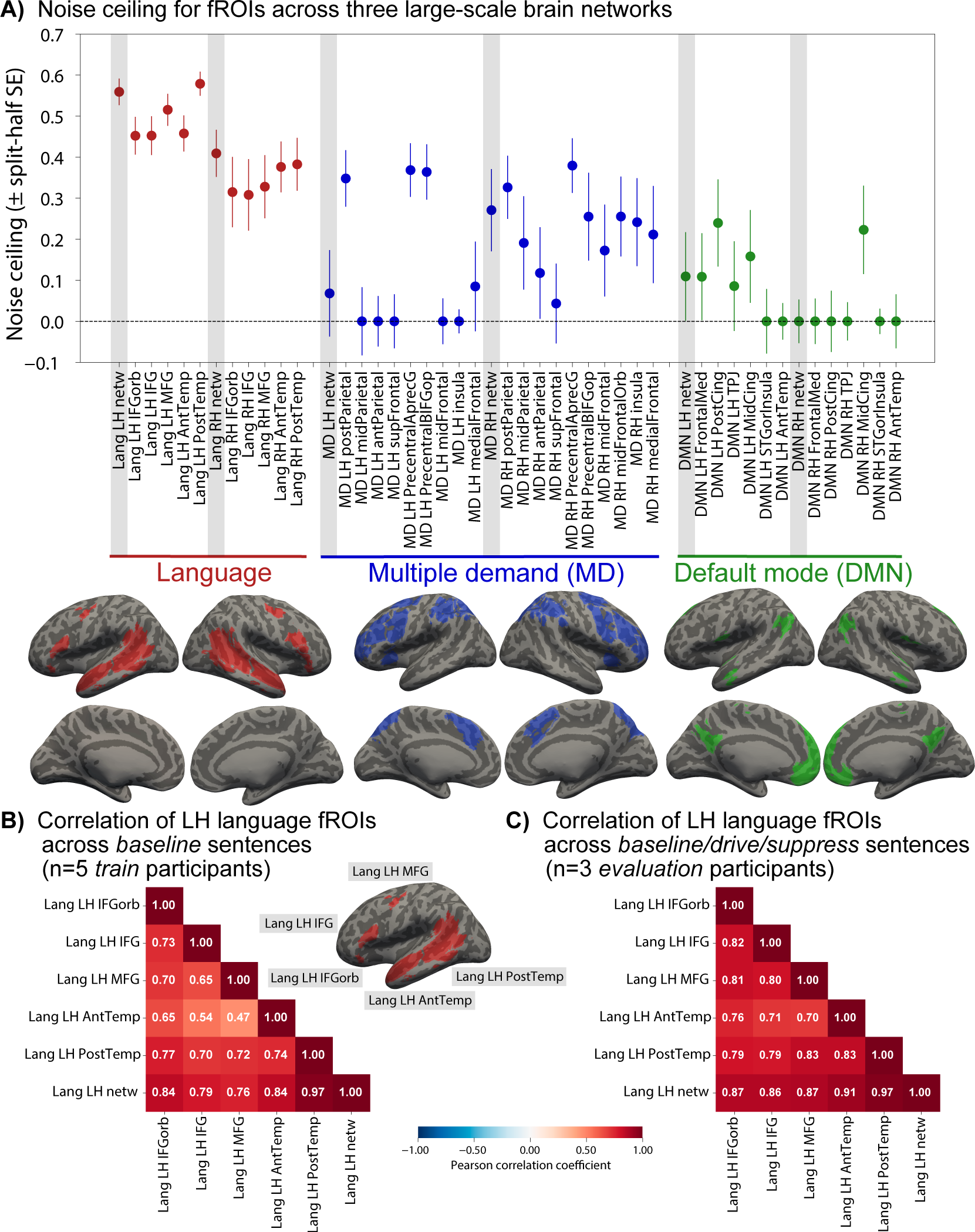
Left-hemisphere language regions show a high degree of stimulus-related activity for linguistic input relative to other brain areas and the left-hemisphere language regions show functionally similar responses. **A)** We quantified the noise ceiling (NC), a measure of stimulus-related response reliability, across all functionally defined ROIs in the language network (red), multiple demand (MD) network (blue), and the default mode network (DMN; green). For the language network, we defined 10 such fROIs (along with the “Language LH/RH network” fROI which is the mean across all voxels in the fROIs within the network and hemisphere, i.e., yielding 12 ROIs in total), for the MD network, we defined 21 fROIs (note that one participant did not show a response to the MD localizer in the MD LH midFrontalOrb ROI, and hence this ROI was excluded in the NC computation), and for the DMN network, we defined 14 fROIs. Grey shaded areas indicate the network-level fROIs. The dots show the NC estimate computed across n=1,000 *baseline* sentences across n=5 *train* participants, for each of the ROIs. Error bars show the NC reliability quantified as the standard error over NC values computed from 1,000 splits of the data (<u>SI 5B</u>). The brain illustrations show the anatomical parcels (demarcations) that were used to constrain the selection of participant-specific fROIs for each network in the surface-inflated MNI152 template brain. **B)** The Pearson correlation matrix computed over n=1,000 *baseline* sentences for the average of n=5 *train* participants. The first five rows/columns show the five core LH language fROIs (IFGorb, IFG, MFG, AntTemp, and PostTemp; <u>Methods; Definition of ROIs).</u> The sixth row/column shows the full LH language network consisting of the average of the voxels from the five fROIs; these values show how representative the language network as a whole is of each of the five fROIs. **C)** Same as in panel B, but for the n=1,500 *drive*/*suppress*/*baseline* sentences for the average of n=3 *evaluation* participants (derived using the main, *search* approach). Correlation matrices for individual participants are shown in <u>SI 4</u>.

Prior studies have demonstrated high consistency of responses in language regions across participants using naturalistic story-listening paradigms ^75–78^. In line with those studies, we found that in our single-sentence paradigm, language regions were also characterized by high NC. The ceiling values were higher than those observed in the two other functional networks (**Figure 4A**) and in anatomical areas across the brain (including anatomical areas that fall in spatially similar locations to the language areas, which provides further evidence for the advantages of functional localization ^79,80^; <u>SI 8</u>). ^75–78^In particular, for the LH language areas, the NC was estimated to be r=0.56 (split-half standard error (SE)=0.03), i.e., ∼31% of the variance in the responses of these areas at the group-level can be considered “true”, stimulus-related, signal. For comparison: for the MD network, the NC was estimated to be r=0.07 (SE=0.11) (for the LH MD areas) and r=0.27 (SE=0.10) (for the RH MD areas; see ^77^ for convergent evidence from a different approach), and for the DMN, the NC was estimated to be r=0.11 (SE=0.11) (for the LH DMN areas) and r=0 (SE=0.05) (for the RH DMN areas), The LH language network NC values were significantly higher than the NC in each of these four networks—LH and RH MD and DMN (dof=1999, all four p<.001, all four t>126 via Bonferroni-corrected one-sided t-tests using split-half bootstrap NC values). Thus, other brain regions implicated in high-level cognition (MD, DMN) were not as reliable as the language regions in their responses to linguistic stimuli (and similarly not as well-predicted by GPT2-XL features; <u>SI</u> 8). In summary, the high NCs of the language regions show that these regions process stimulus-related information in a similar way across participants (see also ^75–78,81,82^), opening the door to investigations of what stimulus properties affect neural responses (see next section).

Second, we examined whether the five regions that comprise the LH language network are similar in their responses at the fine-grained level of single sentences. Prior work has demonstrated that the LH language regions exhibit a) similar functional response profiles in terms of their selectivity for language relative to non-linguistic inputs (e.g., ^13–15,17,18^) and similar sensitivity to diverse linguistic manipulations (e.g., ^10,21,58^), as well as b) highly correlated time courses during naturalistic paradigms (e.g., ^59,60,83,11^). Here, we investigated whether the 5 LH language regions have similar preferences for some sentences over others across n=1,000/n=1,500 sentences.

**Figure 4B** shows the Pearson correlation across the n=1,000 *baseline* sentences for LH fROIs from the average of n=5 *train* participants. Correspondingly, **Figure 4C** shows the correlation across the n=1,500 *drive*/*suppress*/*baseline* sentences for LH fROIs from the average of n=3 *evaluation* participants. Both plots show high inter-fROI correlations for the LH language network (correlation range 0.47-0.83), which suggests that even in their fine-grained preferences for particular linguistic stimuli, the LH language fROIs show a high degree of similarity. Along with the prior body of evidence noted above, these high correlations motivated our decision to investigate what kinds of linguistic input engage this network as a whole (see next section).

### Sentence complexity modulates language network responses

In order to gain understanding of what sentence properties modulate brain responses in the language network, we obtained a set of 11 features to characterize our experimental materials (n=2,000 sentences: 1,000 *baseline*, 250 *drive* and 250 *suppress* sentences from the *search* approach, and 250 *drive* and 250 *suppress* sentences from the exploratory *modify* approach, <u>SI 16</u>) and correlated these features with sentence-level brain responses (<u>Methods; Sentence</u> <u>properties that modulate brain responses</u>). The choice of features was inspired by past work in linguistics/psycholinguistics and cognitive neuroscience of language. First, building on prior evidence that surprisal (the degree of contextual predictability, which is typically estimated as negative log probability), modulates language processing difficulty in both behavioral psycholinguistic work (e.g., ^84–88^) and brain imaging investigations (e.g., ^89–97^), we computed sentence-level log probability estimates for each of 2,000 sentences using GPT2-XL (<u>Methods;</u> <u>Sentence properties that modulate brain responses</u>). And second, we collected 10 behavioral rating norms from a total of n=3,600 participants (on average, 15.23 participants per sentence per norm, min: 10, max: 19). The norms spanned five broad categories and were selected based on prior behavioral (e.g., ^98–100,85,101–103^) and neural studies (e.g., ^104–111,21^). The first category targeted two core aspects of sentences: grammatical well-formedness (how much does the sentence obey the rules of English grammar?; for details of the instructions, see <u>SI</u> <u>22C</u>) and plausibility (how much sense does the sentence make?). Because sentence surprisal (log probability), as estimated with GPT2-XL, is likely to capture both of these aspects to some extent (e.g., ^112–117^), we grouped these two norms with surprisal in the analyses. Furthermore, because more generally, surprisal likely captures diverse aspects of form and meaning, we examined the norm-brain relationships for all other norms after factoring out variance due to surprisal. Inspired by work on distributed neural representation of meaning, including across the language network (e.g., ^118–121^), the next three norms probed different aspects of the sentence content: how much does the sentence make you think about i) others’ mental states, ii) physical objects and their interactions, and iii) places and environments. The latter two have to do with the physical world, and the former — with internal representations; the physical vs. social distinction is one plausible organizing dimension of meaning ^122,123^. Two norms probed emotional dimensions of the sentences: valence (how positive is the sentence’s content?) and arousal (how exciting is the sentence’s content?). One norm targeted visual imagery (how visualizable is the sentence’s content?). Finally, the last two norms probed people’s perception of how common the sentence is, in general vs. in conversational contexts.

**Figure 5A** shows the correlation between the language network’s response and each of 11 sentence properties across the five categories. The sentences spanned a broad range of brain responses as evidenced in the sentence-level scatter plots in **Figure 5C** (y-axis). Importantly, this broad range was made possible by our approach of specifically designing stimuli to drive and suppress neural responses. Notice how the *drive* and *suppress* sentences cover parts of the linguistic space that are barely covered by the set of naturalistic *baseline* sentences (for comparisons of linguistic properties among conditions, see <u>SI 19</u>).

**Figure 5.**
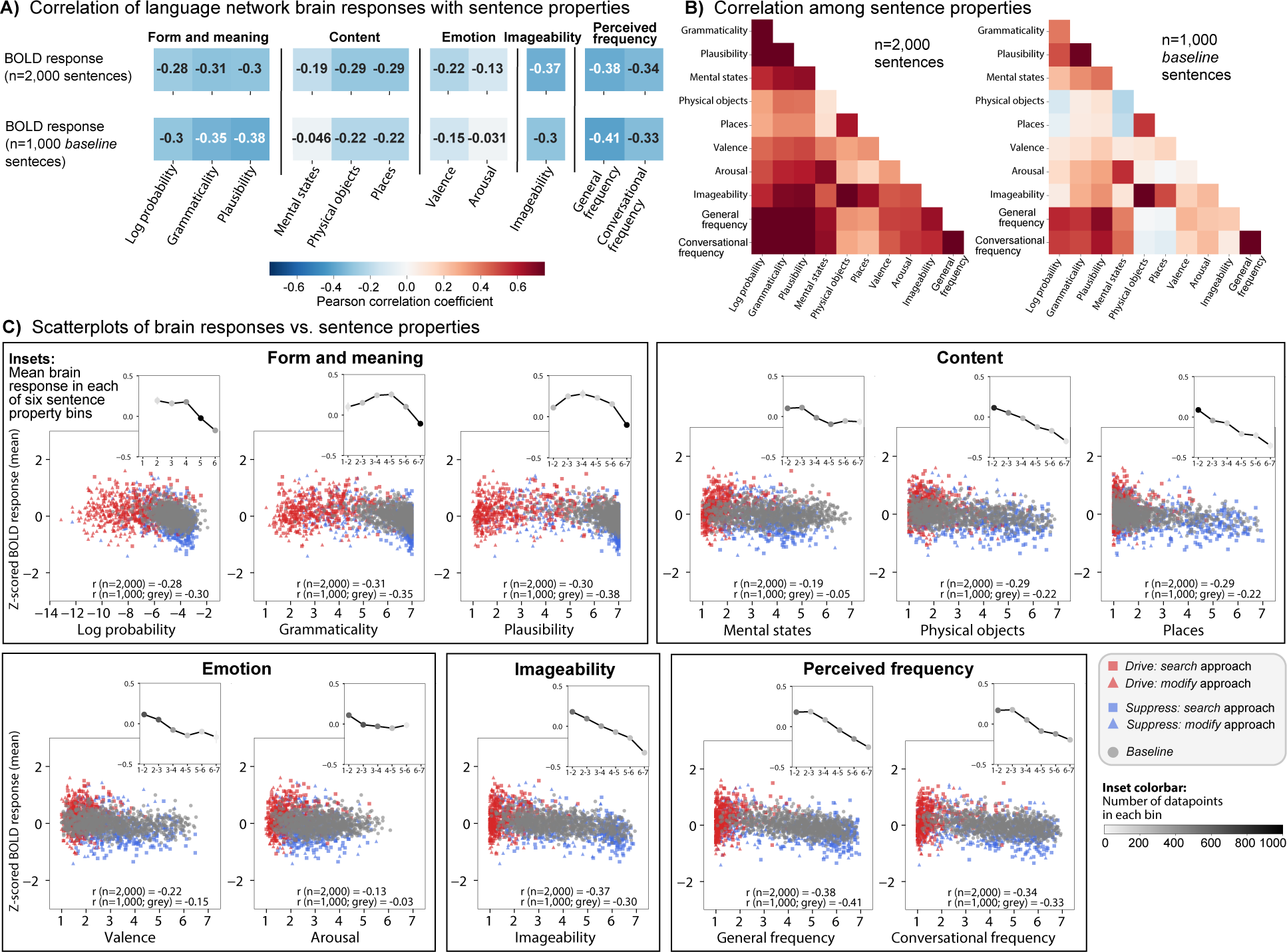
Surprisal and several other sentence properties modulate responses in the language network. **A)** Correlation of the LH language network’s response with 11 sentence properties (columns) within five categories for all n=2,000 sentences (first row; *drive, suppress,* and *baseline* sentences averaged across n=5 *train* and n=5 *evaluation* participants) and n=1,000 *baseline* sentences (second row; similarly averaged across n=5 *train* and n=5 *evaluation* participants). **B)** Correlation among the sentence properties shown for either n=2,000 sentences (left matrix) or n=1,000 sentences (right matrix). Color scale same as in A. **C)** Sentence-level brain responses as a function of sentence property. The brain responses (y-axis) were averaged across n=5 *train* and n=5 *evaluation* participants. The sentence properties were derived from behavioral norming experiments in independent participants (besides the “Log probability” feature which was obtained from GPT2-XL). The inset line graphs show the average brain response with each property grouped into six uniformly sized bins. Error bars show standard error of the mean across items in each bin (often not visible given the large number of data points). For the behavioral norms, the bins were defined according to the rating scale, i.e., {1,2}, {2,3}, {3,4}, {4,5}, {5,6}, and {6,7}. For log probability, the bins were similarly uniformly spaced, but according to the range of surprisal values: {-13.1,-11.3}, {-11.3,-9.4}, {-9.4,-7.5}, {-7.5,-5.7}, {-5.7,-3.8}, and {-3.8,-1.9} (omitted in the x-axis label). The color of the points in these graphs denotes the amount of data in each bin (darker dots correspond to larger amounts of data; bins containing less than 1% of the data, i.e., 20 responses, were omitted from the line graphs). Statistical comparisons accompanying the inset plots can be found in <u>SI 24</u>.

In terms of the effects of different sentence properties on neural responses, first, we found that less probable, i.e., more **surprising**, sentences elicited higher brain responses (**Figure 5C**) (r=-0.30 for the n=1,000 *baseline* sentences, dof=998, p<.001, t=-9.83, SE=0.03; see **Figure 5C** for the correlation values for the full set of n=2,000 sentences and <u>SI 19</u> for robustness to model choice to derive surprisal). This result aligns with previous evidence for a positive effect of surprisal on brain responses in MEG/EEG (e.g., ^89,95^) and fMRI (e.g., ^90–92,96^). Similarly, for the predictors related to a sentence’s **grammaticality and plausibility**, sentences that were rated as less grammatical or plausible elicited higher responses (r=-0.31, r=-0.30, dof=998, t=-11.92, t=-12.79, both p<.001, both SE=0.03; the two norms were correlated with each other at r=0.74). To understand whether grammaticality or plausibility explained variance above and beyond surprisal and each other, we fitted linear mixed effect models with different sets of sentence properties as predictors and compared these using likelihood ratio tests (see <u>Methods ;</u> <u>Statistical analyses</u> and <u>SI 23</u>). Plausibility explained variance beyond surprisal and grammaticality (*X*^2^=17.86, p<.001; all likelihood ratio statistics reported on the *baseline set*). Similarly, grammaticality explained variance beyond surprisal and plausibility (*X*^2^=12.97; p<.001), albeit to a lesser extent. Interestingly, a finer-grained examination of the relationship between these features and neural responses reveals a non-linearity, such that sentences in the mid-range of grammaticality and plausibility elicit stronger responses than sentences on the lower and higher ends of the scales (**Figure 5C**; note also that the response to surprisal asymptotes at the higher end of the surprisal scale such that more surprising sentences no longer lead to stronger responses). This pattern suggests that two effects may be at play: an increase in neural response is seen i) for sentences that better adhere to form and meaning regularities of language (similar to the previously reported stronger responses to sentences than lists of words; e.g., ^2,124^), and ii) for sentences that may have greater processing costs due to their unexpected form and/or meaning (e.g., see ^91,125^ for evidence of a strong relationship between behavioral processing difficulty and the strength of neural response in the language areas).

For the properties that relate to the sentence **content**, we evidenced no increase in explained variance (beyond surprisal) related to whether the sentence concerned others’ mental states (*X*^2^=0.69, p=0.407). This finding aligns with evidence that the language network does not support mental state inference and is robustly dissociated from the Theory of Mind network (e.g., ^60,78,126^) and challenges claims that the language areas are modulated by social content (e.g., ^127–129^). However, whether the sentence’s content concerned physical objects or places correlated negatively with brain responses (both r=-0.22, dof=998, both p<.001, t=-7.04 and t=- .11, both SE=0.03) and explained variance beyond surprisal (physical objects: *X*^2^=74.26, p<.001; places: *X*^2^=63.47, p<.001; the two norms were correlated with each other at r=0.54). Note, however, that these two aspects of the sentence content were also strongly correlated with imageability (discussed below), which may be the underlying driver of these effects.

For the properties that relate to the **emotional** aspects of sentences, we found that valence correlated negatively with brain responses, such that more positive sentences elicited a lower response (r=-0.15, dof=998, p<.001, t=-4.69, SE=0.03) and explained some variance beyond surprisal (*X*^2^=16.53, p<.001). In contrast, whether the sentence was exciting did not explain additional variance beyond surprisal (r=-0.03, dof=998, p=0.329, t=-0.98, SE=0.03; likelihood ratio *X*^2^=0.18, p=0.668).

**Imageability**–whether sentences are easy to visualize–was strongly correlated with whether the sentence’s content concerned physical objects (r=0.75) and places (r=0.49). Imageability strongly modulated brain responses, such that sentences rated as more imageable elicited a lower response (r=-0.30, dof=998, p<.001, t=-10.04, SE=0.03) and explained variance beyond surprisal (*X*^2^=93.03, p<.001).

Finally, for **perceived frequency**, we found that sentences that are perceived as more frequent (either in general or in conversational settings; these two norms were correlated with each other at r=0.77) elicited lower responses (r=-0.41 and r=-0.33, dof=998, both p<.001, t=-14.14 and t=- 10.89, both SE=0.03), with additional variance explained beyond surprisal (general perceived frequency: *X*^2^=96.63, p<.001; conversational perceived frequency: *X*^2^=44.46, p<.001).

To summarize the findings in this section, sentences that are surprising, fall in the middle of the grammaticality and plausibility range, and are perceived as not very frequent elicit a stronger response in the language network. In contrast, sentences that have positive content, talk about physical objects and places, and, more generally, are easy to visualize elicit a lower response in the language network (**Figure 5**). These patterns were highly similar across individual LH language fROIs and anatomically defined language ROIs, but showed some differences from the RH language network in line with some past claims (<u>SI 21)</u>.

## Discussion

We provide the first demonstration of non-invasive neural activity control in areas that are implicated in higher-level cognition: a brain-aligned Transformer model (GPT2-XL) can be used to drive and suppress brain responses in the language network of new individuals. We also provide a rich characterization of stimulus properties that modulate neural responses in the language network and find that less probable sentences generally elicit higher responses, with additional contribution from several form- and meaning-related features.

A number of studies have now shown that representations extracted from neural network models of language can capture neural responses to language in human brains, as recorded with fMRI or intracranial methods (e.g., ^130–132,41–47,133,50,49,51–53^). These studies have been conducted in an ‘open-loop’ manner: brain responses are simply acquired to a set of stimuli without any attempt to achieve specific levels of brain activity according to quantitative predictions. These stimulus sets have been limited to naturally occurring sentences, which cover a restricted portion of the space of linguistic/semantic variation. Further, the encoding model is typically trained and tested on data from the same participant (e.g., ^130–132,41–47,50,49,53,52^, cf. ^51,133^), potentially making it overly reliant on patterns of participant-specific idiosyncrasies. Thus, prior work has established similarity between LLM and humans on a narrow distribution of linguistic input and using within-participant evaluation in an open-loop fashion. In this work, we go beyond these studies by taking inspiration from closed-loop stimulus design in visual systems neuroscience ^134,54–57^: we evaluate the ability of an LLM-based encoding model to modulate the strength of neural responses in new individuals via new model-selected stimuli. Unlike typical encoding or representational similarity approaches to testing neural networks as models of the brain, we here utilize their predictive power to generate stimuli that would maximally drive or suppress responses for the language network. We emphasize that although using LLMs to identify new stimuli requires similarity to the human brain, this similarity need not hold at the implementation level, only at the level of representations. We, and others, acknowledge that the hardware of LLMs differs in many ways from human neural circuits (but see ^135^). These hardware differences, possibly coupled with factors such as training data and objective, could explain why LLMs sometimes diverge in from human-level performance for common linguistic phenomena such as negation and quantifier use (e.g., ^136,137^). Nevertheless, in spite of these differences, LLMs and the human language system appear to arrive at a similar representational space (see ^138^ for similar findings in vision), making LLMs currently the most predictive models of the human language network at the granularity of fMRI voxels and intracranial recordings (e.g., ^43,45^) and allowing us to modulate brain responses via targeted stimulus selection.

A priori, one might expect this approach to not be feasible within the domain of language because obtaining reliable neural responses to particular linguistic stimuli is challenging. First, unlike largely bottom-up brain systems such as the ventral visual stream ^139^, the language system extracts *abstract meaning representations* from linguistic sequences, which makes these representations further removed from the stimulus proper and thus more divergent across individuals, especially for more abstract meanings ^140^. And second, language processing requires attentional engagement ^141^, and such engagement is difficult to sustain for an extended period of time, especially if stimuli are repeated. One recent approach to combat fatigue/boredom has been to turn to rich naturalistic stimuli, like stories, podcasts, or movies and to collect massive amounts of data (sometimes, many hours’ worth) from a small number of individuals (e.g., ^118,142,143^)—what is often referred to as the ‘deep data’ approach (e.g., ^144–149^). However, such stimuli plausibly do not sample the space of linguistic and/or semantic variation well (see <u>SI 10</u> for evidence), and consequently, do not allow for testing models on stimuli that differ substantially from those used during training. We solved these methodological challenges by collecting neural responses to each of 1,000 semantically, syntactically, and stylistically diverse sentences for each participant in rapid, event-related fMRI, presented once to maximize engagement. We extended existing state-of-the-art methods for single-trial modeling ^150^ and reliability estimation (e.g., ^63^) to obtain robust neural responses to each sentence. Even with robust neural data, it was unclear whether encoding model performance is contingent on features that are specific to the stimulus set and/or participant at hand ^51^, which would limit generalization to i) stimuli that differ from the ones in the training set and/or ii) brain data from new individuals. By showing that model-selected stimuli successfully modulate brain responses in new individuals in ways predicted by the model, we established that LLM representations contain information that can be utilized for causal perturbation of language responses in the human brain in a general, participant-independent fashion.

We identified sentences that would push neural activity towards the edges of the stimulus-response distribution (driving and suppressing) using quantitative model-based predictions. Obtaining neural responses that span a wide range of activation levels enables us to ask which stimulus properties maximally (or minimally) engage the language network in the human brain, bringing us closer to understanding the representations and computations that support language comprehension. This general approach dates back to the pioneering work of Hubel and Wiesel ^151,152^ that provided an understanding of visual cortical computations by examining what stimuli cause each neuron to respond the most. Because linguistic input is extremely rich and language-responsive neuronal populations could, in principle, be tuned to many (possibly interacting) dimensions related to lexical, syntactic, semantic, or other linguistic properties, including ones that were not hypothesized in advance, we here identified target *drive* and *suppress* sentences using model predictions, thus removing experimenter bias.

Of course, a predictive model can be developed using features from *any quantitative representation* of sentences, including hidden states from an LLM (as we do here) but also much simpler univariate measures of different linguistic properties. Following a reviewer’s suggestion, we explicitly compared the predictivity performance of our encoding model, which uses GPT2-XL hidden states as features, to the performance of encoding models that use three univariate measures of surprisal (we focus on surprisal given its prominence in theorizing and empirical work on language ^89–97^). The encoding models based on univariate surprisal estimates perform substantially lower than the encoding model based on GPT2-XL hidden states (<u>SI 17</u>). Importantly, however, our motivation for using GPT2-XL representations goes beyond predictivity performance. LLMs allow for an *assumption-neutral and multi-faceted approach* for stimulus identification. Because LLMs are optimized for next-word prediction, their representations contain information about linguistic regularities at all levels, from word-level properties (including both word forms and their meanings), to syntactic structure, to semantic compositional meanings ^153–159^. This is because all of these properties can inform what word is likely to come next. By virtue of its assumption neutrality, this approach allows for *bottom-up discovery*. Surprisal models (e.g., based on n-grams or structure probabilities in a PCFG parser; SI 20) have the advantage of being interpretable, but can only be used for testing specific hypotheses. Neural network language models also allow for the testing of specific hypotheses, but additionally enable bottom-up discoveries of features that may not have been hypothesized in advance.

Indeed, we identified *drive* sentences that we could not have come up with in advance. These sentences were unusual on various dimensions related to their linguistic properties (<u>SI 19</u>) and highly distinct from the naturalistic *baseline* sentences (<u>SI 11;</u> note that the *suppress* sentences were more akin to naturalistic sentences), making these sentences a priori unlikely to be created or selected by experimenters and unlikely to be present in naturalistic stimuli, like stories or movies (<u>SI 10</u>). Yet these stimuli were able to drive responses in the language network.

To understand what stimulus properties modulate neural responses, we examined the effects of 11 sentence properties on the brain responses to the linguistically diverse set of 2,000 sentences. In line with much past work (e.g., ^89–97^), we found that surprisal has a strong effect on neural activity, with less probable sentences eliciting higher responses. However, a number of other properties explained variance beyond surprisal, including grammatical well-formedness and plausibility. Examining responses to a highly diverse set of sentences revealed a non-linearity in neural response in the form of an inverted-U shape. Sentences in the mid-range of well-formedness and plausibility elicit the highest response. This response is higher than a) the response to sentences in the low range, similar to the previously reported effects of stronger responses to phrases and sentences than lists of unconnected words (e.g., ^2,160,124^). The response is also higher than b) the response to sentences in the high range—sentences that are highly plausible and use common grammatical structures—which are easy to process (e.g., ^87^). Put differently, it appears that in order to elicit a strong response in the language network, a stimulus has to sufficiently resemble the kind of input we encounter in our experiences with language, given that our experiences presumably tune the language network to those kinds of stimuli ^161^. However, once some minimal level of language-likeness is reached, neural responses are modulated by processing difficulty, which depends on a combination of lexical, syntactic, and semantic features. Finally, one contribution of this work relative to past brain imaging studies is that we show sensitivity to these different linguistic properties at the fine-grained level of individual sentences (cf. standard blocked or event-related designs where groups of sentences are compared). In this way, we believe this rich dataset powerfully complements and extends prior evidence (e.g., ^104,105,91,90,96,125^) and allows for testing of new hypotheses about linguistic/semantic properties affecting neural responses.

A few limitations and future directions are worth noting. First, we here studied the language network—comprised of three frontal and two temporal areas—as a whole. As discussed earlier, there are good reasons to adopt this approach: the different regions of this network i) have similar functional response profiles, both with respect to their selectivity for language (e.g., ^13–15,17,18^) and their responses to linguistic manipulations (e.g., ^21,58^), and ii) exhibit highly correlated time courses during naturalistic cognition paradigms (e.g., ^59,60,83,11^). However, some functional heterogeneity has been argued to exist within the language network (e.g., ^162–164,160,165,166^). Future efforts using an approach like the one adopted here may discover functional differences within the language network (by searching for stimuli that would selectively drive particular regions within the network) as well as between the core LH language network and the RH homotopic areas and other language-responsive cortical, subcortical, and cerebellar areas. Second, the current results are limited to English but can be extended to other languages given the advances in multi-lingual language models (e.g., ^167^). Third, we have here relied on fMRI—a method with an inherently limited temporal resolution. Data from fMRI could be fruitfully supplemented with data from intracranial recordings, which would allow for model representations to be related to neural activity in a temporally resolved, word-by-word fashion and potentially uncover functional dissociations that are obscured when activity is averaged across adjacent words. Finally, novel ways of quantifying properties of linguistic input, e.g., based on the LLM representational space (e.g., ^157,158^), hold great potential to further understand how certain sentences modulate responses in the mind and brain.

In conclusion, we demonstrate modulation of brain responses in new individuals in the language network in a ‘closed-loop’ manner. This work has far-reaching implications for neuroscientific research and clinical applications. In particular, an accurate model-to-brain encoding model can serve as a quantitative, assumption-neutral tool for deriving experimental materials aimed at understanding the functional organization of the language network and putatively downstream areas that support abstract knowledge and reasoning (e.g., ^168,169,73,170^). Moreover, accurate encoding models can be used as a ‘virtual language network’ to simulate experimental contrasts in silico (e.g., ^171–173^). In particular, the model-selected sentences can be queried in a high-throughput manner to analyze the response properties of the language network in detail, providing the ability to rapidly generate novel hypotheses about language processing that can then be tested in a ‘closed-loop’ manner. For prospective clinical application, stimuli can be optimized for eliciting a strong response, thus allowing for efficient identification of language circuits, which may be especially important for individuals with brain disorders and other special populations, or in circumstances where time is of essence (e.g., neurosurgical planning and intraoperative testing). Finally, integrating the rapid advancements of artificial neural network models with larger and/or time-resolved measures of neural activity opens the door to even more fine-grained control of areas implicated in higher-level cognition.

## Methods

All experiments were performed with ethical approval from MIT’s Committee on the Use of Humans as Experimental Subjects (COUHES) (protocol number 2010000243). All participants gave informed written consent before starting the experiments.

We developed an encoding model to predict brain responses to arbitrary new sentences in the language network and evaluated this model by **i)** identifying novel sentences that are predicted to activate the language network to a maximal (or minimal) extent, and **ii)** collecting brain responses to these sentences in new participants. We then investigated which stimulus properties drive the responses in the language network (see **Figure 6** for an overview of the study).

**Figure 6.**
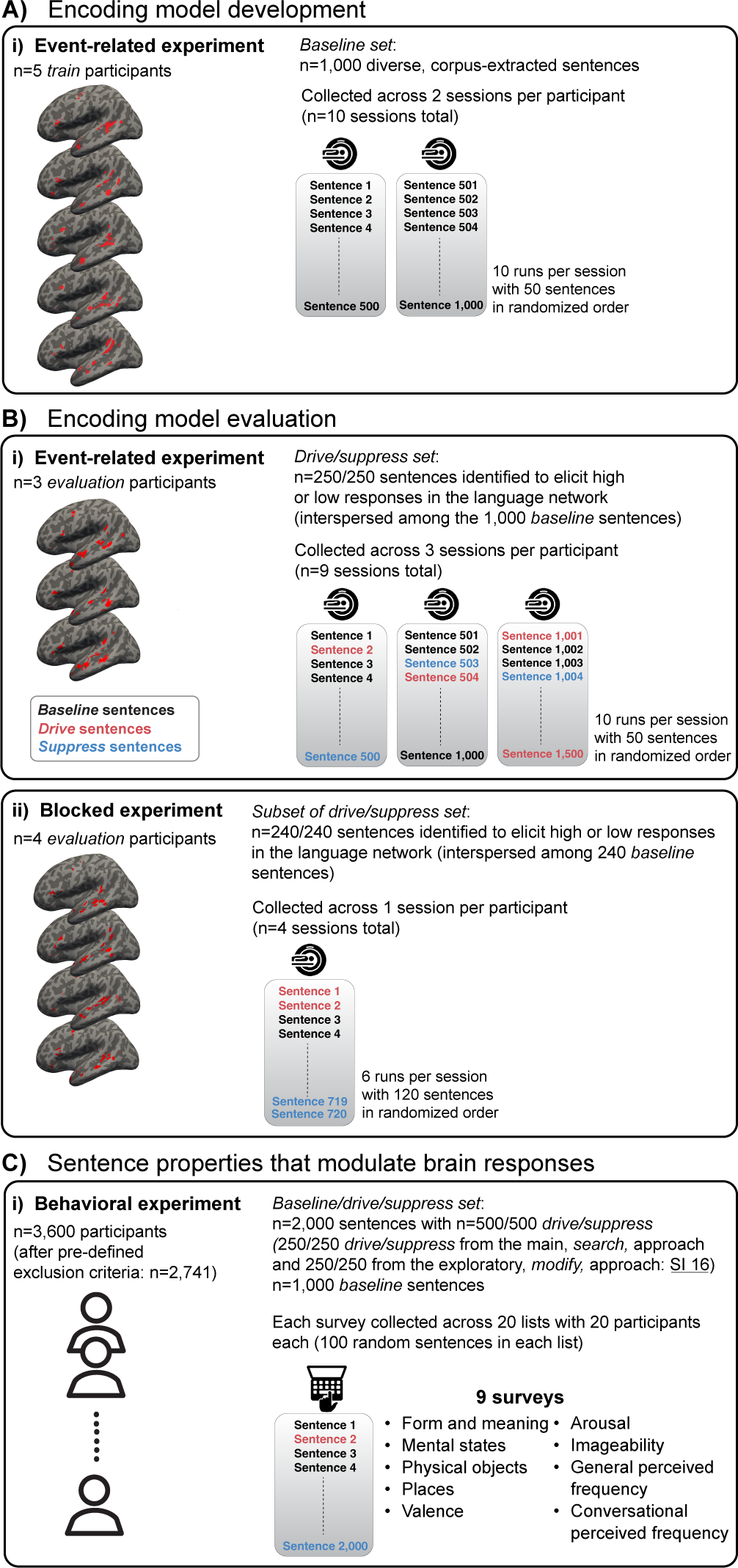
Experimental overview. **A) Encoding model development:** We curated a large set (n=1,000) of diverse, corpus-extracted sentences (*baseline set)* and collected brain responses in n=5 *train* participants in an event-related fMRI design across two sessions per participant. **B) Encoding model evaluation:** We identified a set of sentences to activate the language network to a maximal extent (250 *drive* sentences) or minimal extent (250 *suppress* sentences) by searching across ∼1.8 million sentences (*search* approach). We collected responses to these 500 *drive*/*suppress* sentences randomly interspersed among the *baseline* sentences in n=3 new participants (*evaluation* participants) across three sessions per participant in the event-related design (**panel i**). Moreover, we collected responses to a large subset of the *drive, suppress,* and *baseline* sentences (240 from each condition, a total of 720 sentences) in n=4 new participants in a blocked fMRI design within one session for each participant. **C) Sentence properties that modulate brain responses**: In order to understand what sentence properties modulate brain responses in the language network, we collected 10 behavioral rating norms (across 9 surveys) to characterize our experimental materials (n=2,000 sentences: 1,000 *baseline*, 250 *drive* and 250 *suppress* sentences from the *search* approach, and 250 *drive* and 250 *suppress* sentences from the exploratory *modify* approach; see <u>SI 16</u>) across n=3,600 participants.

### Encoding model development

#### General approach and data collection

We developed an encoding model of the left hemisphere (LH) language network in the human brain. Developing an encoding model requires brain responses to a broad range of linguistic input. Therefore, we curated a large set of diverse, corpus-extracted 6-word sentences (n=1,000, *baseline set*), collected brain responses while five participants (*train* participants) read each sentence in an event-related, condition-rich fMRI paradigm (each sentence equals a condition), across two sessions each, and modeled those responses using a recently developed single-trial modeling framework ^150^, which we adapted for no-repeats designs (<u>Methods; fMRI</u> <u>experiments</u> and <u>SI 3)</u>. The *baseline set* consisted of two subsets: the first subset (n=534 sentences) aimed to maximize semantic diversity to cover a broad range of topics, and the second subset (n=466 sentences) was selected from across diverse genres and styles (newspaper text, web media, transcribed spoken language, etc.) (<u>SI 1</u>). In five *train* participants, we recorded brain responses to the sentences in the *baseline set* across two scanning sessions (**Figure 6A**). Participants were instructed to read attentively and think about the sentence’s meaning. To encourage engagement with the stimuli, prior to the session, participants were informed that they would be asked to perform a short memory task after the session (<u>Methods;</u> <u>fMRI experiments</u>). Sentences were presented one at a time for 2 seconds with a 4 second inter-stimulus interval. Each run contained 50 sentences (5:36 minutes) and sentence order was randomized across participants.

The language network was defined functionally in each participant using an extensively validated localizer task (e.g., ^2,3^; <u>Methods; Definition of ROIs</u>). Although the network consists of five areas (two in the temporal lobe and three in the frontal lobe), we treat it here as a functionally integrated system given i) the similarity among the five regions in their functional response profiles across dozens of experiments (e.g., ^21,58,96^; see Figure 4B,C and <u>SI 4</u> for evidence of similar preferences for the *baseline set* in the current data), ii) high inter-regional correlations during naturalistic cognition paradigms (e.g., ^81,76,59,60,83,11^). To mitigate the effect of collecting data across multiple scanning sessions and to equalize response units across voxels and participants, the blood-oxygen-level-dependent (BOLD) responses were z-scored session-wise per voxel. BOLD responses from the voxels in the LH language network were averaged within each *train* participant (<u>Methods; Definition of ROIs</u>) and averaged across participants to yield an average language network response to each of the 1,000 *baseline set* sentences.

#### Encoding model

To develop an encoding model of the language network, we fitted a linear model from the representations of a large language model (LLM) to brain responses (an encoding approach; ^174^). The brain data that were used to fit the encoding model were the averaged LH language network’s response from the n=5 *train* participants. To map from LLM representations to brain responses, we made use of a linear mapping model. Note that the term “mapping model” refers to the regression model from LLM representations to brain activity, while the term “encoding model” encompasses both the LLM used to transform a sentence into an embedding representation as well as the mapping model.

The mapping model was a L2-regularized (“ridge”) regression model which can be seen as placing a zero-mean Gaussian prior on the regression coefficients ^175^. Introducing the L2-penalty on the weights results in a closed-form solution to the regression problem, which is similar to the ordinary least-squares regression equation:

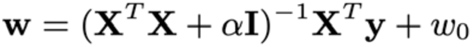

Where *X* is a matrix of regressors (*n* stimuli by *d* regressors). The regressors are unit activations from the sentence representations derived by exposing an LLM to the same stimuli as the human participant was exposed to and hence *d* refers to the number of units in the LLM embedding representation (“hidden size”). *y* is an n-length column vector containing the relevant brain ROI’s mean response to each stimulus. *I* is the identity matrix (d by d). *w* is a d-length column vector with the weights learned for each regressor. *w*_0_ is the intercept term. Alpha (*⍺*) is the regularization parameter that trades off between the fit to the data and the penalty for weights with high coefficients. To select this regularization parameter, we used leave-one-out cross-validation implemented using the *scikit-learn* Python library function *RidgeCV* (^176^; version 0.24.2). Specifically, for each of 60 logarithmically-spaced *⍺* regularization parameter values (1e-30, 1e-29, …, 1e28, 1e29), we measured the squared error in the resulting prediction of the left-out stimulus using regression weights derived from the other stimuli in the data. We computed the average of this error (across the stimuli) for each of the 60 potential *⍺* regularization parameter values. We then selected the *⍺* regularization parameter that minimized this mean squared error (*⍺* = 10,000). When cross-validation was performed, the *⍺* regularization parameter was always selected using the stimuli in the train split, and with the *⍺* parameter selected, the regression model using that parameter was used on the test split.

##### Encoding model performance

We obtained an unbiased estimate of encoding model performance using three different approaches: i) Cross-validated predictivity performance on held-out sentences (SI 6), ii) Cross-validated predictivity performance on held-out participants within the *train* participants (SI 7), and iii) held-out prediction performance on new participants (*evaluation* participants) *and* sentences (Methods; Encoding model evaluation and Results; Model captures most explainable variance in new participants).

All modeling and analysis code was written in Python (version 3.8.11), making heavy use of the *numpy* (^177^; version 1.21.2), *scipy* (^178^; version 1.7.3), *scikit-learn* (^176^; version 0.24.2), *pandas* (^179^; version 1.4.2) and *transformers* (^180^; version 4.11.3) libraries.

#### Sentence representations from large language models (LLMs)

To obtain sentence representations for the encoding model, we used the unidirectional-attention Transformer LLM GPT2-XL ^36^, which was identified as the most brain-aligned language base model in prior work ^43^ and which was the largest unidirectional OpenAI GPT model available on HuggingFace ^180^ at the time of the experiments (summer 2021)). (Supplementary analyses were performed using BERT-large (<u>SI 6C</u>).) We used the pretrained model available via the HuggingFace library (^180^, *transformers* version 4.11.3; https://huggingface.co/gpt2-xl). GPT2-XL has 48 layers (i.e., Transformer blocks) in addition to the embedding layer. The embedding dimension is 1,600. We obtained model representations by tokenizing each sentence using the model’s standard tokenizer (GPT2TokenizerFast) and passing each sentence through the model. We retrieved model representations for each model layer (i.e., at the end of each Transformer block). Given that human participants were exposed to the whole sentence at once, we similarly computed a sequence summary representation for each sentence. We obtained the representation of the last sentence token, given that unidirectional models aggregate representations of the preceding context (i.e., earlier tokens in the sentence). Further, to ensure that the results were robust to this choice of summary representation, we also obtained a sequence summary representation by computing the arithmetic mean of the representations associated with each token in each sentence (<u>SI 6B</u>). The resulting features were used as regressors in the LLM-brain comparisons. Each LLM layer (model stage for which representations were extracted, i.e., Transformer blocks) was treated as a separate set of regressors in LLM-brain comparisons. Layer 22 features were selected as regressors in the encoding model based on cross-validated model performance evaluation (<u>SI 6A</u>).

### Encoding model evaluation

#### General approach and data collection

Using our trained encoding model, we identified a set of novel sentences to activate the language network to a maximal extent (*drive* sentences) or minimal extent (*suppress* sentences). To do so, we searched across ∼1.8 million sentences to identify sentences predicted to elicit high or low fMRI responses (250 sentences of each kind) (**Figure 6B**; <u>SI 9</u>). We collected brain responses to these novel sentences in three new participants (across three sessions each). The *drive* and *suppress* sentences were randomly interspersed among the 1,000 *baseline* sentences (for a total of n=1,500 sentences), collected across three scanning sessions per participant (n=9 sessions total). (In a more exploratory component of the study, we complemented the *search* approach with another approach—the *modify* approach—where we used gradient-based modifications to transform a random sentence into a novel sentence/string predicted to elicit high or low fMRI responses. We collected brain responses to these novel sentences in two new participants (see <u>SI 16</u> for the details of methods and the results).)

To ensure that the results are robust and generalizable to different experimental paradigms, we additionally collected fMRI responses to a large subset of the *drive* and *suppress* sentences along with the *baseline* sentences in a traditional blocked design in four independent participants (one scanning session each). The participants for the blocked experiment were exposed to a total of 720 unique sentences (from the *baseline*, *drive*, *suppress* conditions; 240 per condition which were randomly sampled for each participant). Sentences were grouped into blocks of 5 sentences from the same condition and were presented on the screen one at a time for 2s with a 400ms inter-stimulus interval. Each run contained 120 sentences in 24 blocks (5:36 minutes). Condition order was counterbalanced across runs and participants (<u>Methods; fMRI</u> <u>experiments</u>).

### fMRI experiments

#### Participants

A total of 14 neurotypical adults (9 female), aged 21 to 31 (mean 25.3; std 3), participated for payment between October 2021 and December 2022. The sample size was based on those used for previous fMRI semantic decoding experiments ^118,119^. All participants had normal or corrected-to-normal vision, and no history of neurological, developmental, or language impairments. 12 participants (∼86%) were right-handed, as determined by self-report and the Edinburgh handedness inventory ^181^ and 2 (∼14%) were left-handed. All participants had a left-lateralized/bilateral language network as determined by the examination of the activation maps for the language localizer ^2^. All participants were native speakers of English. Each scanning session lasted between 1 and 2 hours. All participants gave informed written consent in accordance with the requirements of the MIT’s Committee on the Use of Humans as Experimental Subjects (COUHES) (protocol number 2010000243). Participants were compensated for their time ($30/hour).To err on the conservative side, no participants were excluded from the study based on data quality considerations.

#### Critical fMRI tasks

##### Sentence-reading task: Event-related design

We developed a paradigm to collect brain responses to as many individual sentences as possible (similar to recent paradigms in visual neuroscience, e.g., the Natural Scenes Dataset ^63^). Participants passively read each sentence once, in a condition-rich, event-related fMRI design (each sentence is effectively a condition). Sentences were presented (in black font) on a light grey background one at a time for 2s with a 4s inter-stimulus interval (ISI) consisting of a fixation cross. Each run contained 50 unique sentence trials and three 12s fixation blocks (in the beginning, middle (i.e., after 25 sentences) and end of each run). Each run lasted 336s (5:36 minutes).

Participants were instructed to read attentively and think about the sentence’s meaning. To encourage engagement with the stimuli, prior to the session, participants were informed that they would be asked to perform a short memory task after the session (outside of the scanner). The first five participants *(train* participants) were exposed to the set of n=1,000 *baseline* sentences and therefore completed 20 experimental runs (across two scanning sessions). The sentences were randomly assigned to runs for each participant (i.e., participants were exposed to different presentation orders).

The next five participants (*evaluation* participants) were exposed to n=250 *drive* and n=250 *suppress* sentences interspersed among the set of n=1,000 *baseline* sentences–a total of n=1,500 sentences–and therefore completed 30 runs of the experiment (across three scanning sessions). The n=1,500 sentences were randomly assigned to experimental runs for each participant while ensuring a balanced distribution of *baseline*, *drive*, and *suppress* sentences in each run, leading the following distribution of *baseline/drive/suppress* sentences in the three scanning sessions: 333/84/83, 333/83/84, and 334/83/83.

##### Sentence-reading task: Blocked design

To evaluate the robustness of brain responses to the *drive* and *suppress* sentences, we additionally presented a subset of the *drive*, *suppress*, and *baseline* sentence materials in a traditional blocked design.

Sentences were grouped into blocks of 5 sentences from the same condition (*baseline*, *drive*, *suppress*) and were presented on the screen (in black font on a light grey background) one at a time for 2s with a 400ms ISI consisting of a fixation cross (for a total block duration of 12s). Each run consisted of 24 blocks with 8 blocks (40 sentences) per condition. There were five 12s fixation blocks: in the beginning and end of each run, as well as after 6, 12, and 18 blocks. Each run lasted 348s (5:48 minutes).

As in the event-related experiment, participants were instructed to read attentively and think about the sentence’s meaning. Prior to the session, participants were informed that they would be asked to perform a short memory task after the session (outside of the scanner).

The participants for the blocked experiment were exposed to a total of 720 unique sentences (from the *baseline*, *drive*, *suppress* conditions; 240 per condition) across 6 runs in a single scanning session. These sentences were sampled randomly without replacement from the full set of materials (consisting of 250 *drive*, 250 *suppress*, and 1,000 *baseline* stimuli from the *search* approach). The sentences were randomly sampled and assigned to runs for each participant (i.e., participants were exposed to different presentation orders of different subsets of the materials). Condition order was counterbalanced across runs and participants.

##### Memory task for the sentence-reading task

For both the event-related and blocked critical sentence-reading experiments, participants completed a memory task at the end of each scanning session (outside of the scanner) to incentivize attention throughout the session.

Participants were informed ahead of time that they would be asked to perform a memory task after the scanning session. Participants were presented with a set of sentences, one at a time, and asked to decide for whether they had read it during the scanning session. For both the event-related and blocked experiment, the memory task consisted of 30 sentences: 20 sentences from the set used in the scanning session and 10 foil sentences. For the event-related experiment, the 20 correct targets were randomly sampled without replacement from each of the 10 runs in that session, 2 from each run. For the blocked experiment, the 20 correct targets were randomly sampled without replacement from each of the 6 runs in that session, 3 from each run, with an additional 2 sentences from random runs.

The 10 foil sentences were randomly sampled without replacement from a set of 100 sentences. These 100 foil sentences were manually selected from the same corpora that were used to construct the baseline stimulus set (15 sentences from each of the three genres from The Toronto Book Corpus–45 in total–and 55 sentences from the additional eight corpora, see <u>SI 1</u>). The average accuracy (sum of correct responses divided by total number of responses; chance-level is 50%) was 70.4% (SD across sessions: 11.4%) for the event-related participants (n=24 sessions – responses for one session were not saved due to an error in the script), and 61.7% (SD across sessions: 10%) for the blocked participants (n=4 sessions).

#### fMRI data acquisition, preprocessing and first-level analysis

##### fMRI data acquisition

Structural and functional data were collected on the whole-body, 3 Tesla Siemens Prisma scanner with 32-channel head coil, at the Athinoula A. Martinos Imaging Center at the McGovern Institute for Brain Research at MIT. T1-weighted, Magnetization Prepared RApid Gradient Echo (MP-RAGE) structural images were collected in 176 sagittal slices with 1 mm isotropic voxels (TR = 2,530 ms, TE = 3.48 ms, TI = 1100 ms, flip = 8 degrees). Functional, blood oxygenation level dependent (BOLD), data were acquired using an SMS EPI sequence (with a 90 degree flip angle and using a slice acceleration factor of 2), with the following acquisition parameters: fifty-two 2 mm thick near-axial slices acquired in the interleaved order (with 10% distance factor) 2 mm × 2 mm in-plane resolution, FoV in the phase encoding (A ≫ P) direction 208 mm and matrix size 104 × 104, TR = 2,000 ms and TE = 30 ms, and partial Fourier of 7/8. The first 10 s of each run were excluded to allow for steady state magnetization.

##### fMRI preprocessing

fMRI data were preprocessed using SPM12 (release 7487), CONN EvLab module (release 19b), and custom MATLAB scripts. Each participant’s functional and structural data were converted from DICOM to NIfTI format. All functional scans were coregistered and resampled using B-spline interpolation to the first scan of the first session. Potential outlier scans were identified from the resulting subject-motion estimates as well as from BOLD signal indicators using default thresholds in CONN preprocessing pipeline (5 standard deviations above the mean in global BOLD signal change, or framewise displacement values above 0.9 mm; ^182^. Note that the identification of outlier scans was leveraged in the blocked first-level modeling, but not in the data-driven event-related first-level modeling). Functional and structural data were independently normalized into a common space (the Montreal Neurological Institute [MNI] template; IXI549Space) using SPM12 unified segmentation and normalization procedure ^183^ with a reference functional image computed as the mean functional data after realignment across all timepoints omitting outlier scans. The output data were resampled to a common bounding box between MNI-space coordinates (−90, −126, −72) and (90, 90, 108), using 2 mm isotropic voxels and 4^th^ order spline interpolation for the functional data, and 1 mm isotropic voxels and trilinear in^te^rpolation for the structural data. Last, the functional data were smoothed spatially using spatial convolution with a 4 mm FWHM Gaussian kernel.

##### First-level modeling of event-related experiments

The critical, event-related experiment was analyzed using GLMsingle ^150^, a framework for obtaining accurate response estimates in quick event-related single-trial fMRI designs. Modeling such responses is challenging due to temporal signal autocorrelation, participant head motion, and scanner instabilities. The GLMsingle framework introduces three main steps to combat noise in a data-driven manner: 1) Choice of HRF to convolve with the design matrix: an HRF is identified from a library of 20 candidate functions (derived from independent fMRI data ^63^) as the best fitting for each voxel separately, 2) Noise regressors: a set of voxels that are unrelated to the experimental paradigm are identified and these voxels’ time courses are used to derive an optimal set of noise regressors by performing principal component analysis (PCA), and 3) Regularization of voxel responses: instead of an ordinary least squares (OLS) regression, GLMsingle uses fractional ridge regression ^184^ to model voxel responses in order to dampen the noise inflation in a standard OLS regression due to correlated predictors from rapid, successive trials.

Using this framework, a General Linear Model (GLM) was used to estimate the beta weights that represent the blood oxygenation level dependent (BOLD) response amplitude evoked by each individual sentence trial (fixation was modeled implicitly, such that all timepoints that did not correspond to one of the conditions (sentences) were assumed to correspond to a fixation period). Data from different scanning sessions for a given participant were analyzed together. The ‘sessionindicator’ option in GLMsingle was used to specify how different input runs were grouped into sessions. For each voxel, the HRF which provided the best fit to the data was identified (based on the amount of variance explained). The data were modeled using a fixed number of noise regressors (5) and a fixed ridge regression fraction (0.05) (these parameters were determined empirically using an extensive joint data modeling and data evaluation framework, see <u>SI 3</u>).

By default, GLMsingle returns beta weights in units of percent signal change by dividing by the mean signal intensity observed at each voxel and multiplying by 100. Hence, the beta weight for each voxel can be interpreted as a change in BOLD signal for a given sentence trial relative to the fixation baseline. To mitigate the effect of collecting data across multiple scanning sessions, the betas were z-scored session-wise per voxel (see <u>Methods; Definition of ROIs</u>).

##### First-level modeling of blocked experiments

Blocked experiments were analyzed using standard analysis procedures using SPM12 (release 7487), CONN EvLab module (release 19b). Effects were estimated using a GLM in which the beta weights associated with each experimental condition was modeled with a boxcar function convolved with the canonical HRF) (fixation was modeled implicitly, such that all timepoints that did not correspond to one of the conditions were assumed to correspond to a fixation period). Temporal autocorrelations in the BOLD signal timeseries were accounted for by a combination of high-pass filtering with a 128 seconds cutoff, and whitening using an AR(0.2) model (first-order autoregressive model linearized around the coefficient a = 0.2) to approximate the observed covariance of the functional data in the context of Restricted Maximum Likelihood estimation (ReML). In addition to experimental condition effects, the GLM design included first-order temporal derivatives for each condition (included to model variability in the HRF delays), as well as nuisance regressors to control for the effect of slow linear drifts, subject-motion parameters, and potential outlier scans on the BOLD signal.

### Definition of regions of interest (ROIs)

#### Language regions of interest (ROIs)

Language regions of interest (ROIs) were defined in individual participants using functional localization (e.g., ^79,2,80,185^). This approach is crucial because many functional regions do not exhibit a consistent mapping onto macro-anatomical landmarks (e.g., ^186–188^) and this variability is problematic when functionally distinct regions lie in close proximity to each other, as is the case with both frontal and temporal language areas (e.g., ^189,83^; see Fedorenko and Blank^18^ for discussion of this issue for ‘Broca’s area’).

For each participant, functional ROIs (fROIs) were defined by combining two sources of information ^2,190^: i) the participant’s activation map for the localizer contrast of interest (t-map), and ii) group-level constraints (“parcels”) that delineated the expected gross locations of activations for the relevant contrast and were sufficiently large to encompass the extent of variability in the locations of individual activations (all parcels are available for download from https://evlab.mit.edu/funcloc/download-parcels).

##### Language network localizer task

The task used to localize the language network was a reading task contrasting sentences (e.g., THE SPEECH THAT THE POLITICIAN PREPARED WAS TOO LONG FOR THE MEETING) and lists of unconnected, pronounceable nonwords (e.g., LAS TUPING CUSARISTS FICK PRELL PRONT CRE POME VILLPA OLP WORNETIST CHO) in a standard blocked design with a counterbalanced condition order across runs (introduced in Fedorenko et al. ^2^)). The sentences > nonwords contrast targets higher-level aspects of language, including lexical and phrasal semantics, morphosyntax, and sentence-level pragmatic processing, to the exclusion of perceptual (speech- or reading-related) processes. The areas identified by this contrast are strongly selective for language relative to diverse non-linguistic tasks (e.g., ^14^; see Fedorenko and Blank ^18^ for a review). This paradigm has been extensively validated and shown to be robust to variation in the materials, modality of presentation, language, and task (e.g., ^2,7,11^, *inter alia*). Further, a network that corresponds closely to the localizer contrast emerges robustly from whole-brain task-free data—voxel fluctuations during rest ^83^.

Each stimulus consisted of 12 words/nonwords. Stimuli were presented in the center of the screen, one word/nonword at a time, at the rate of 450ms per word/nonword. Each stimulus was preceded by a 100ms blank screen and followed by a 400ms screen showing a picture of a finger pressing a button, and a blank screen for another 100ms, for a total trial duration of 6s. Experimental blocks lasted 18s (with 3 trials per block), and fixation blocks lasted 14s. Each run (consisting of 5 fixation blocks and 16 experimental blocks) lasted 358s. Participants completed 2 runs. Participants were instructed to read attentively (silently) and press a button on the button box whenever they saw the picture of a finger pressing a button on the screen. The button-pressing task was included to help participants remain alert.

The materials and scripts are available from the Fedorenko Lab website (https://evlab.mit.edu/funcloc).

##### Language network fROIs

The language fROIs were defined using the *sentences > nonwords* contrast from the language localizer collected in each participant’s first scanning session (see e.g., Mahowald and Fedorenko ^191^, for evidence that localizer maps are highly stable within individuals over time, including across sessions). This contrast targets higher-level aspects of language, to the exclusion of perceptual (speech/reading) and motor-articulatory processes (for discussion, see Fedorenko and Thompson-Schill ^192^).

To define the language fROIs, each individual *sentences > nonwords* t-map was intersected with a set of ten binary parcels (five in each hemisphere). These parcels were derived from a probabilistic activation overlap map using watershed parcellation, as described by Fedorenko et al. ^2^ for the *sentences > nonwords* contrast in 220 independent participants and covered extensive portions of the lateral frontal, temporal, and parietal cortices. Specifically, five language fROIs were defined in the dominant hemisphere: three on the lateral surface of the frontal cortex (in the inferior frontal gyrus, *IFG*, and its orbital part, *IFGorb*, as well as in the middle frontal gyrus, *MFG*), and two on the lateral surface of the temporal and parietal cortex (in the anterior temporal cortex, *AntTemp*, and posterior temporal cortex, *PostTemp*). Following prior work (e.g., ^59^), to define the RH fROIs, the LH language parcels were transposed onto the RH, allowing the LH and RH homotopic fROIs to differ in their precise locations within the parcels.

Within each of these ten parcels, the 10% of voxels with the highest t-values for the *sentences > nonwords* contrast were selected (see SI 15E for number of voxels in each fROI).

#### Control regions of interest (ROIs)

In addition to language regions, we examined i) two large-scale brain networks linked to high-level cognitive processing—the multiple demand (MD) network ^64–68^ and the default mode network (DMN) ^69–73^ which—similar to the language regions—were functionally defined using independent localizer tasks in each participant, and ii) a set of anatomical parcels ^74^ in an effort to cover the entire cortex (see SI 15 for details).

#### Aggregation of voxels within each regions of interest (ROI)

The voxels belonging to each functional ROI (language, MD, and DMN) and each anatomical Glasser ROI were aggregated by averaging. For the fMRI data reported in the main text, each voxel was z-scored session-wise prior to averaging, in order to minimize potential non-stationarities that exist across different scanning sessions and to equalize response units across voxels. In <u>SI 12</u> and <u>SI 14</u>, we report fMRI data without any normalization (the key patterns of results are not affected).

On average, we extracted responses from 10 language fROIs (SD=0), 19.43 MD fROIs (SD=1.28), 12 DMN fROIs (SD=0), and 353.71 anatomical Glasser parcels (SD=10.34) across n=14 participants (5 *train* participants, 5 *evaluation* participants in the event-related fMRI design from the *search* and *modify* approaches, and 4 *evaluation* participants in the blocked fMRI design). In a few cases, (f)ROIs could not be extracted due to a negative t-statistic for the contrast of interest or lack of coverage in our functional acquisition sequence.

### Sentence properties that modulate brain responses

#### General approach

Finally, to shed light on what property or properties make some sentences elicit stronger responses in the language network, we collected an extensive set of norms to characterize the full set of sentences in this study (n=2,000: 1,000 *baseline* sentences, 250 *drive* and 250 *suppress* sentences from the *search* approach, and 250 drive and 250 suppress sentences from the exploratory *modify* approach) (**Figure 6C**) and examined the relationship between these properties and fMRI responses. First, building on the body of evidence for surprisal modulating language processing difficulty, in both behavioral psycholinguistic work (e.g., ^84–88^) and brain imaging investigations (e.g., ^89–97^), we computed the average log probability for each sentence using GPT2-XL (surprisal is negative log probability; see <u>Surprisal features</u>). And second, we collected 10 behavioral rating norms across a total of n=3,600 participants (on average, 15.23 participants per sentence per rating norm, min: 10, max: 19). The norms spanned five broad categories and were all motivated by prior work in linguistics and psycholinguistics (<u>see</u> Behavioral norms).

#### Surprisal features

We estimated the log probability of a word given its context for the words in each sentence. The negative log probability of a word/sentence is known as “surprisal” ^193,194^. The log probability of each sentence was computed using the pre-trained unidirectional-attention language model GPT2-XL ^36^ from the HuggingFace library (^180^, *transformers* version 4.11.3). GPT2-XL was trained on 40GB on web text from various domains (WebText dataset). Each sentence was tokenized using the model’s standard tokenizer (GPT2Tokenizer) and the special token, [EOS], was prepended to each sentence. Punctuation was retained. We obtained the sentence-level surprisal by taking the mean of the token-level surprisals.

For supplementary analyses, we obtained surprisal estimates from an n-gram model and a probabilistic context-free grammar model in addition to GPT2-XL (SI 20).

#### Behavioral norms

##### Participants

Participants were recruited using crowd-sourcing platforms: Prolific (n=8 surveys) and Amazon Mechanical Turk (mTurk; n=1 survey). For Prolific, the study was restricted to workers with English as their first language and their most fluent language, USA as their location, and a submission approval rate greater than or equal to 90%. For mTurk, the study was restricted to “Mechanical Turk Masters” workers. Across the 9 surveys, a total of 3,600 participants took part in the experiment (400 participants for each survey; see <u>SI 21C</u> for details). 2,741 participants remained after pre-defined exclusion criteria (<u>SI 21A</u>). The experiments were conducted with approval from and in accordance with MIT’s Committee on the Use of Humans as Experimental Subjects (COUHES) (protocol number 2010000243). Participants gave informed consent before starting each experiment and were compensated for their time (minimum $12/hour).

##### Materials, design, and procedure

The n=2,000 sentences were randomly assigned to 20 unique sets containing 100 sentences each. For each survey, the participants first provided informed consent. Then they answered several demographic questions (whether English is their first language, which country they are from, and what age bracket they fall into); they were explicitly told that payment is not contingent on their answers to these questions. Finally, they were presented with the survey-specific instructions and the following warning: “*There are some sentences for which we expect everyone to answer in a particular way. If you do not speak English or do not understand the instructions, please do not do this hit – you will not get paid.*”. One survey targeted two core aspects of sentences: grammatical well-formedness (how much does the sentence obey the rules of English grammar?; for details of the instructions, see <u>SI</u> <u>22C</u>) and plausibility (how much sense does the sentence make?). Three surveys probed different aspects of the sentence content: how much does the sentence make you think about i) others’ mental states, ii) physical objects and their interactions, and iii) places and environments. The latter two have to do with the physical world, and the former — with internal representations; the physical vs. social distinction is one plausible organizing dimension of meaning ^122,123^. Two surveys probed emotional dimensions of the sentences: valence (how positive is the sentence’s content?) and arousal (how exciting is the sentence’s content?). One survey targeted visual imagery (how visualizable is the sentence’s content?). Finally, the last two surveys probed people’s perception of how common the sentence is, in general vs. in conversational contexts. The first survey (with two questions per sentence) took 25.01 minutes on average; the remaining surveys took 14.25 min on average. After the participants answered the rating question(s) for the 100 sentences (the order was randomized separately for each participant), they were asked to complete 6 sentence preambles (e.g., “*When I was younger, I would often …*”; see <u>SI 22B</u> for the full set), which were used post-hoc to evaluate English proficiency. See <u>SI 22</u> for details on experimental procedures.

### Statistical analyses

Linear mixed effects (LME) models (implemented using the *lmer* function from the *lme4* R package ^195^; version 1.1-31) were used to evaluate the statistical significance i) of the differences in the BOLD response among the sentence conditions (*baseline, drive,* and *suppress*) and ii) of the effect of sentence properties on the BOLD response. The critical variable of interest (either condition or sentence property) was modeled as fixed effect(s). As additional effects, we modeled other variables that could modulate the BOLD response but that were not our critical variables of interest, including item (sentence), run order within a session (1-10), and sentence order within a run (1-50):

*BOLD response ∼ variable_of_interest + (1 | sentence) + run_within_session + trial_within_run.* (Note that because the BOLD responses were z-scored session-wise, there was no additional variance to explain by including session number or participant as a model term).

The models were fitted using maximum likelihood estimation and used the Satterthwaite method for estimating degrees of freedom. For each LME model reported, we provide (in <u>SI 18</u> and <u>SI</u> <u>23</u>) a table with model formulae, effect size estimates, standard error estimates, t-statistics, p-values, degrees of freedom, and R2 values. We evaluated the statistical significance of differences between pairs of conditions using estimated marginal means (implemented using the *emmeans* function from the *emmeans* R package ^196^; version 1.8.4-1) using Tukey’s multiple comparison method. Finally, we evaluated the statistical significance of differences between pairs of LMEs using likelihood ratio tests using the Chi Square value, *X*^2^, as the test statistic (implemented using the *anova* function from the *lme4* R package).

## Data Availability

Data are publicly available and can be downloaded via the following repository: https://github.com/gretatuckute/drive_suppress_brains/ (available upon publication).

## Code Availability

Code is publicly available in the following repository: https://github.com/gretatuckute/drive_suppress_brains/ (available upon publication).

## Acknowledgements

This work was supported by the Amazon Fellowship from the Science Hub (administered by the MIT Schwarzman College of Computing) (G.T.), the International Doctoral Fellowship from American Association of University Women (AAUW) (G.T.), the K. Lisa Yang ICoN Center Graduate Fellowship (G.T.), MIT-IBM Watson AI Lab (S.S.), NIH awards R01-DC016607 (E.F.), R01-DC016950 (E.F.), and U01-NS121471 (E.F.), as well as funds from the McGovern Institute for Brain Research (E.F.), the Simons Center for the Social Brain (E.F.), and the Brain and Cognitive Sciences Department (E.F.). The funders had no role in study design, data collection and analysis, decision to publish or preparation of the manuscript.

We thank Colton Casto, Elizabeth Lee, and Edward Gibson for their help on the project; Cory Shain for comments on an earlier draft of the manuscript; and Nancy Kanwisher, N Apurva Ratan Murty, Noga Zaslavsky, Kohitij Kar, Jacob Prince, Josh McDermott, Benjamin Lipkin, Anna Ivanova, and Una-May O’Reilly for valuable discussions.

## Author Contributions Statement

*Conceptualization:* G.T., M.S., E.F. *Methodology*: G.T., M.S., K.K., E.F. *Software*: G.T., A.S., S.S. *Validation*: G.T., A.S., S.S., M.W. *Formal Analysis:* G.T., A.S., S.S., M.W. *Investigation (data collection)*: G.T., A.S., M.T. *Data Curation:* G.T., M.T. *Writing - Original Draft*: G.T., E.F. *Writing – Review and Editing*: G.T., A.S., S.S., M.T., M.W., M.S., K.K., E.F. *Visualization:* G.T. *Supervision*: E.F., K.K. *Project Administration*: E.F. *Funding Acquisition:* E.F.

