## Supplemental Information (SI) for "Driving and suppressing the human language network using large language models"

### Supplementary Information (SI)

### Sections related to encoding model development

#### SI 1: Experimental materials: *Baseline* sentences

We selected a large sample (n=1,000) of naturally occurring sentences in order to probe brain responses to diverse linguistic stimuli. (Note that although not every stimulus constituted a sentence by some definitions, we use the term 'sentence' throughout for convenience.) As detailed below, this set consists of two subsets: the first subset (n=534 sentences) was selected to maximize semantic diversity, and the second subset (n=466 sentences) was selected via random sampling across diverse genres and styles. Across the two subsets, sentences were drawn from nine corpora. All corpora were filtered to only include 6-word sentences with only printable ASCII characters and had a letter as the first character. We chose to use sentences of fixed length because longer connected linguistic strings elicit higher magnitude of response in the language areas [1]. And we chose the length of 6 words because a) we wanted to maximize the number of sentences that could be presented, and a larger number is feasible with shorter stimuli; b) the temporal receptive window (e.g., [2,3]) of the language system appears to be ~6 words (e.g., [4,1,5]).
The corpora were preprocessed to remove repeated/leading/trailing whitespace, strip whitespace before common punctuation characters (?.!,:;), append a final period if the last sentence character was not a punctuation character, and uppercase the first letter.

**Subset 1: Maximizing semantic diversity**
To select a semantically diverse set of sentences, we made use of the 180 semantic clusters identified by Pereira et al. [6] (publicly available at https://osf.io/crwz7/wiki/home). These clusters were derived by performing spectral clustering [7] on GloVe co-occurrence vectors [8] for n=29,805 English words [9] and spanned diverse semantic categories, including some that corresponded to classic concrete concept categories (like 'body' or 'food') and others that were more abstract (like 'soul' or 'argument').
To select the sentences for this subset, we used a large collection of amateur fiction (The Toronto Book Corpus [10]) from three genres: Adventure (n=62,661 sentences after filtering, as described above), Fantasy (n=488,841 sentences), and Mystery (n=177,523 sentences).

For each of the 180 clusters, we took the cluster target word (a word manually assigned as a descriptive label for each cluster in Pereira et al. [6]) and 20 most frequent cluster member words and computed their average GloVe (840B) embedding (either 20 or 21 unique words per cluster, depending on whether the cluster target word belonged to the 20 most frequent cluster member words). GloVe embeddings were available for all 3,769 unique cluster words (169 clusters x 21 words each + 11 clusters x 20 words). For each sentence from each of the three corpora (Toronto Adventure, Toronto Fantasy, Toronto Mystery), we computed the average GloVe embedding of its content words (nouns, verbs, adjectives and adverbs, as identified using the POS tagger from the NLTK Python library [11]); for the sentences that contained no content words (fewer than 0.02% of all sentences within each corpus), we used all words in the sentence.

Next, we computed the pairwise cosine distance between each candidate sentence from the three corpora and each cluster. For each corpus, we outputted the 20 sentences that were most semantically related to each cluster (i.e., had the smallest Cosine distance to the cluster's average embedding) and manually selected one sentence per corpus per cluster. The manual selection process ensured that the sentence is topically related to the target cluster and is not inappropriate/offensive. To minimize idiosyncratic biases, sentences from each of the three corpora were selected by three different people. For Toronto Adventure and Toronto Mystery, no appropriate sentences could be found for two clusters ('pleasure' and 'stupid'), thus yielding 178 sentences from those two corpora. For Toronto Fantasy, no appropriate sentences could be found for two clusters ('pleasure' and 'sexy'), thus yielding 178 sentences from that corpus. Thus, we obtained 534 sentences (178 sentences * 3 corpora) to maximize semantic diversity.

**Subset 2: Random sampling**

The remaining 466 sentences were randomly sampled from eight diverse corpora that spanned three main categories:

1) **Published written text:** Subset of The Wall Street Journal articles published in 1996 (n=2,997 sentences; [12]), The Brown Corpus (n=1,421 sentences; [11]), Subset of The Universal Dependencies Corpus (n=1,063 sentences; https://universaldependencies.org/), The Contract Corpus (n=264 sentences; legal texts from Goźdź-Roszkowski [13] and texts collected by Martinez, Mollica & Gibson [182]), and The Colorado Richly Annotated Full-Text (CRAFT) Corpus (n=34 sentences; biomedical articles from PubMed; [15]).
2) **Web media text**: Subset of The Common Crawl C4 Corpus (n=921,099 sentences; [16]).
3) **Transcribed spoken text**: The Cornell Movie-Dialogs Corpus (n=41,058 sentences; [17]), and a subset of the spoken component of The Corpus of Contemporary American English (COCA) (n=101,193 sentences; [18]).

The total number of sentences across these eight corpora was 1,069,129. Given that the corpora varied substantially in size (between 34 and 921,099), we implemented a stratified sampling scheme to ensure that sentences were sampled from all eight corpora. Specifically, for corpora with fewer than 0.025% of all sentences (two corpora: The Contract Corpus and The CRAFT Corpus), we sampled 5 sentences from each. For corpora with 0.1-0.3% of all sentences (three corpora: The Wall Street Journal Corpus, The Brown Corpus, The Universal Dependencies Corpus), we sampled 50 sentences each. For corpora with 3.8-9.5% of all sentences (two corpora: The Cornell Movie-Dialogs Corpus and COCA), we sampled 75 sentences each. This sampling yielded 310 sentences. The remaining 156 sentences were sampled from the remaining (largest) corpus (The Common Crawl C4 Corpus) with 86.2% of all sentences. Thus, we obtained 466 sentences from diverse corpora that spanned different genres (e.g., written vs. spoken language) and styles (e.g., formal vs. conversational language).

After collecting fMRI data from the first participant (unique ID 853; these are numbers in the lab-internal database and can be cross-referenced with the Fedorenko lab's studies on OSF), several minor edits and replacements were made to the original set of 1,000 sentences. Twenty unique sentences were affected: for 8 sentences, we made minor changes to the punctuation, and 12 sentences were replaced (with sentences from the original sentence sources, as described above, and according to the manual filtering criteria reported in SI 2).

#### SI 2: Sentence exclusion criteria

The manual sentence exclusion criteria were defined prior to selection of any materials in the study. The automatic exclusion criteria were defined prior to selection of *drive* and *suppress* sentences. In the following section, the term “token’’ refers to groups of alphanumeric characters separated by whitespace.

##### Manual exclusion criteria

- The sentence is inappropriate/offensive (e.g., contains a racial slur or a taboo word).
- All tokens in the sentence are not English.
- The sentence contains some type of emoji face, e.g., “:)”.
- All tokens in the sentence are trademarks, website names, brand names, product names, person names, journal identifiers, journal footers, date and geographical locations.
- The sentence contains more or fewer than 6 words due to tokenization/punctuation errors.

Additionally, the following three criteria were applied to the *baseline* materials (n=1,000):

- The sentence contains several sub-sentences (two sentences in one marked by a period, an exclamation point, or a question mark).
- The sentence has a grammatical error (e.g., agreement error).
- The sentence has a typo.

##### Automatic exclusion criteria

- More than 50% of tokens in the sentence contain numerical characters.
- More than 50% of all characters in the sentence are uppercase.
- The sentence contains one or more tokens that are longer than 20 characters.
- The sentence contains more than 4 consecutive punctuation characters.
- The sentence contains non-ascii characters (unicode index larger than 127) or a character in the following set: *, @, [, ], \, /, <, >, =, ^, _, `, {, }, |, ~

#### SI 3: Joint fMRI data modeling and data evaluation framework for robust estimation of single-trial fMRI responses

We developed an experimental paradigm to collect brain responses in an event-related fMRI design where each unique trial (i.e., sentence) was presented on the screen one at a time for 2s with 4s inter-stimulus interval (**SI Figure 3A**, Step 1 and Methods; fMRI experiments). Data analysis using these rapid, event-related designs is complex due to temporal signal autocorrelation, participant head motion, and scanner instabilities [19] and is therefore dependent on accurate general linear model (GLM) beta estimates of each trial. To obtain as accurate and robust responses as possible, we utilized a state-of-the-art framework for event-related modeling, GLMsingle [19]. As described in Prince et al. [19], GLMsingle relies on internal cross-validation to estimate two key modeling hyperparameters (the number of noise regressors and the voxel-wise levels of ridge regression regularization). This cross-validation requires at least some repeated presentations of the same stimulus in an experiment, the underlying assumption being that repetitions of the same stimulus should produce similar neural responses. In the current study, we did not want to make this assumption and presented each stimulus only once for a given participant (see Discussion in the main text). In order to exploit the denoising benefits of GLMsingle, we coupled the GLMsingle framework to a downstream data evaluation metric, as described below.

The premise of the proposed procedure is to exploit inter-participant similarity in responses in functionally defined brain areas in order to set the two hyperparameters of interest, as illustrated in **SI Figure 3A**.

First, we modeled each of the first 5 participants' (*train* participants) fMRI data using different GLMs each with a unique combination of the two hyperparameters (number of noise regressors and ridge regression regularization fraction) (**SI Figure 3A**; Step 2, e.g., one GLM instantiation may use 5 noise regressors and a regularization fraction of 0.4). Specifically, we ran 126 different GLM instantiations per participant spanning 1-10 noise regressors and a ridge regression fraction of 0.05 to 1.

For each GLM instantiation, we extracted responses from the functionally defined left hemisphere language network (**SI Figure 3A**; Step 3), defined as the top 10% most language-responsive voxels within pre-defined anatomical parcels using an extensively validated language localizer task (Methods; Definition of ROIs). For each participant, we computed the arithmetic average of the language-selective voxels in the IFGorb, IFG, MFG, AntTemp, and PostTemp fROIs for all sentence trials. Next, we quantified how correlated the sentence-level responses were across the 5 participants, yielding a correlation value for each participant pair (**SI Figure 3A**; Step 4). To quantify the overall consistency across participants, we computed the median of the pairwise participant correlation values. This procedure was performed for each GLM hyperparameter instantiation, resulting in a median inter-participant correlation per GLM model instantiation. The highest inter-participant correlation (Pearson r=0.095) was obtained using 5 noise regressors and a ridge regularization fraction of 0.05 (**SI Figure 3B**). This GLM hyperparameter combination was thus selected as the GLM data instantiation for use.

The 5 participants' data used for this procedure were fixed using this set of identified modeling hyperparameters, and new participants' data (*evaluation* participants) were modeled using the same set of parameters. The assumption underlying this joint data modeling and evaluation framework is that some level of participant-to-participant consistency is expected, particularly for functionally defined brain areas [20–23]. Hence, any method that demonstrably improves participant-to-participant consistency is likely to be a sound denoising method [24]. To sum up, the coupling of GLM hyperparameters to the downstream inter-participant evaluation metric allows for the usage of state-of-the-art single-trial modeling tools [19] with single stimulus repetition designs and more generally, foregoing the assumption that the same stimulus should elicit the same response across repetitions.

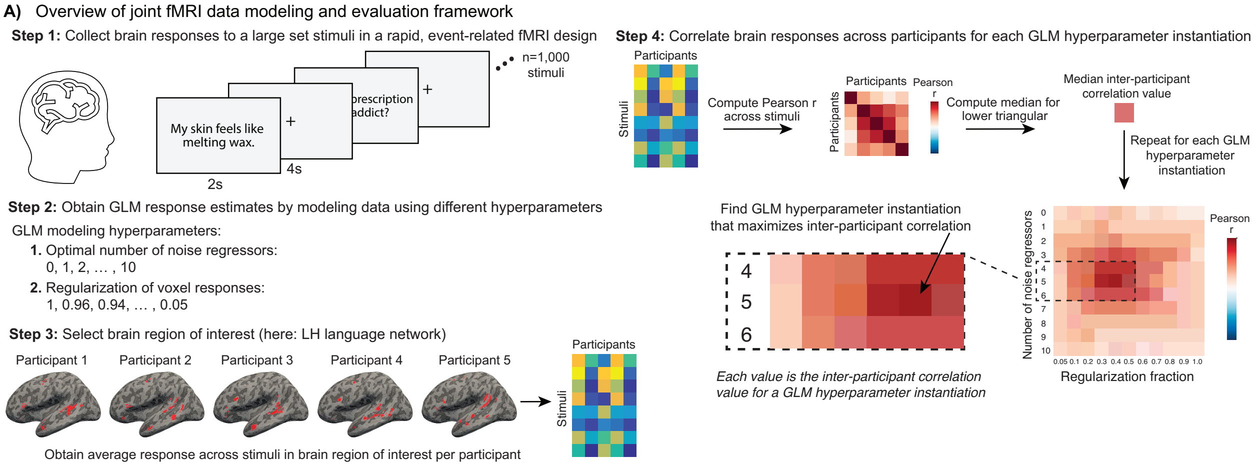

**SI Figure 3A. Overview of methodological approach for estimation of GLM modeling parameters.** The figure is explained in the corresponding text. For illustration purposes, the plots do not show real data.

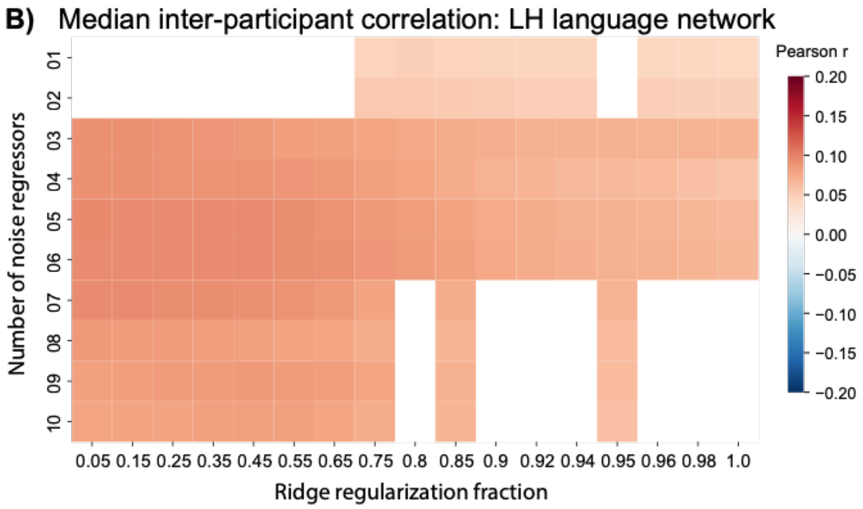

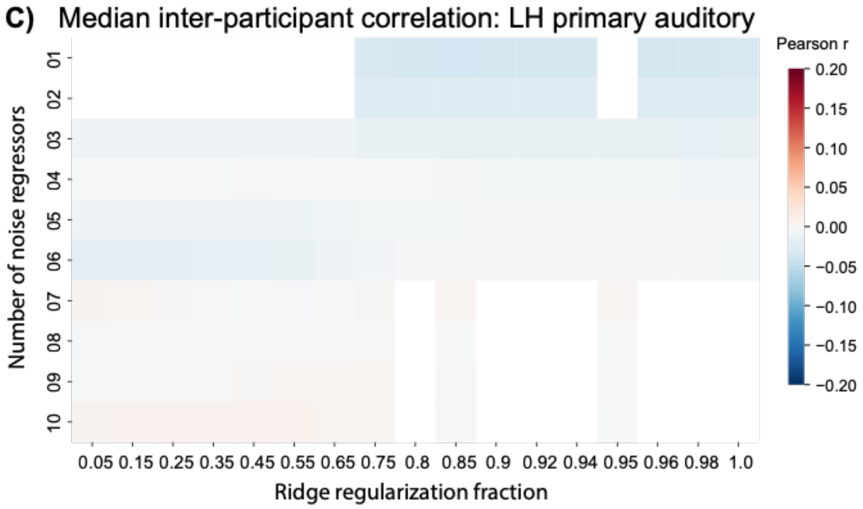

**SI Figure 3B,C. Joint data modeling and evaluation framework successfully maximizes inter-participant correlation in the functionally-defined LH language network, but not in a control brain region (primary auditory).**

Each value in the grids represents the median inter-participant correlation (n=5 participants) across n=1,000 sentence trials for different GLM instantiations. Each GLM instantiation used a different combination of two GLM modeling hyperparameters: Number of noise regressors (y-axis) and ridge regularization fraction (x-axis; ranges from 0 (maximal regularization) to 1 (no regularization, OLS solution); [190]). There were 126 unique combinations in total. The white parts of the grid were not computed given high computational cost for each data point.

**B)** Median inter-participant correlations from the functionally defined left hemisphere language network (the average of the top 10% language-selective voxels in the IFGorb, IFG, MFG, AntTemp, and PostTemp fROIs within pre-defined anatomical parcels, defined using an independent language localizer task; Methods; Definition of ROIs). The highest inter-participant correlation (Pearson r = 0.095) was obtained using 5 noise regressors and a ridge regularization fraction of 0.05.

**C)** Median inter-participant correlations from the anatomically defined left hemisphere primary auditory region, included as a control region where there is no expectation about inter-participant correlations (the mean of the voxels in the anatomically defined left TE1.1 and TE1.2 regions based on human post-mortem histology [214]). The highest inter-participant correlation (Pearson r=0.01) was obtained using 10 noise regressors and a ridge regularization fraction of 0.35.

### SI 4: Correlation across sentences for LH language fROIs

**A)** Correlation across n=1,000 *baseline* sentences for LH language fROIs (*train* participants)

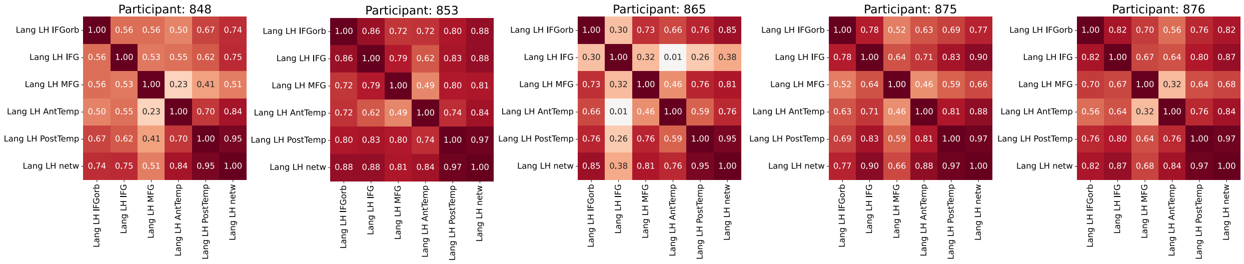

**B)** Correlation across n=1,500 *baseline, drive,* and *suppress* sentences for LH language fROIs (*evaluation* participants)

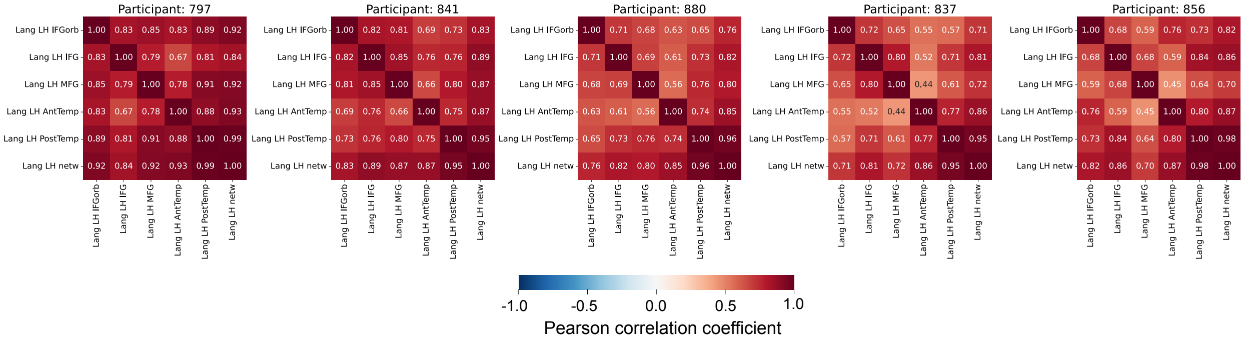

**SI Figure 4. LH language regions are highly correlated across both naturalistic and model-selected sentences.**

**A)** The Pearson correlation matrix computed over n=1,000 *baseline* sentences for each of n=5 *train* participants. The first five rows/columns show the five core LH language fROIs (IFGorb, IFG, MFG, AntTemp, and PostTemp; Methods; Definition of ROIs). The sixth row/column shows the LH language fROI consisting of the average of the voxels from the five fROIs.

**B)** Correlation matrices computed over n=1,500 *baseline, drive* and *suppress* sentences for each of n=5 *evaluation* participants. The first three participants (797, 841, 880) were exposed to *drive/suppress* materials derived from the *search* approach, while the next two participants (837, 856) were exposed to *drive/suppress* materials derived from the exploratory *modify* approach (SI 16).

Note that the participant numbers refer to unique identifiers in the lab internal database and can be cross-referenced with the Fedorenko lab's studies on OSF.

#### SI 5: Noise ceiling

##### SI 5A: Estimation of noise ceiling

We estimated noise ceilings for the regions of interest (ROIs) in the brain data. We define the noise ceiling as the theoretical upper limit for the correlation between an external predictor and brain ROI measurements, given the presence of measurement noise. The noise ceiling depends solely on the signal-to-noise ratio of the data and is independent of the specific external predictor (i.e., LLM model) being evaluated. Given that the primary modeling work in the current study was performed on participant-averaged data (specifically, the average of brain data from the five *train* participants), we estimated the noise ceiling for this participant-averaged ROI-level data. Our method for estimating the noise ceiling follows the general framework laid out in previous studies [27–29] with one core difference: The previous studies make use of the variability across repeated presentations of the same trial to estimate the noise ceiling, while in the current work we make use of the repeated presentations of the same trial across participants, as detailed below.

In the noise ceiling estimation procedure, we make the following four assumptions: 1) the signal contained in the ROI's response is determined solely by the presented sentence, 2) the variability of the ROI's response across different sentences is Gaussian distributed, 3) the noise (i.e., variability of the response to a given sentence across participants) is Gaussian distributed with zero mean, and 4) the response to a sentence is equal to the signal plus noise. Given these assumptions, any observed response for a single subject is a sample from a sum of Gaussian distributions:

$$\mathrm{RESP} \sim \mathcal{N}(\mu_{\mathrm{signal}}, \sigma_{\mathrm{signal}}) + \mathcal{N}(0, \sigma_{\mathrm{noise}}) \quad (1)$$

Where $RESP$ indicates the observed BOLD response for a single subject. $\mu_{signal}$ is the mean ROI-level BOLD response signal across different sentences, $\sigma_{signal}$ is the standard deviation of the signal across different sentences, and $\sigma_{noise}$ is the standard deviation of the noise (across participants).

The data used to compute the noise ceiling consists of the ROI responses across 1,000 sentences ($N_s$ = 1,000) for five participants ($N_p$ = 5). We denote the participant-averaged data as $RESP_{avg}$ (i.e., a column vector). Each sentence was presented once for each participant. Note that we treat the ROI-level sentence responses across participants as different 'repetitions' of a given sentence.

The estimation of the noise ceiling proceeds in 3 steps: i) Estimation of the noise standard deviation, ii) Estimation of the signal standard deviation, iii) Estimation of signal-to-noise and noise ceiling.

*i) Estimation of the noise standard deviation*

We first compute the standard deviation of the distribution that characterizes the noise. To do so, we start by calculating the variance of the ROI responses across the $N_p$ presentations of each sentence (using the unbiased estimator that normalizes by $N_p - 1$ where $N_p$ is the population, i.e., number of participants). We then average this variance across sentences and compute the square root of the resulting number. This produces an estimate of the noise standard deviation:

$$\hat{\sigma}_{\text{noise}} = \sqrt{\text{mean}(\beta_\sigma^2)} \quad (2)$$

where $\beta_\sigma^2$ indicates the variance across the ROI responses observed for a given sentence across participants. Intuitively, the noise standard deviation ($\hat{\sigma}_{noise}$) reflects the noise that is contributed by participants.

*ii) Estimation of the signal standard deviation*

Second, we compute the standard deviation of the distribution that characterizes the signal. To do so, we average the ROI responses across the $N_p$ presentations of each sentence (i.e., across participants, and then calculate the variance across sentences ($\hat{\sigma}^2_{RESP_{avg}}$) (using the unbiased estimator that normalizes by $N_s - 1$ where $N_s$ is the number of sentences). This quantifies variance in the data. Then, we subtract the variance that is contributed by noise:

$$\hat{\sigma}_{\text{signal}} = \sqrt{\left| \hat{\sigma}^2_{\text{RESP}_{\text{avg}}} - \frac{\hat{\sigma}^2_{\text{noise}}}{N_p} \right|_+} \quad (3)$$

where $\hat{\sigma}_{signal}$ is the signal standard deviation, $\hat{\sigma}^2_{RESP_{avg}}$ is the variance of the group-averaged data, $\hat{\sigma}^2_{noise}$ is the variance of the noise distribution (as computed in Eq. (2)), and ||+ indicates positive half-wave rectification. Note that $\hat{\sigma}^2_{noise}$ is divided by the number of participants because $\hat{\sigma}^2_{noise}$ was derived using a population of $N_p$ participants.

*iii) Estimation of signal-to-noise and noise ceiling*

We express the noise and signal standard deviations computed in respectively Eq. (2) and (3) as a single scalar noise ceiling (NC) signal-to-noise (SNR) ratio (a value between 0 and 1):

$$\text{NCSNR} = \frac{\hat{\sigma}_{\text{signal}}}{\hat{\sigma}_{\text{noise}}} \quad (4)$$

We want to express the noise ceiling as the amount of variance contributed by the signal as a fraction of the total amount of variance in the data (see the NSD Data Manual for derivation https://naturalscenesdataset.org; [29])

$$\mathrm{NC} = \frac{\mathrm{NCSNR}^2}{\mathrm{NCSNR}^2 + \frac{1}{N_p}} \quad (5)$$

where $N_p$ indicates the number of participants that are averaged together. The $N_p$ parameter allows to flexibly express the noise ceiling for different levels of participant averaging (e.g., for averaging trials across 2, 3, …, $N_p$).

Finally, to make the noise ceiling estimate comparable to the encoding model performance Pearson correlation scores, we convert this to Pearson correlation units by taking the square root:

$$\mathrm{NC}_{\mathrm{Pearson}} = \sqrt{\mathrm{NC}} \quad (6)$$

Using this approach, $NC_{Pearson}$ was estimated to be 0.56 for the LH language network. Note that this framework treats response variability that is unrelated to the stimulus and not shared across participants as ‘noise’, but such variability might reflect both non-stimulus related activity (e.g., head motion, physiological noise) and true neural variability across participants.

#### SI 5B: Estimation of noise ceiling reliability

The noise ceiling computed in Section SI 5A is itself an estimate that is dependent on the data that was used to compute the noise ceiling (in our case, $N_s$= 1,000 sentences across $N_p$ = 5 participants). We quantified the reliability of the noise ceiling using a split-half procedure: across 1,000 iterations, we partitioned the sentences into two independent, random sets of 500 sentences each and estimated the noise ceiling for each half as described in SI 5A. For each iteration, this step yielded two independent noise ceiling estimates. We computed the standard deviation across these two noise ceiling estimates (using the unbiased estimator, $N_{split} - 1$, *where* $N_{split}$ = 2) per iteration, leaving us with 1,000 standard deviation values. To obtain a standard error estimate, we averaged the 1,000 standard deviation values (by squaring the standard deviations, averaging, and transforming back to standard deviations by taking the square root). Finally, we divided this average standard deviation by the square root of $N_{split}$ (i.e., $\sqrt{2}$) and used this as a measure of the noise ceiling reliability.

#### SI 6: Cross-validated encoding model performance on held-out sentences

To obtain an unbiased estimate of encoding model performance, we implemented a 5-fold cross-validation procedure using held-out sentences (80%–20% train-test splits; i.e. 800 of the 1,000 *baseline* sentences in the train split and 200 sentences in the test split).
For a given brain region of interest (ROI), we fitted the ridge regression model from the LLM's representations of the training sentences to the ROI's corresponding brain recordings for those sentences (participant-averaged brain data from n=5 *train* participants). The $\alpha$ regularization parameter was identified using leave-one-out cross-validation on the training split. We applied the regression model with the identified regularization parameter on LLM representations of the held-out 20% of sentences to generate predicted brain responses for those sentences. Performance of the model was evaluated by correlating (Pearson r) the predicted ROI response with the observed ROI response. If there was no variance in the predicted ROI responses across sentences (defined as a standard deviation less than 1e-7 of the predicted ROI responses), the Pearson correlation coefficient was set to 0. This process was repeated five times, holding out a different 20% of sentences in each fold. For a given ROI, we then took the mean of the resulting five Pearson correlation scores to give us a mean predictivity score and computed the standard error of the mean over the five cross-validation folds.

We emphasize that the test sentences on which predictions were ultimately evaluated were not incorporated into the procedure for selecting the $\alpha$ regularization parameter nor for estimating the linear mapping from LLM features to an ROI's response – i.e., the procedure was fully cross-validated.

The following subsections show the cross-validated predictivity performance using different sentence representations from GPT2-XL (SI 6A, SI 6B) and from a different language model (a bidirectional attention Transformer: BERT-large) (SI 6C). Finally, in SI 6D, we show predictivity performance on anatomically defined language regions using GPT2-XL features.

All modeling and analysis code was written in Python (version 3.8.11), making heavy use of the *numpy* ([30]; version 1.21.2), *scipy* ([31]; version 1.7.3), *scikit-learn* ([32]; version 0.24.2), *pandas* ([33]; version 1.4.2) and *transformers* ([34]; version 4.11.3) libraries.

SI 6A: GPT2-XL (primary approach: last token representation)

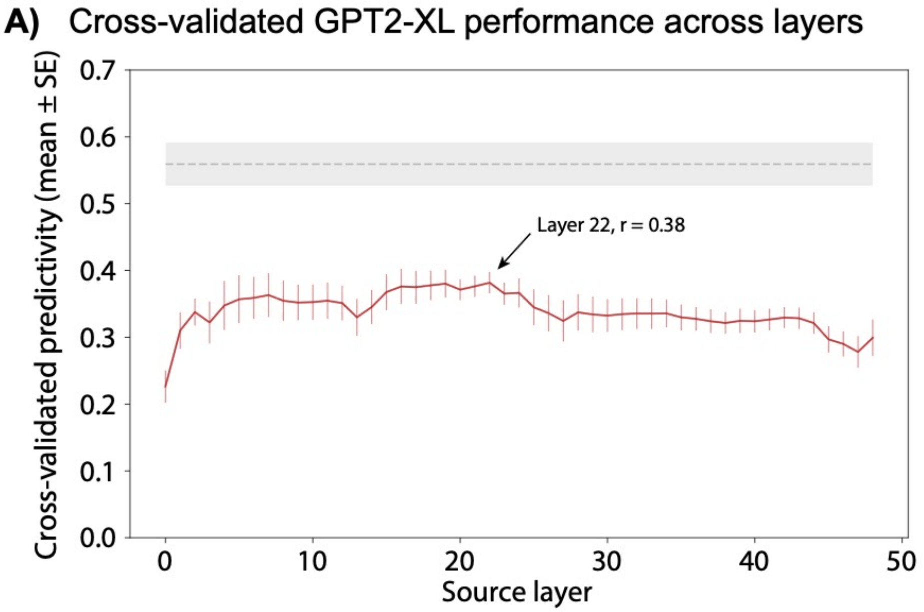

**SI Figure 6A. Cross-validated GPT2-XL performance across layers using last-token GPT2-XL representations.**

We obtained cross-validated predictivity performance (y-axis) of the encoding model for each layer (i.e., Transformer block) of GPT2-XL (x-axis) using the primary sequence summary representation for each sentence: last token (Methods; Encoding model development). The error bar shows the standard error of the mean across five cross-validation folds (n=200 sentences in each test fold; a total of n=1,000 *baseline* sentences in the cross-validation analysis). The dashed grey line shows the noise ceiling of the LH language network with shaded regions showing the noise ceiling reliability (split-half standard error; SI 5B). The highest prediction performance was obtained using layer 22 (arrow; r = 0.38) and hence features from this layer were used as regressors for the encoding model.

SI 6B: GPT2-XL (additional approach: average token representation)

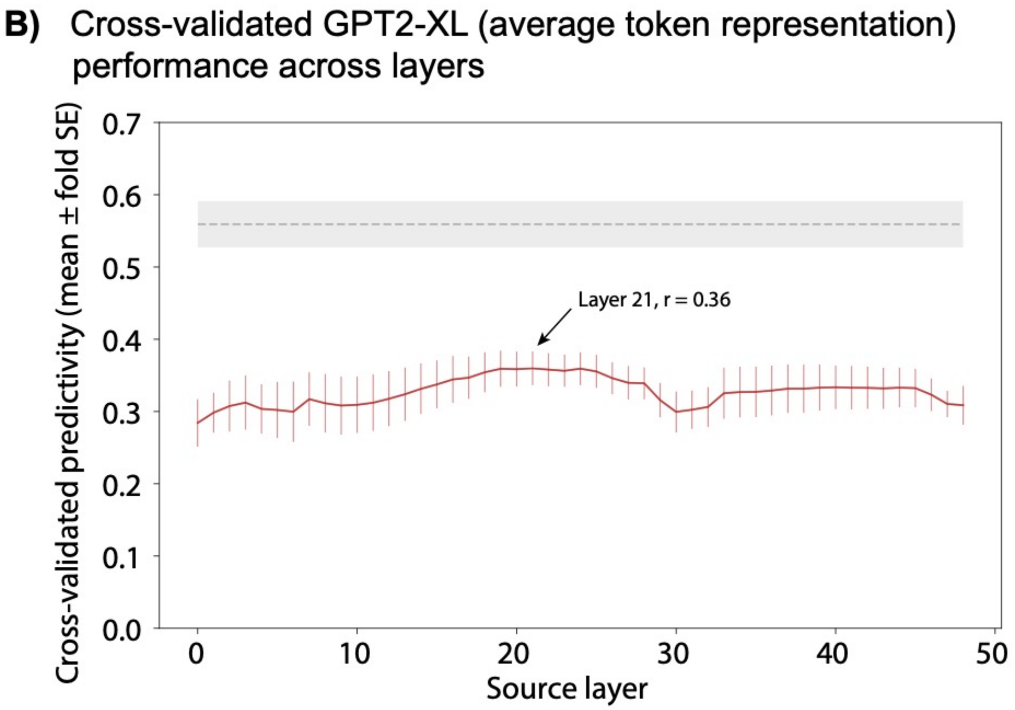

**SI Figure 6B. Cross-validated GPT2-XL performance across layers using average token GPT2-XL representations.**
We tested the robustness of our encoding model by extracting sentence representations from GPT2-XL using a different sequence summary method: instead of obtaining the last token representation (SI Figure 6A), we computed the average of all tokens in each sentence. Identical to SI Figure 6A, we obtained cross-validated predictivity performance (y-axis) of the encoding model for each layer. The error bar shows the standard error of the mean across five cross-validation folds (n=200 sentences in each test fold; a total of n=1,000 *baseline* sentences in the cross-validation analysis). The dashed grey line shows the noise ceiling of the LH language network with shaded regions showing the noise ceiling reliability (split-half standard error; SI 5B). The highest prediction performance was obtained using layer 21 (arrow; r = 0.36).

#### SI 6C: BERT-large

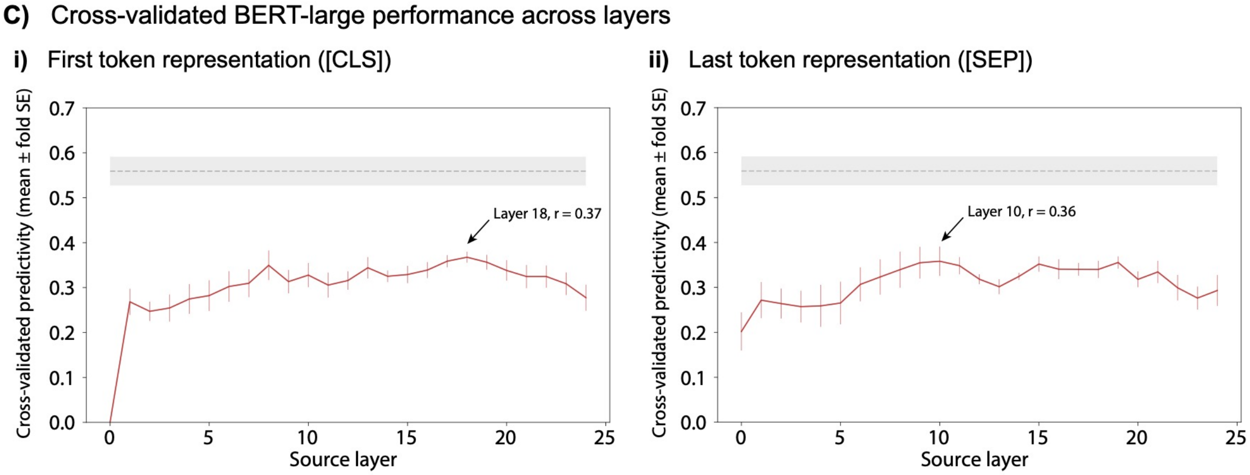

**SI Figure 6C. Cross-validated BERT-large performance across layers using two different token representations.**

We tested the robustness of our encoding model by extracting sentence representations from a different LLM architecture, a bidirectional-attention Transformer model: BERT-large-cased [35] (from two different special tokens; respectively panel **i** and **ii**). The sentence representations were obtained from the pretrained model available via the HuggingFace library ([34]; *transformers version* 4.11.3; https://huggingface.co/bert-large-cased). BERT-large-cased has 24 layers (i.e., Transformer blocks) in addition to the embedding layer. The embedding dimension is 1,024. We obtained model representations by tokenizing each sentence using the model's standard tokenizer (BertTokenizerFast) and passing each sentence through the model. We retrieved model representations for each model layer (i.e., at the end of each Transformer block). We obtained a sequence summary representation of each sentence using two approaches: In panel **i**, we used the special classification, [CLS], token which is prepended to the sequence and is standardly used as a token for classification output [35], and in panel **ii**, we used the special separator token, [SEP], which is appended to the sequence and standardly used to separate input [35]. The highest prediction performance for the [CLS] token method (panel **i**) was obtained using layer 18 (arrow; r = 0.37), and the highest prediction performance for the [SEP] token method (panel **ii**) was obtained using layer 10 (arrow; r = 0.36). The error bar shows the standard error of the mean across five cross-validation folds (n=200 sentences in each test fold; a total of n=1,000 *baseline* sentences in the cross-validation analysis). The dashed grey line shows the noise ceiling of the LH language network with shaded regions showing the noise ceiling reliability (split-half standard error; SI 5B).

SI 6D: GPT2-XL (anatomically defined LH language network)

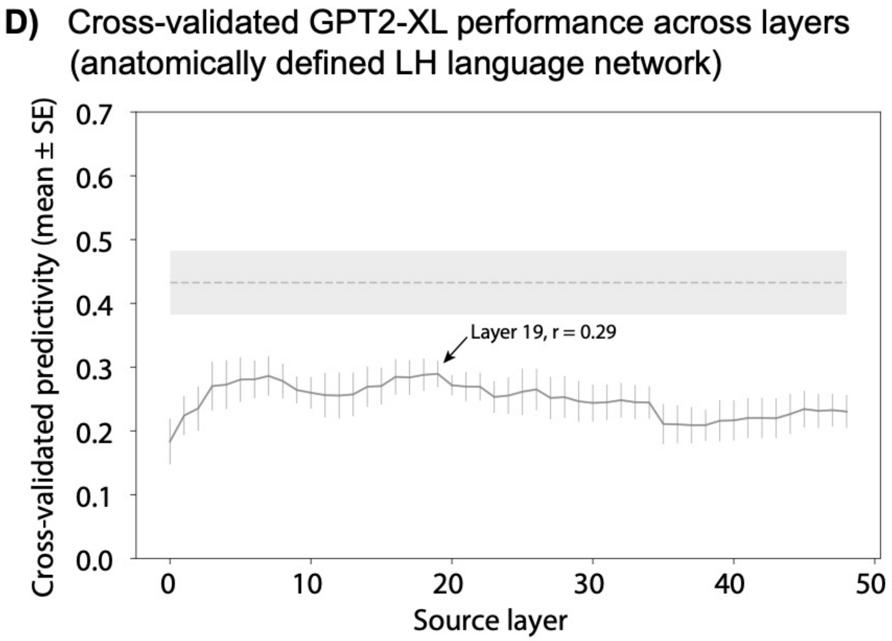

**SI Figure 6D. Cross-validated GPT2-XL performance across layers using last-token GPT2-XL representations for the anatomically defined language network.**

We tested whether our encoding model (using the primary GPT2-XL last token approach) could predict language regions defined anatomically (as opposed to functionally defined in each participant; Methods; Definition of ROIs). To define the anatomical language network, we used the Glasser parcellation [36] to select a subset of ROIs that approximately correspond to the language network. We identified parcels that overlapped by at least 25% of voxels with one of the five anatomical LH language parcels (Methods; Definition of ROIs), as was done in Lipkin et al. [37], resulting in n=21 parcels in total for the LH language network. Between 2 and 8 Glasser parcels overlapped with each of the five language parcels, as detailed below.

Our parcel: corresponding Glasser parcels
LangIFGorb: 47l, 45
LangIFG: IFSp, IFJa, 44
LangMFG: FEF, 55b
LangAntTemp: TA2, STSva, STSda, STGa, PI, A5
LangPostTemp: TPOJ2, TPOJ1, STV, STSvp, STSdp, PSL, PHT, pGi

The highest prediction performance was obtained using layer 19 (arrow; r = 0.29). The error bar shows the standard error of the mean across five cross-validation folds (n=200 sentences in each test fold; a total of n=1,000 *baseline* sentences in the cross-validation analysis). The dashed grey line shows the noise ceiling of the anatomical LH language network with shaded regions showing the noise ceiling reliability (split-half standard error; SI 5B).

#### SI 7: Cross-validated encoding model performance on held-out participants

To obtain an unbiased estimate of encoding model performance, we implemented a cross-validation procedure using held-out participants from the set of n=5 *train* participants (five such combinations. For model fitting, we averaged the data from four of the participants and for model testing, we left out a different participant each time).
For the LH language network, we fitted the ridge regression model from the LLM's representations of the n=1,000 *baseline* sentences to the language network's corresponding brain recordings for those sentences (the average language network's response across n=4 participants in the train split). The $\alpha$ regularization parameter was identified using leave-one-out cross-validation on the training participants. We used the regression model with the identified regularization parameter to generate predicted brain responses for the n=1,000 *baseline* sentences. Performance of the model was evaluated by correlating (Pearson r) the predicted language network response with the observed language network response of the held-out participant. If there was no variance in the predicted language network responses across sentences (defined as a standard deviation less than 1e-7 of the predicted language network responses), the Pearson correlation coefficient was set to 0.
This process was repeated five times, holding out a different participant each time. We then took the mean of the resulting five Pearson correlation scores to give us a mean predictivity score, and computed the standard error of the mean over the five cross-validation combinations. We emphasize that the test stimuli on which predictions were ultimately evaluated were not incorporated into the procedure for selecting the $\alpha$ regularization parameter nor for estimating the linear mapping from LLM features to the language network's response – i.e., the procedure was fully cross-validated.

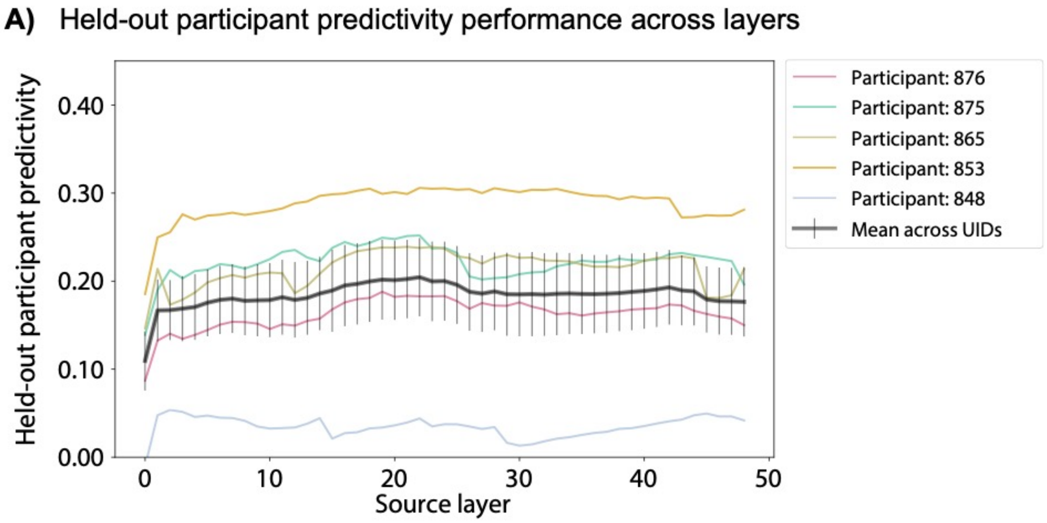

**SI Figure 7. Held-out participant GPT2-XL performance across layers using last-token GPT2-XL representations.**
We estimated the predictivity of an encoding model (using the primary GPT2-XL last token approach) trained on the average of 4 participants' brain data, and predicting on a single, held-out participant (within the *train* participants) using the full *baseline set* (n=1,000 sentences). Colored lines show prediction performance on each held-out participant, and the black line shows the mean across participants with error bar showing the standard error of the mean across participants.

#### SI 8: Noise ceiling as a function of encoding model performance

We examined how well GPT2-XL features can explain brain responses relative to the noise ceiling in language regions and a set of control brain regions. The control brain regions included i) two large-scale brain networks that have been linked to high-level cognitive processing—the multiple demand (MD) network [38–42] and the default mode network (DMN) [43–47]—which we defined using independent functional localizers (see SI 15 for details) and ii) a set of anatomical parcels [36] that cover a large fraction of the cortical surface.

The noise ceiling (NC) for a brain region is a measure of stimulus-related response reliability and provides an upper bound on the amount of variance that any external predictor can theoretically explain. We computed the NC from the brain responses to the 1,000 *baseline* sentences for the n=5 *train* participants (SI 5).
We quantified prediction performance by training separate encoding models to predict responses in the language regions as well as control brain regions (defined functionally and anatomically) using features from GPT2-XL on the participant-averaged data from n=5 *train* participants on the *baseline set* (as in the main encoding model approach, see Methods; Encoding model development).

First, we investigated the NC and prediction performance of language regions versus the two functionally defined brain networks (MD and DMN). **SI Figure 8A** shows the prediction performance on held-out sentences plotted against the NC for the functionally defined control brain regions across all 49 layers of GPT2-XL. Language regions were most reliable and well-predicted (red points in the upper right part of the plot). For example, the GPT2-XL features explained 67.9% of the variance in the LH language network (layer 22). The MD and DMN regions were clustered in the lower left part of the plot, characterized by relatively low NC with large uncertainty estimates and low absolute predictivity values.

Next, we investigated the NC and prediction performance of language regions versus a set of anatomical parcels [36]. **SI Figure 8B** shows the NC for the anatomical parcels on the surface-inflated brain. **SI Figure 8C** shows the prediction performance on held-out sentences plotted against the NC for the anatomical parcels across all 49 layers of GPT2-XL. Similar to **SI Figure 8B**, most parcels were clustered in the lower left part of the plot with relatively low NC and predictivity values. Hence, the language regions were accounted for well by GPT-XL features in a spatially specific manner: other brain areas, including those that support high-level cognition, were not as reliable or well-predicted as language areas.

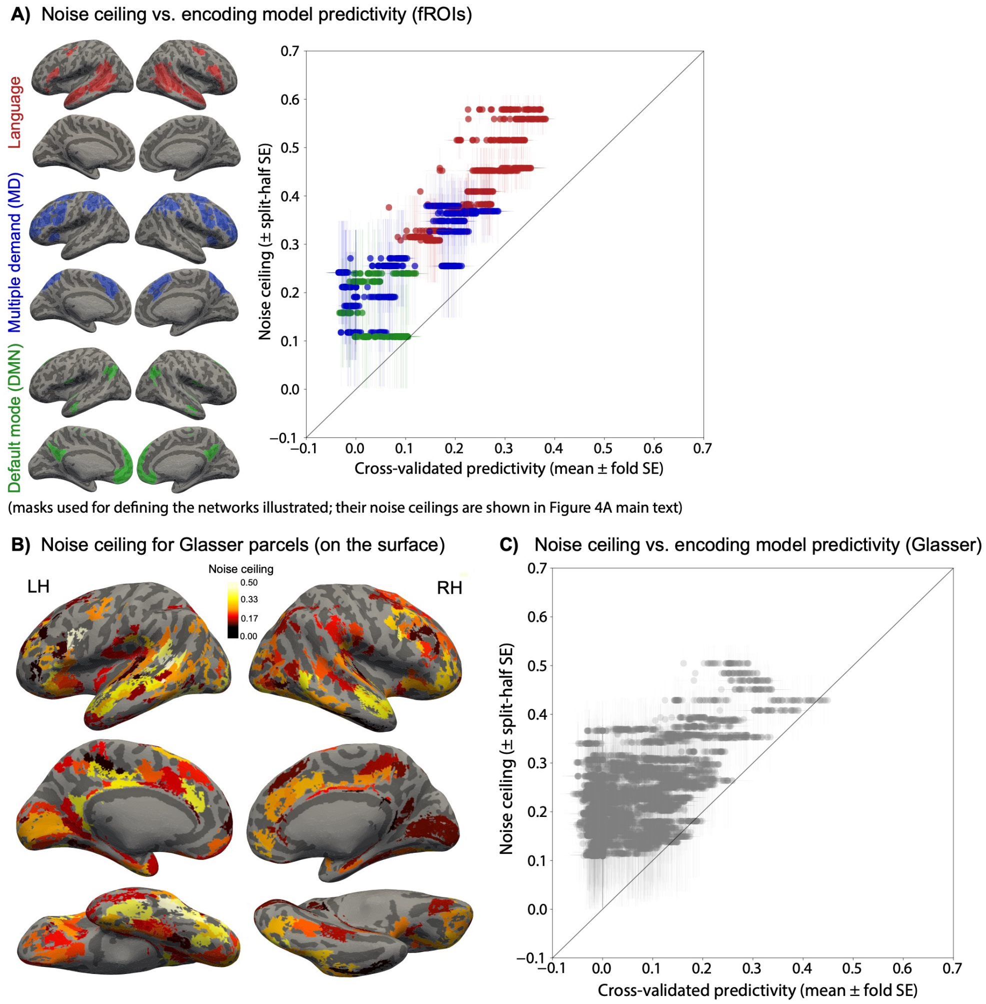

**SI Figure 8. Language regions are reliably well-predicted, better than control brain regions.**

**A)** The noise ceiling (NC; y-axis), a measure of stimulus-related response reliability, as a function of encoding model performance (x-axis) across all functionally defined ROIs in the language network (red), multiple demand (MD) network (blue), and the default mode network (DMN; green). The fROIs for which the NC reliability did not overlap with zero were included in the analysis (30 fROIs in total; red points: language fROIs; blue points: MD fROIs; green points: DMN fROIs). (The NC estimates are plotted in **Figure 4A** in the main text.). Encoding model performance was obtained using 5-fold cross-validation (SI 6). The error bar on the x-axis shows the standard error of the mean across cross-validation folds. Performance for all layers of GPT2-XL is shown (49 layers). The error bar on the y-axis shows the split-half NC reliability.

**B)** We quantified the NC for a set of anatomical parcels that cover the whole brain [36], here illustrated on the surface-inflated MNI152 template brain. The color shows the NC with lighter colors indicating higher values. Anatomical ROIs where the NC split-half reliability overlapped with zero are not shown (leaving 189 anatomical ROIs: 100 in LH; 89 in RH).

**C)** The NC (y-axis) as a function of encoding model performance (x-axis) for the anatomical ROIs for which the NC reliability did not overlap with zero (189 ROIs in total; the NCs for these ROIs are illustrated in panel B). Encoding model performance was obtained using 5-fold cross-validation (see SI 6 for details). The error bar on the x-axis shows the standard error of the mean across cross-validation folds. Performance for all layers of GPT2-XL is shown (49 layers). The error bar on the y-axis shows the noise ceiling reliability.

### Sections related to encoding model evaluation

#### SI 9: Experimental materials: *Drive* and *suppress* sentences

The *drive* and *suppress* materials were designed to elicit high or low fMRI activity in the language network, respectively, based on the predictions from the encoding model (Methods; Encoding model evaluation). The *drive* and *suppress* materials were selected by searching across ~1.8M sentences to identify sentences that are predicted to elicit high or low fMRI responses (the *search* approach).

In a more exploratory component of the study, we complemented the *search* approach with another approach—the *modify* approach—where we used gradient-based modifications to transform a random sentence into a novel sentence/string predicted to elicit high or low fMRI responses (SI 16).

Using each method, we identified 250 unique *drive* and 250 unique *suppress* sentences. These 500 sentences were interspersed among the 1,000 *baseline* sentences (SI 1) in the fMRI scanning sessions (for a total of n=1,500 unique sentences collected across three scanning sessions with 500 unique sentences in each session). This section is focused on the *search* experimental materials, while details on the *modify* materials can be found in SI 16.

##### SI 9A: Identifying candidate materials

The *search* approach searches through large-scale corpora of diverse English text to identify candidate *drive* and *suppress* sentences. Specifically, to identify candidate *drive/suppress* sentences we searched across 1,871,413 6-word sentences (unique sentences: 1,805,623) from nine diverse large-scale text corpora. The corpora were partly overlapping with the corpora described in SI 1 and were filtered and preprocessed in the same manner as the *baseline* materials. The corpora spanned three main categories (see SI Table 9):

1) **Published written text:** Subset of The Toronto Book Corpus [10] from three genres: Adventure (n=62,661 sentences after filtering, as described above), Fantasy (n=488,841 sentences), and Mystery (n=177,523 sentences), Subset of The Wall Street Journal articles published in 1996 (n=2,997 sentences; [12]), The Brown Corpus (n=1,421 sentences; [11]), Subset of The Universal Dependencies Corpus (n=1,063 sentences; https://universaldependencies.org/), The Contract Corpus (n=264 sentences; legal texts from Goźdź-Roszkowski [13] and texts collected by Martinez, Mollica & Gibson [14]), and The Colorado Richly Annotated Full-Text (CRAFT) Corpus (n=34 sentences; biomedical articles from PubMed; [15]).
2) **Web media text**: Subset of The Common Crawl C4 Corpus (n=878,552 sentences; [16]).
3) **Transcribed spoken text**: The Cornell Movie-Dialogs Corpus (n=41,058 sentences; [17]), and the spoken component of The Corpus of Contemporary American English (COCA) (n=216,999 sentences; [18]).

We obtained LLM embeddings from GPT2-XL for each sentence in these corpora. Each corpus was partitioned into chunks of 5,000 sentences each (or fewer, if a corpus contained fewer than 5,000 sentences) due to the computational memory load of obtaining LLM embeddings for large numbers of sentences. The corpora were partitioned into a total of 395 chunks. We used our encoding model to predict the left hemisphere language network's response to each sentence in each corpus chunk. For each chunk, we stored the 10 sentences that were predicted to elicit the highest language network response out of all the sentences in that chunk. Similarly, we stored the 10 sentences that were predicted to elicit the lowest language network response. Across the 395 corpus chunks, we hence obtained 10 sentences * 2 * 395 chunks = 7,900 candidate sentences (3,950 *drive* and 3,950 *suppress* sentences).

##### SI 9B: Selecting the final sets from the candidate materials

To identify a set of 250/250 *drive* and *suppress* sentences (500 total), we followed a five-step procedure.
First, we filtered the sentences according to the automatic exclusion criteria reported in SI 2 (to ensure that the candidate sentences did not consist of e.g., exclusively numerical characters or contained excessive punctuation). The automatic exclusion criteria excluded 220 sentences out of 7,900 (213 for *drive*, 7 for *suppress*), leaving us with 7,680 sentences (3,737 for *drive*, 3,943 for *suppress*). Second, we removed duplicate sentences, which excluded 325 additional sentences (55 for *drive*, 270 for *suppress*), leaving us with 7,355 sentences (3,682 for *drive*, 3,673 for *suppress*). Third, out of these candidate 3,682 *drive* sentences and 3,673 *suppress* sentences, we selected the *drive* sentences above the 50th percentile of predicted language network response (out of 3,682 *drive* sentences) and selected the *suppress* sentences below the 50th percentile of predicted language network response (out of 3,673 for *suppress* sentences), effectively filtering out half of our candidate sentences. Hence, for *drive* sentences the percentile exclusion left us with 1,841 sentences (the 50th percentile was 0.497). For *suppress* sentences, the percentile exclusion left us with 1,836 sentences (the 50th percentile was -0.341). Fourth, we manually marked sentences for exclusion according to the manual exclusion criteria reported in SI 2 (to ensure that the candidate sentences were not e.g., inappropriate/offensive or fully in a foreign language). As a result, 838 *drive* sentences and 0 *suppress* sentences were excluded, leaving 1,003 *drive* candidate sentences and 1,836 *suppress* candidate sentences respectively. However, for the *suppress* candidate sentences, 6 sentences occurred in the *baseline set*, and these were excluded, leaving us with 1,830 *suppress* candidate sentences. Fifth and finally, to make our sentence selection independent of human judgment, we randomly sampled 250 sentences from each set (*drive* and *suppress*), leaving us with the final set of 500 *drive/suppress* sentences from the *search* approach.

| **Corpus** | **Number of sentences** | **Contribution (%)** |
|---|---|---|
| **Total** | **1,871,413 (unique: 1,805,623)** | **100%** |
| **Written:** Toronto Adventure | 62,661 | 3.35% |
| **Written:** Toronto Fantasy | 488,841 | 26.12% |
| **Written:** Toronto Mystery | 177,523 | 9.49% |
| **Written:** Wall Street Journal | 2,997 | 0.16% |
| **Written:** Brown | 1,421 | 0.08% |
| **Written:** Universal Dependencies | 1,063 | 0.06% |
| **Written:** Contract Corpus | 264 | 0.01% |

| | | |
|---|---|---|
| **Written:** CRAFT | 34 | 0.00% |
| **Web:** Common Crawl C4 | 878,552 | 46.95% |
| **Spoken:** Cornell Movie Dialogs | 41,058 | 2.19% |
| **Spoken**: COCA | 216,999 | 11.58% |

**SI Table 9.** We identified *drive* and *suppress* sentences by using our encoding model to generate predicted LH language network responses to 1,805,623 unique sentences from diverse, large text corpora. The table contains information on corpus name (as well as main category: published written text, web media text, or transcribed spoken text), number of sentences, and the percent contribution of the total set.

#### SI 10: Breakdown of text sources for *drive*, *suppress*, and *baseline* sentences

To test whether and how sentences differ across the three conditions (*drive*, *suppress*, and *baseline*) and from commonly used stimuli sources in past work, we performed two analyses. First, we analyzed the distribution of text sources across our three conditions. In general, our sentences came from four types of sources: i) written fiction, ii) written miscellaneous, iii) spoken, and iv) web. The “Written Fiction” category consists of fiction narratives from the Toronto Book Corpus [10] and most resembles the materials from prior investigations with naturalistic, fiction materials (e.g., [148–50]). The “Written Misc” category consists of texts from the Brown Corpus [11] (different text genres, all written text), the Wall Street Journal Corpus [12] (newswire), the Universal Dependencies Corpus (different text genres, all written text), the Contract Corpus (legal texts), and the CRAFT Corpus (biomedical articles). The “Spoken” category consists of spoken language from the Cornell Movie Dialogs corpus [17] (conversations from raw movie transcripts) and the spoken component of the Corpus of Contemporary American English [18] (unscripted conversations from TV and radio programs). Finally, the “Web” category consists of text scraped from the web [16] (C4 Common Crawl). **SI Figure 10A** shows the number of sentences from each of the four text sources for the *drive*, *suppress*, and *baseline* sentences. Although a large fraction of stimuli in the *baseline* and *suppress* sets come from the Written Fiction category, the *drive* set contains hardly any such sentences and is instead dominated by the Web category. Moreover, sentences from the Web sources span a larger range of surprisal values (demonstrated to modulate brain responses in Figure 5 in the main text) than the other text sources (**SI Figure 10B**). We note that the breakdown of text sources into these four categories is not meant to suggest that these are the main categories within “naturalistic” stimuli more broadly. Rather, our analysis aims to provide an intuition for what kinds of corpora the *drive*, *suppress*, and *baseline* materials are sampled from. Several prior studies leverage e.g., radio/internet podcasts or other types of storytelling (e.g., [51,52]) which may not nicely fit into the four categories investigated here.

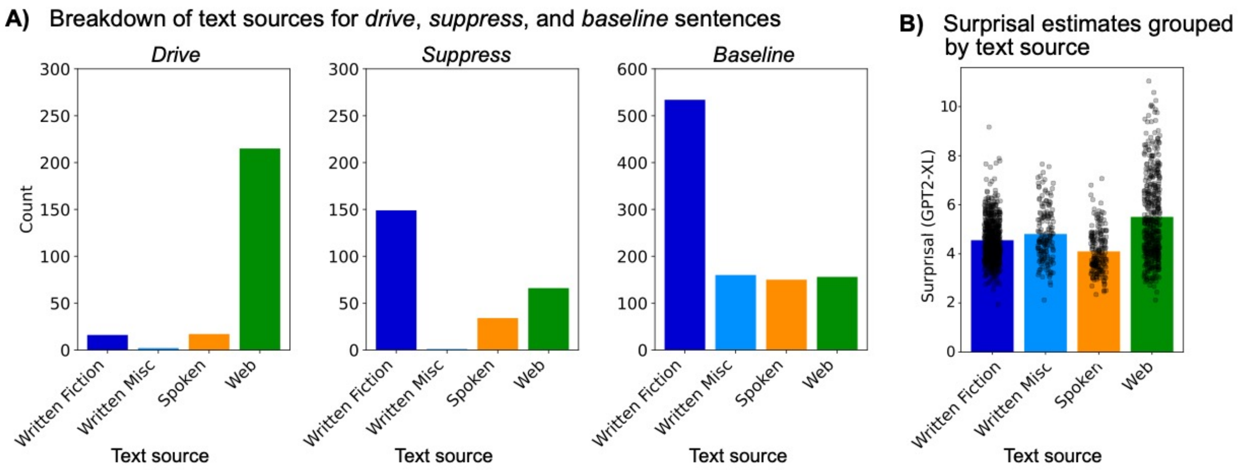

**SI Figure 10. *Drive* and *suppress* sentences are largely from different types of text sources.**
**A)** We quantified which text sources the *drive*/*suppress*/*baseline* sentences came from. The bars show the number of sentences from each text source across the *drive*, *suppress*, and *baseline* sentences separately.
**B)** We quantified the surprisal values across the total of n=1,500 *drive*/*suppress*/*baseline* grouped by text source. The bars show the average surprisal values (estimated by GPT2-XL; note that surprisal is the negative log probability reported in Figure 5 in the main text) for the *baseline/drive/suppress* materials grouped by text source, with scatter points showing average surprisal estimates for individual sentences.

#### SI 11: Quantification of n-gram overlaps between the *baseline* set and *drive/suppress* set

In order to examine how much generalization beyond the training (*baseline*) set is required for predicting neural responses to the *drive* and *suppress* sentences, we investigated the extent of overlap between the *drive/suppress* sentences (n=500 sentences derived from the main *search* approach) and the *baseline* sentences. Moreover, we quantified, on a sentence-by-sentence level, whether greater overlap with the *baseline* set was associated with better predictions. To quantify n-gram overlap, we lower-cased all sentences, stripped punctuation and counted the number of unique n-grams across the *baseline* set (n=1,000 sentences). There were 2,310 unique individual words, 4,415 unique bigrams, 3,951 trigrams, 2,996 4-grams, 1,999 5-grams, and finally 1,000 6-grams (as there were no duplicate sentences in the set, and the sentences are 6 words long). First, we note that no sentences within the *drive/suppress* set (n=500 *drive/suppress* sentences derived from the main *search* approach) had any 5- or 6-gram overlaps with the *baseline* set. Only 1 of the 500 sentences (0.2%) had a 4-gram overlap with the *baseline* set, and only 39 sentences (7.8%) had 3-gram overlaps with the *baseline* set. Hence, the overlaps were largely on the individual word level (97.4% sentences had at least one overlapping word with the *baseline* set) or on the bigram level (50.0% sentences had at least one bigram overlap with the *baseline* set), which suggests that the *drive/suppress* sentences indeed were quite different from the *baseline* set. The number of unique individual words in the *drive/suppress* set that overlapped with the *baseline* set was 457 (out of 2,310 words, i.e., 19.8%), and the number of unique overlapping bigrams was 198 (out of 4,415 bigrams, i.e., 4.5%). A large fraction of the overlapping bigrams (58.6%) contained exclusively function words (e.g., “are in” or “out to”), again suggesting that the *drive/suppress* sentences were largely distinct from the *baseline* sentences.

Next, we quantified whether n-gram overlap had a systematic effect on the sentence-level predictions of the *drive/suppress* sentences. **SI Figure 11A** shows the observed versus predicted brain responses from the n=3 *evaluation* participants using the main *search* approach, colored according to the n-gram overlap. As noted above and evident from the unigram subplot, the majority of *drive/suppress* sentences (97.4%) have at least one word in common with words that occurred in the *baseline* set. The proportion of overlapping words was generally higher in the *suppress* sentences as evidenced by darker colors (i.e., larger counts; see histogram insets) than in the *drive* sentences, indicating that the *drive* sentences in particular were highly non-overlapping with the *baseline* sentences. As evident from the bigram and trigram subplots in **SI Figure 11A**, the number of bigrams and trigrams was in general lower compared to unigrams, with almost no trigram overlaps (7.8%).

**SI Figure 11B** shows the absolute difference between the encoding model predictions and the observed brain responses versus n-gram overlap. For unigram overlaps, we evidenced a negative correlation: sentences with a larger number of overlapping words with the *baseline* set were in general predicted better ($r = -0.23$, $p = 1.55e-07$). Similarly, sentences with bigram overlap with the *baseline* set were predicted better ($r = -0.16$, $p = 2.16e-04$), whereas that was not the case for sentences with trigram overlaps ($r = -0.04$, $p = 0.32$). Although we do observe negative trends in these analyses, as one might expect (generalization to stimuli that are further away from the training distribution is expected to be more challenging), we note that these trends do not explain much variability in prediction performance (unigrams: $r^2 = 0.053$, bigrams: $r^2 = 0.026$, trigrams: $r^2 = 0.002$).

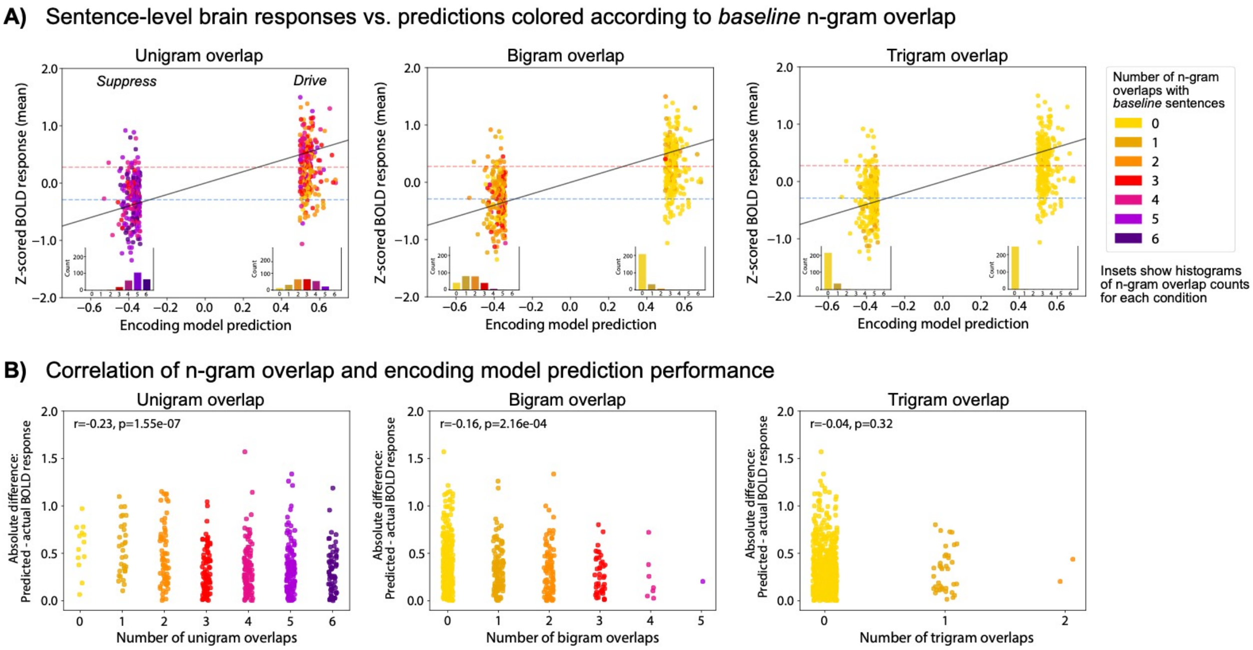

**SI Figure 11. *Drive* (and less so, *suppress*) sentences are largely distinct from the *baseline* sentences, and *drive*/*suppress* sentences that have greater n-gram overlap with *baseline* sentences are predicted slightly better.**
**A)** Sentence-level brain responses (y-axis) as a function of the predicted responses (x-axis) from Figure 3 in the main text, colored according to the n-gram overlap (unigram, bigram, and trigram in separate subplots). A value of 0 means that none of the words in the sentence occurred in the *baseline* set, while a value of 6 means that every single word in the sentence occurred in the *baseline* set. Histogram insets show the distribution of the number of n-gram overlaps for the *drive* and *suppress* conditions separately. Dashed horizontal lines show the mean of the *drive* and *suppress* conditions.
**B)** The absolute difference between the encoding model predictions and the observed brain responses (y-axis) as a function of the number of n-gram overlaps (x-axis; unigram, bigram, and trigram in separate subplots). Individual data points are sentences.

#### SI 12: Condition-level brain responses for the LH language network (not normalized)

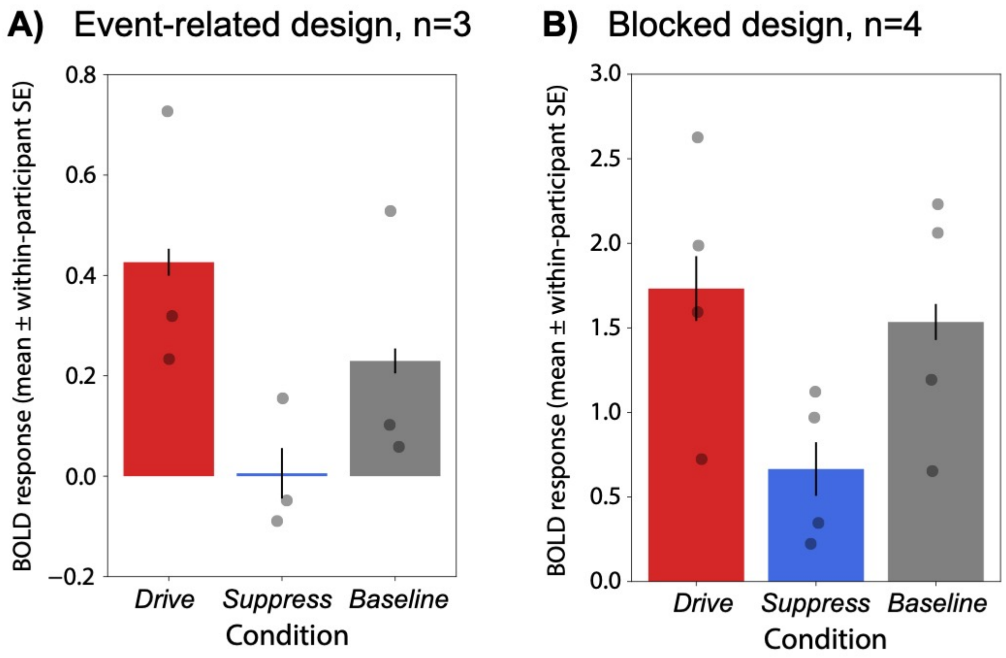

**SI Figure 12. Condition-level responses for the LH language network (not normalized; this figure mirrors Figure 2 in the main text).**
**A**) The mean LH language network fMRI response (not z-scored) across respectively 250 *drive,* 250 *suppress*, and 1,000 *baseline* sentences for n=3 *evaluation* participants, collected in an event-related, single-trial fMRI paradigm (Methods; fMRI experiments). In both A and B, individual points show the average of each condition per participant. Error bars show within-participant standard error of the mean.
**B)** The mean LH language network fMRI response (not z-scored) across respectively 240 *drive,* 240 *suppress*, and 240 *baseline* sentences (randomly sampled from the superset of 250/250 *drive/suppress* sentences and 1,000 *baseline* sentences) for n=4 *evaluation* participants, collected in a blocked fMRI design (Methods; fMRI experiments).

### SI 13: Sentence-level brain responses versus predictions for individual participants

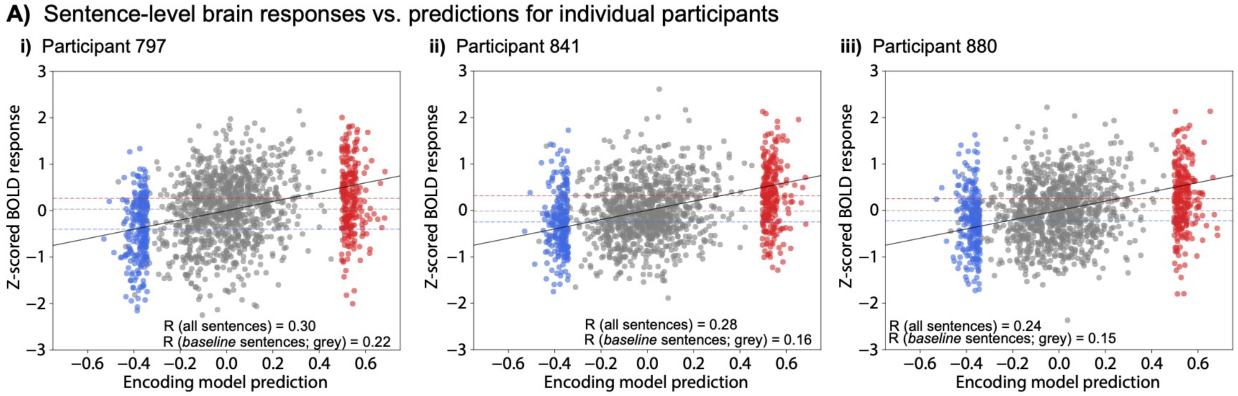

**SI Figure 13. Sentence-level brain responses from each of the three *evaluation* participants versus predicted responses from the encoding model (this figure supplements Figure 3 in the main text).** Predicted brain responses were obtained from the encoding model (x-axis). The observed brain responses (y-axis) are from each of the *evaluation* participant's LH language network (panels i-iii). The blue points represent the *suppress* sentences, the grey points represent the *baseline* sentences, and the red points represent the *drive* sentences. The *suppress* and *drive sentences* were selected to yield respectively low or high brain responses and are therefore clustered on the low and high end of the prediction axis (x-axis). Dashed horizontal lines show the mean of each condition.

#### SI 14: Sentence-level brain responses versus predictions (not normalized)

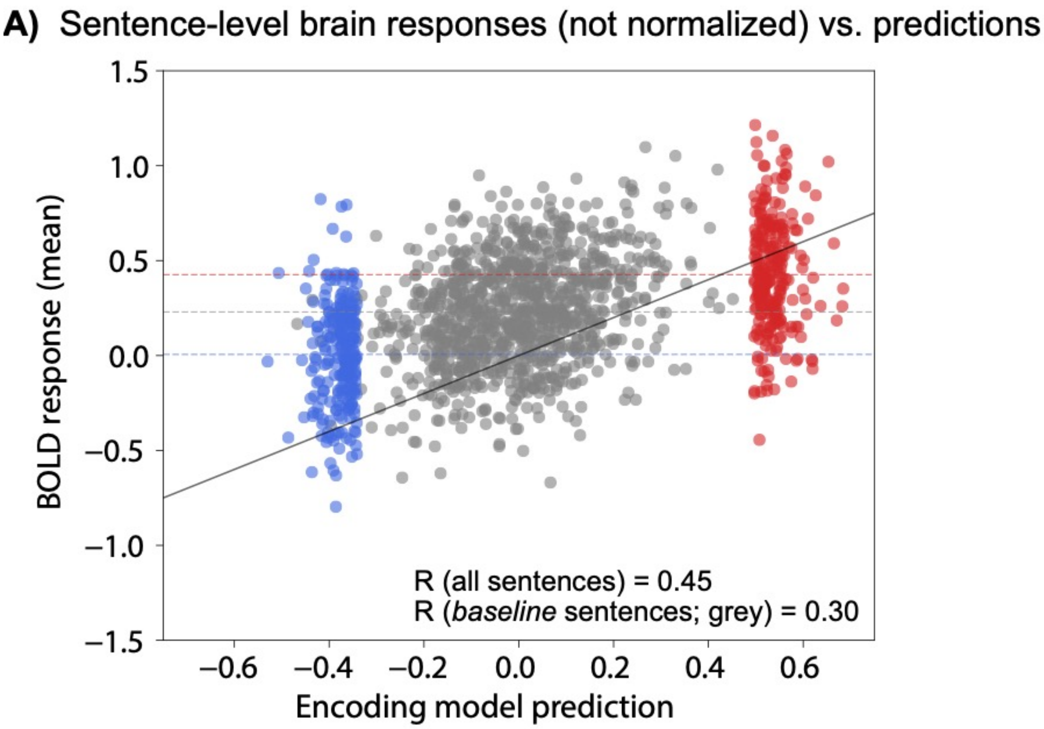

**SI Figure 14. Sentence-level brain responses from the average of three *evaluation* participants versus predicted responses from the encoding model (this figure mirrors Figure 3 in the main text where the responses are z-scored).**

Predicted brain responses were obtained from the encoding model (x-axis). The observed brain responses (y-axis) are the average of n=3 *evaluation* participants' unnormalized LH language network responses (panels i-iii). The blue points represent the *suppress* sentences, the grey points represent the *baseline* sentences, and the red points represent the *drive* sentences. The *suppress* and *drive sentences* were selected to yield respectively low or high brain responses and are therefore clustered on the low and high end of the prediction axis (x-axis). Dashed horizontal lines show the mean of each condition.

#### SI 15: Control brain regions of interest

In addition to language regions, we examined i) two large-scale brain networks linked to high-level cognitive processing—the multiple demand (MD) network [38–42] and the default mode network (DMN) [43–47] which—similar to the language regions—were functionally defined using independent localizer tasks in each participant, and ii) a set of anatomical parcels [36] in an effort to cover the entire cortex.

##### SI 15A: Multiple demand (MD) and default mode network (DMN) localizer task

The task used to localize the domain-general Multiple Demand (MD) network was a spatial working memory task contrasting a harder condition with an easier condition in a standard blocked design with a counterbalanced condition order across runs (e.g., [40,53,54]). The *hard* > *easy* contrast targets brain regions engaged in cognitively demanding tasks. Fedorenko et al. [40] have established that the regions activated by this task are also activated by a wide range of other demanding tasks (see also [55,56]). Note that the reverse contrast, *easy* > *hard*, has been shown to robustly activates default mode network (DMN) regions in prior work using similar tasks and contrasts [57–59].
The *easy* > *hard* contrast targets brain regions which support introspective processes, such as mind wandering, reminiscing about the past, and imagining the future (e.g., [44,60]).

On each trial (8s), participants saw a fixation cross for 500ms, followed by a 3x4 grid within which randomly generated locations were sequentially flashed (1s per flash) two at a time for a total of eight locations (hard condition) or one at a time for a total of four locations (easy condition). Then, participants indicated their memory for these locations in a two-alternative, forced-choice paradigm via a button press (the choices were presented for 1,000 ms, and participants had up to 3s to respond). Feedback, in the form of a green checkmark (correct responses) or a red cross (incorrect responses), was provided for 250ms, with fixation presented for the remainder of the trial. Hard and easy conditions were presented in a standard blocked design (4 trials in a 32s block, 6 blocks per condition per run), counterbalanced condition order across runs. Each run included 4 blocks of fixation (16s each) and lasted a total of 448s. Participants completed 2 runs. Participants were instructed to perform the task to their best ability.

##### SI 15B: Multiple demand (MD) network fROIs

The MD fROIs were defined using the *hard* > *easy* working memory contrast from the MD localizer collected in each participant's first scanning session.
To define the MD fROIs, each individual *hard* > *easy* working memory contrast was intersected with a set of twenty binary parcels (ten in each hemisphere). These parcels were derived from a probabilistic activation overlap map for the same *hard* > *easy* contrast [40,53] in 197 participants using watershed parcellation, as described by Fedorenko et al. [61], and covered extensive portions of the lateral parietal and frontal cortices. Specifically, MD fROIs were defined in each hemisphere in the posterior (*postParietal*), middle (*midParietal*), and anterior (*antParietal*) parietal cortex, precentral gyrus (*Precentral_A_precG*), superior frontal cortex (*supFrontal*),

middle frontal gyrus (*midFrontal*) and its orbital part (*midFrontalOrb*), opercular part of the inferior frontal gyrus (*Precentral_B_IFGop*), the anterior cingulate cortex and pre-supplementary motor cortex (medialFrontal), and the insula (*insula*). These parcels were constrained to be bilaterally symmetric by averaging individual *hard* > *easy* contrast maps across the two hemispheres prior to generating the group-level parcel representation (only the group-based parcels, covering large swaths of cortex, were constrained in this way; fROIs were free to vary in their location across hemispheres, within the borders of these parcels).
Within each of these twenty parcels, the 10% of voxels with the highest t-values for the *hard* > *easy* contrast were selected (see SI Table 15E for number of voxels in each fROI). In the rare cases where the top 10% t-statistic threshold was equal to or less than 0 (meaning that the voxels showed effects in the opposite direction), no voxels were extracted for that given ROI.

#### SI 15C: Default mode network (DMN) fROIs

The DMN fROIs were defined using the reverse contrast (i.e., *easy* > *hard*) from the MD localizer collected in each participant's first scanning session.
To define the DMN fROIs, following Mineroff, Blank et al. [62], each individual *easy* > *hard* contrast was intersected with a set of twelve binary parcels (six in each hemisphere). These parcels were derived from a probabilistic activation overlap map for the same *easy* > *hard* contrast in 197 participants [40,53] using watershed parcellation, as described by Fedorenko et al. [61] and covered extensive portions of the cingulate cortices and lateral frontal and temporal cortices. Specifically, DMN fROIs were defined in each hemisphere in the posterior cingulate cortex (*PostCing*) and middle cingulate cortex (*MidCing*), the temporoparietal junction (*TPJ*), in the medial frontal cortex (*FrontalMed*), the superior temporal gyrus (*STGorInsula*), and anterior temporal gyrus (*AntTemp*). These parcels were constrained to be bilaterally symmetric by averaging individual *hard* > *easy* contrast maps across the two hemispheres prior to generating the group-level parcel representation (only the group-based parcels, covering large swaths of cortex, were constrained in this way; fROIs in the current study were free to vary in their location across hemispheres, within the borders of these parcels).
Within each of these twelve parcels, the top 10% of voxels with the highest t-scores for the *easy* > *hard* contrast were selected (see SI Table 15E for number of voxels in each fROI). In the rare cases where the top 10% t-statistic threshold was equal to or less than 0 (meaning that the voxels showed effects in the opposite direction), no voxels were extracted for that given ROI.

(Because of small amounts of overlapping voxels between the language, MD, and DMN networks (0.12%, on average between the language and MD networks, and 0.35%, on average, between the language and the DMN), if a voxel was selected as belonging to both the language and the MD network, it was assigned to the language network; similarly, if a voxel was selected as belonging to the language network and the DMN, it was assigned to the language network.)

#### SI 15D: Anatomical Glasser parcels

To supplement the functionally defined ROIs, we included anatomical parcels from the Glasser parcellation derived from multi-modal data in the Human Connectome Project [36]. This parcellation contains 360 parcels in total (180 in each hemisphere). Because our functional MRI sequence did not cover the entire brain for some of the participants, we could not extract responses from the full set of 360 Glasser parcels for all participants. On average, responses were extracted from 353.71 parcels (std: 10.34) across n=14 participants (5 *train* participants, 5 *evaluation* participants in the event-related fMRI design from the *search* and *modify* approaches, and 4 *evaluation* participants in the blocked fMRI design).

#### SI 15E: Number of voxels in each (f)ROI

| ***Network*** | ***ROI*** | ***Mean voxels*** | ***Median voxels*** | ***Std voxels*** |
|---|---|---|---|---|
| Language | Lang LH Network | 615.86 | 617 | 4.28 |
| Language | Lang LH IFGorb | 36.29 | 37 | 2.67 |
| Language | Lang LH IFG | 74.93 | 75 | 0.27 |
| Language | Lang LH MFG | 47.0 | 47 | 0.0 |
| Language | Lang LH AntTemp | 162.93 | 163 | 0.27 |
| Language | Lang LH PostTemp | 294.71 | 295 | 1.07 |
| Language | Lang RH Network | 615.43 | 617 | 5.88 |
| Language | Lang RH IFGorb | 36.71 | 37 | 1.07 |
| Language | Lang RH IFG | 75.0 | 75 | 0.0 |
| Language | Lang RH MFG | 47.0 | 47 | 0.0 |
| Language | Lang RH AntTemp | 162.21 | 163 | 2.94 |
| Language | Lang RH PostTemp | 294.5 | 295 | 1.87 |
| MD | MD LH Network | 1477.29 | 1526 | 141.85 |
| MD | MD LH postParietal | 394.92 | 396 | 1.66 |
| MD | MD LH midParietal | 86.92 | 87 | 0.29 |
| MD | MD LH antParietal | 200.86 | 201 | 0.36 |
| MD | MD LH supFrontal | 179.79 | 180 | 0.8 |
| MD | MD LH PrecentralAprecG | 117.71 | 118 | 0.83 |
| MD | MD LH PrecentralBIFGop | 63.21 | 64 | 2.67 |
| MD | MD LH midFrontal | 159.86 | 160 | 0.36 |
| MD | MD LH insula | 52.64 | 56 | 12.75 |
| MD | MD LH medialFrontal | 110.64 | 112 | 2.21 |
| MD | MD RH Network | 1483.71 | 1532 | 142.44 |
| MD | MD RH postParietal | 394.31 | 396 | 2.9 |
| MD | MD RH midParietal | 86.92 | 87 | 0.29 |
| MD | MD RH antParietal | 201.0 | 201 | 0.0 |
| MD | MD RH supFrontal | 179.86 | 180 | 0.36 |
| MD | MD RH PrecentralAprecG | 116.07 | 118 | 4.91 |
| MD | MD RH PrecentralBIFGop | 62.43 | 64 | 2.95 |
| MD | MD RH midFrontal | 159.79 | 160 | 0.58 |
| MD | MD RH midFrontalOrb | 163.0 | 163 | 0.0 |
| MD | MD RH insula | 55.46 | 64 | 15.27 |
| MD | MD RH medialFrontal | 109.43 | 111 | 4.62 |
| DMN | DMN LH Network | 538.64 | 542 | 16.36 |
| DMN | DMN LH FrontalMed | 334.0 | 335 | 2.94 |
| DMN | DMN LH PostCing | 68.29 | 72 | 8.34 |
| DMN | DMN LH TPJ | 65.21 | 67 | 3.64 |
| DMN | DMN LH MidCing | 15.86 | 18 | 4.69 |
| DMN | DMN LH STGorInsula | 32.0 | 32 | 0.0 |

| | | | | |
|---|---|---|---|---|
| DMN | DMN LH AntTemp | 23.29 | 26 | 6.24 |
| DMN | DMN RH Network | 540.71 | 544 | 12.69 |
| DMN | DMN RH FrontalMed | 333.36 | 334 | 2.13 |
| DMN | DMN RH PostCing | 68.64 | 72 | 7.09 |
| DMN | DMN RH TPJ | 64.71 | 67 | 4.55 |
| DMN | DMN RH MidCing | 17.07 | 18 | 2.37 |
| DMN | DMN RH STGorInsula | 32.0 | 32 | 0.0 |
| DMN | DMN RH AntTemp | 24.93 | 26 | 2.3 |
| Glasser | Glasser LH LangNetw | 4821.36 | 4883 | 195.75 |
| Glasser | Glasser RH LangNetw | 4998.21 | 5028 | 168.4 |

**SI Table 15E.** The table shows the mean/median/standard deviation number of voxels in each fROI for the language network, the multiple demand (MD) network, the default mode network (DMN) as well as the anatomically defined language network (Glasser) across n=14 participants in the study.

#### SI 15F: Condition-level brain responses for the five left-hemisphere language fROIs

**F)** Condition-level responses for individual LH language fROIs
Event-related design, n=3 (*search* approach)

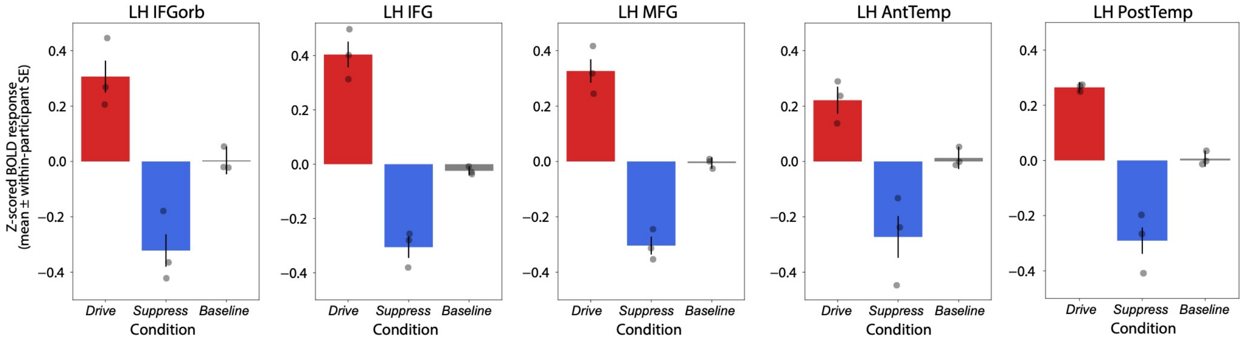

**SI Figure 15F. Condition-level brain responses for individual left-hemisphere (LH) fROIs**
The mean BOLD response (z-scored) across respectively 250 *drive,* 250 *suppress*, and 1,000 *baseline* sentences for n=3 *evaluation* participants, collected in the event-related, single-trial fMRI paradigm. The plots complement Figure 2 in the main text for the full LH language network, and all plots are shown on the same y-axis limits (-0.5, 0.5) except for LH IFG which is plotted on a slightly different y-axis upper bound (0.52 instead of 0.5). Individual points show the average of each condition per participant. Error bars show within-participant standard error of the mean.

SI 15G: Condition-level brain responses for three large-scale brain networks

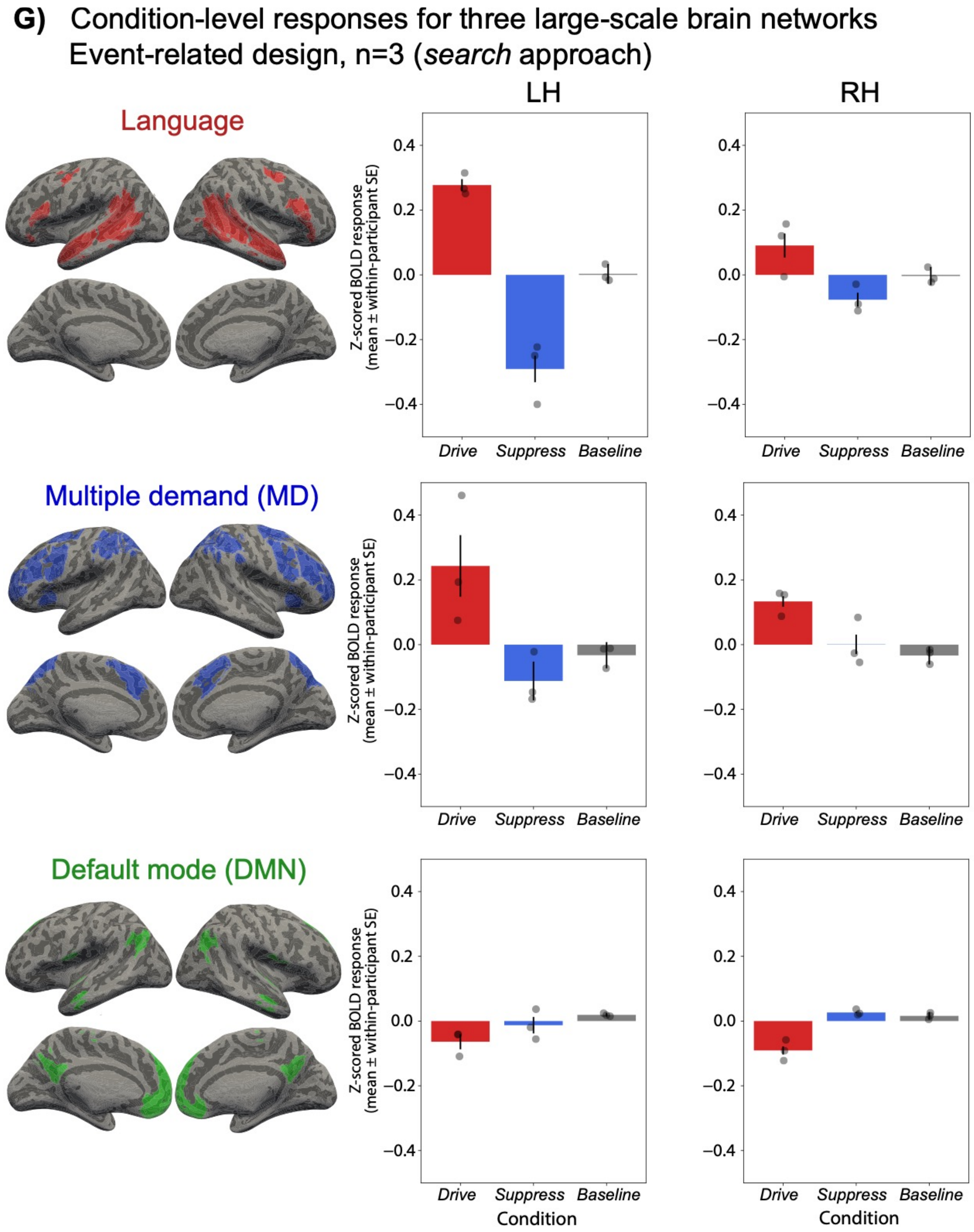

**SI Figure 15G. Condition-level brain responses for language, MD, and DMN networks (both hemispheres).**

The mean BOLD response (z-scored) across respectively 250 *drive,* 250 *suppress*, and 1,000 *baseline* sentences for n=3 *evaluation* participants, collected in the event-related, single-trial fMRI paradigm. Individual points show the average of each condition per participant. Error bars show within-participant standard error of the mean. The brain illustrations show the anatomical parcels (demarcations) that were used to constrain the participant-specific activations for each network in individual participants.

#### SI 16: Driving and suppressing brain responses using the *modify* approach

In an exploratory part of this study, we evaluated whether it is possible to drive and suppress brain responses with completely novel strings (that do not exist in corpora). To do so, we developed a gradient-based *modify* method, in which a subset of words in a random 'seed' sentence are replaced with words such that the modified sentence maximizes or minimizes the predicted language network response. We deliberately avoided constraining the *modify* algorithm to only generate strings that are sensible and grammatically well-formed. As a result, the generated strings often resemble lists of unconnected words (see SI Table 16B and 16C for examples). Similar to the *search* approach, we generated 250 *drive* sentences and 250 *suppress* sentences. We collected brain responses to these 500 *drive*/*suppress* sentences interspersed among the *baseline* sentences in an event-related design in two new participants (3 sessions each, n=6 sessions total) (**SI Figure 16**).

We first describe the sentence generation procedure (SI 16A) followed by evaluation of the recorded brain responses (SI 16B).

##### SI 16A: Generating sentences using the *modify* algorithm

###### The *modify* algorithm

The approach starts with a random *seed* sentence. The goal is to find an optimal modification to this sentence such that the encoding model (Methods; Encoding model development) would predict the modified sentence to yield either a high (*drive*) or a low (*suppress*) response in the language network. Modifying a sentence in this context involves replacing one or more words. For example, a *seed* sentence "Running slow makes me very happy" could be turned into a modified sentence "Running car makes me dull happy" by replacing two words (slow→car and very→dull).

**Input:** Random $\mathcal{S} = \{\boldsymbol{x}_i\}_{i=1}^{n}$, model $\mathcal{M} = \boldsymbol{\theta}_{\text{map}} \circ \boldsymbol{\theta}_{\text{LLM}}$; Learning rate $\alpha$; Loss function $\ell$; Modification iterations $N$; Number of word sites to modify $k$
▷ **Word site selection**
$\mathcal{T} = \text{OrderByImportance}(\{\boldsymbol{x}_i\}_{i=1}^{n})$
▷ From Wallace et. al. (2019)

▷ **Word site modification**
$\boldsymbol{u} = \boldsymbol{x}$
**for** $j$ in $N$ **do**
    **for** $\boldsymbol{x}_i$ in $\mathcal{T}$ **do**
        **if** $k > 0$ **then**
            $\boldsymbol{u}_i^{\text{soft}} = \text{softmax}(\boldsymbol{u}_i)$
            $\boldsymbol{u}_i = \text{Multinomial}(\boldsymbol{u}_i^{\text{soft}})$
            $k = k - 1$
        **end if**
    **end for**
    $y^{\text{pred}} = \mathcal{M}(\boldsymbol{u})$ ▷ Forward pass
    $\nabla = \frac{\partial}{\partial \boldsymbol{u}} \ell(y^{\text{pred}})$ ▷ Backward pass
    $\boldsymbol{u} = \boldsymbol{u} - \alpha.\nabla$
**end for**
$\mathcal{S}^{\text{gen}} = \boldsymbol{u}$
**return** $\mathcal{S}^{\text{gen}}$

**SI Table 16A. Pseudo-code for the *modify* optimization algorithm.**
The pseudo-code is referenced in the text.

Details of the algorithm:

- Decisions about ***how many words*** to replace:

We constrained the number of modifications made to any given *seed* sentence. A larger number of modifications per sentence allows for greater modulation of the sentence's overall surprisal (as estimated by an LLM) and thus can facilitate achieving some target predicted level of brain response, but it is computationally costly. In our initial experiments on a small subset of *seed* sentences, we observed that *suppress* sentences were more difficult to generate than *drive* sentences. We therefore allowed for more modifications for *suppress* sentences (between 1 and 5 word replacements; cf. between 1 and 4 word replacements for the *drive* sentences).
The number of allowed replacements is provided as the input parameter *k* to the algorithm (line 1 in **SI Table 16A**).

- Decisions about ***which words*** to replace (line 3 in **SI Table 16A**):

For a given number of replacements (*k*), we select the *k* words whose replacements modulate the predicted brain responses the most, as determined using the method of integrated gradients [70,72]. This method ranks the words in the *seed* sentence by their sensitivity to the encoding model's predictions. We pick the top-*k* words from the ranked list.
For simplicity, we do not modify words in the *seed* sentence that map to multiple byte-pair tokens. Additionally, if the first word of a *seed* sentence was selected to be modified, we constrain it to be capitalized so as to match the format of sentences from the *baseline set* (i.e., selecting only from capitalized words in the vocabulary).

- Decisions about ***which words to use as replacements*** (lines 6-20 in **SI Table 16A**):

We frame the optimal modification of a sentence as a problem in gradient-based search. Broadly, the algorithm seeks to modify the *seed* sentence (by replacing the top-*k* number of words) such that the predictions of the encoding model for the modified sentence optimize the loss function (elaborated below).

Each word in the *seed* sentence is represented using a vector where each index corresponds to a candidate replacement word. Our candidate replacement words consist of the 50,257 words from the GPT2-XL vocabulary, yielding a vector of size 50,257 for each of the top-*k* words that are to be replaced. If the word "slow" is to be replaced, the index in the vector corresponding to "slow" will have a value of 1, and 0 in all other indices before the optimization pass, meaning that the vector represents the word "slow" with probability 1. During the optimization pass, this vector is iteratively modified to identify candidate replacement words. For example, after one optimization pass, the vector might result in a value of 0.3 in the index corresponding to "slow" and a value of 0.7 in the index for "car", which means that the word "car" is likely to be a good replacement candidate to meet the objective (i.e., achieve a particular value of the predicted brain response for the sentence; see <u>Decisions about goal objectives</u> below). Hence, after the optimization pass (lines 16-18 in **SI Table 16A)**, the vector for each of the top-*k* words contains a distribution of values (higher values mean a replacement that is more likely to lead to the sentence achieving the desired objective). The optimizer modifies all the top-k words simultaneously in a given iteration (i.e., jointly optimizes the vectors corresponding to the top-*k* word sites). In the optimization process, we chose the following mean squared error loss function: (*goal objective – predicted response on the modified sentence*)$^2$. We discuss how we establish *goal objective* in the following section.

The optimization algorithm operates on the continuous-valued vector representations of the top-*k* words in the *seed* sentence, and modifies these representations*.* Selecting a specific replacement word from the resulting optimized vector (the discretizing step; lines 11-12 in **SI Table 16A**) is typically done by simply selecting the vector index with the highest value (e.g., [63,70]). We instead employ multinomial sampling, which samples a word index assuming that the distribution of index values is multinomial. The word index that gets sampled the most times across multiple samples (20-25 samples; see Additional hyperparameters) is selected as the replacement. Multinomial sampling has been shown to provide better estimates than selecting the highest value [71].

In summary, the optimization algorithm identifies optimal replacements for the top-*k* words in the sentence, resulting in a new sentence *u* (line 18 in **SI Table 16A**). We ran the optimization algorithm N times (an input parameter to the algorithm; see Additional hyperparameters) for each *seed* sentence. The modified sentence obtained in one iteration of the algorithm is the input sentence to the next iteration. The modified sentence from the final iteration of the algorithm is the sentence we consider for our experiments (line 20 in **SI Table 16A**).

- Decisions about the ***goal objectives***:

The range of brain response values predicted by the encoding model on the *baseline set* (used to train the encoding model) was in the range [-0.47, +0.54]. To create strings whose predictions would go beyond the upper bounds of the positive and negative responses observed for the *baseline set*, we selected our desired prediction goals to be +1.2 for the *drive* set and -0.8 for the *suppress* set. These values were selected after experimenting with a wider range of goal values and observing that for more extreme values like +5 and -5, the algorithm failed to generate strings with the desired predictions.

- Additional **hyperparameters**:

Aside from i) the number of words (*k*) to replace in a *seed* sentence (between 1 and 4 for the drive stimuli, and between 1 and 5 for the suppress stimuli, as noted above) and ii) the goal objectives (+1.2 for drive and -0.8 for suppress, as noted above), the algorithm's hyperparameters included:

iii) N: the number of iterations that the optimization algorithm is run;
iv) α: the learning rate of the optimizer;
v) the number of multinomial samples drawn in order to select the replacement word.

Empirically, it has been found that assigning larger hyperparameter values in the initial few iterations, followed by smaller values in the later iterations, is effective at finding optimal solutions [73]. Consequently, we ran the algorithm in two rounds for each sentence, where the output string from round 1 was the input string to round 2. We selected hyperparameters that allowed faster convergence towards a solution in the first round, and smaller values in the second round. The hyperparameter values (for hyperparameters not already discussed) were as follows:

- N: round 1: 40; round 2: 60
- α: round 1: 0.01; round 2: 0.005
- number of multinomial samples: round 1: 20; round 2; 25

All sentences were run through two rounds of the algorithm, except for *suppress* sentences that had a predicted brain response above 0.25 after round 1. In our initial experiments, we found that the algorithm was unable of lowering the predictions substantially if the starting point brain response prediction was greater than 0.3, and we therefore decided to save on computation by only running the *suppress* sentences with the lowest predicted brain responses through round 2 of the algorithm.

For details on the *modify* algorithm, see https://github.com/ALFA-group/GOLI for code, **SI Table 16A** for pseudocode; and [63–70] for related approaches.

Selecting the final sets from the candidate materials
We started with 1,500 *seed* sentences. These 1,500 *seed* sentences were obtained from the same 11 corpora as used for *search* material procedure, see SI 9). From each of the three main text categories (written, web, and spoken), we randomly sampled 1,000 sentences. These 3,000 sentences were filtered according to the exclusion criteria reported in SI 2. Finally, we sampled 500 sentences from each text category post-filtering, leading to the final set of 1,500 *seed* sentences. We performed two rounds of the modification algorithm based on the 1,500 *seed* sentences, and ended up with a total of 5,114 candidate sentences (3,000 *drive* and 2,114 *suppress* sentences).
To identify a set of 250/250 *drive* and *suppress* sentences (500 total), we followed a five-step procedure, which mirrors the procedure used for selecting the *search* materials, with one additional consideration to exclude modified sentences created from the same *seed* sentence. First, we filtered the sentences according to the automatic exclusion criteria reported in SI 2. Additionally, in *modify,* we ensured that we selected only one sentence from the sentences produced in each of the two modification rounds that a *seed* sentence underwent. Among the sentences from the two different modification rounds, we selected the one closest to the desired prediction and which passed the automatic exclusion criteria. This automatic exclusion procedure excluded 3,075 sentences out of 5,114 (1,951 for *drive*, 1,124 for *suppress*), leaving us with 2,039 sentences (1,049 for *drive*, 990 for *suppress*). Second, we checked for duplicate sentences in the set, and none were identified. Third, out of these candidate 1,049 *drive* sentences and 990 *suppress* sentences, we selected *drive* sentences above the 50th percentile of predicted language network response (out of 1,049 *drive* sentences) and selected *suppress* sentences below the 50th percentile of predicted language network response (out of 990 for *suppress* sentences), effectively filtering out half of our candidate sentences. Hence, for *drive* sentences the percentile exclusion left us with 524 sentences (the 50th percentile was 0.565). For *suppress* sentences, the percentile exclusion left us with 495 sentences (the 50th percentile was -0.056). Fourth, we manually marked sentences for exclusion according to the exclusion

criteria reported in SI 2. As a result, 19 *drive* sentences and 4 *suppress* sentences were excluded, leaving 505 *drive* candidate sentences and 491 *suppress* candidate sentences respectively. Fifth and finally, to make our sentence selection independent of human judgment, we randomly sampled 250 sentences from each set (*drive* and *suppress*), leaving us with the final set of 500 *drive/suppress* sentences from the *modify* approach.

| *Seed sentence* | *Pred* | *Sentence round 1* | *Pred* | *Sentence round 2* | *Pred* |
|---|---|---|---|---|---|
| Create holiday cards, gifts and decorations. | -0.06 | Create obesity cards, Advisory and Ok. | 0.72 | Create obesity massacre, false and Ok. | 0.83 |
| White Gates - the quintessential country cottage. | 0.13 | White measles - the quintessential country Had. | 0.65 | White measles - the quintessential Dept Had. | 0.79 |
| Reversing that would be too radical. | 0.06 | Reversing that Revenue Roberts too HUM. | 0.67 | Reversing that Revenue Roberts too EXP. | 0.79 |
| Proximity says it very well could. | 0.12 | Proximity clasp it very Cav resp. | 0.71 | Proximity clasp it very Cumm resp. | 0.78 |
| The staff is hospitable and helpful. | -0.03 | The staff is hospitable and helpful. | -0.03 | Elsa staff is hospitable and IND. | 0.78 |
| They would never have taken this. | 0.11 | They would Azerbaijan subs taken MIT. | 0.61 | ASY would Azerbaijan subs taken MIT. | 0.72 |
| In fact I feel your pain. | 0.12 | In issu Perry feel your Former. | 0.63 | In wars Perry feel your Former. | 0.72 |
| The industry does take that position. | 0.16 | The industry Warp take that def. | 0.61 | The industry obstacles Married that def. | 0.71 |
| The Blue pointed off to town.. | 0.12 | The Blue pointed ritual to Recommend.. | 0.61 | The Blue pointed rematch to Recommend.. | 0.71 |
| It is not going to work. | -0.10 | It is Kat going to IND. | 0.58 | ARK is Kat going to IND. | 0.71 |
| You need to wait more time. | -0.11 | You Mason to glowing more prob. | 0.62 | ODY Mason to glowing more prob. | 0.71 |
| Then why are you so upset? | 0.01 | Then why are you so These? | 0.58 | Thor why are you so These? | 0.71 |
| We have the moment you missed. | 0.12 | We have the Dynasty you recomm. | 0.62 | Vote have the Dynasty you recomm. | 0.70 |
| Is there any other reason, really? | 0.00 | Is SELECT any revised wretched, really? | 0.56 | Wars SELECT any revised wretched, really? | 0.68 |
| All that, plus our newsmaker tonight. | 0.21 | All that, Called our newsmaker pred. | 0.66 | All that, Called our newsmaker pred. | 0.66 |
| This house must not be sold! | -0.04 | This house must not be sold! | -0.04 | USA house must not be ex! | 0.65 |

**SI Table 16B.** Example *drive* sentences generated by the *modify* algorithm from a random *seed* sentence (first column). The output from *modify* round 1 was used as input to *modify* round 2 in an attempt to further optimize the loss objective (in this case, higher predicted brain response values). The predictions after each iteration are shown in the "Pred" columns.

| ***Seed sentence*** | ***Pred*** | ***Sentence round 1*** | ***Pred*** | ***Sentence round 2*** | ***Pred*** |
|---|---|---|---|---|---|
| I ran back to the door. | -0.26 | People ran back to the door. | -0.32 | We ran back to the door. | -0.35 |
| First, we have a big story. | -0.06 | First, we have a big frame. | -0.25 | Look, we have a big frame. | -0.35 |
| He glanced back out the door. | -0.28 | Driver pops back out the door. | -0.24 | It pops back out the door. | -0.30 |
| We have very little time left. | -0.21 | Cos have very little time limit. | -0.11 | They have very little time limit. | -0.29 |
| Right in the middle of town. | -0.09 | You in the middle of town. | -0.16 | There in the middle of town. | -0.27 |
| The keys were in the ignition. | -0.21 | The keys were in the ignition. | -0.21 | Some keys were in the ignition. | -0.26 |
| Marcus watched Havily leave the room. | -0.10 | Child watched Havily leave the room. | -0.13 | They watched Havily leave the room. | -0.26 |
| She answered on the first ring. | 0.08 | She answered on the first ledge. | -0.02 | She stretched on the first ledge. | -0.24 |
| And we had a wonderful lunch. | -0.26 | Launch we had a wonderful lunch. | -0.19 | Loop we had a wonderful view. | -0.24 |
| This time two men got out. | -0.08 | This time two men got alone. | -0.16 | Some time two men got alone. | -0.24 |
| She answered on the first ring. | 0.08 | She answered on the first ledge. | -0.02 | She stretched on the first ledge. | -0.24 |
| I got cut a few times. | -0.06 | Owner got awake a few times. | -0.14 | Del got awake a few times. | -0.24 |
| It's time for us to hunt. | -0.10 | There's time for us to jumper. | -0.16 | There's time for us to strut. | -0.23 |
| She couldn't handle any more confessions. | -0.12 | Every couldn't handle any more signalling. | -0.13 | They couldn't handle any more signalling. | -0.23 |
| Sixteen attractive shoes around the pot. | -0.04 | Sixteen settled shoes around the rubbish. | -0.20 | Sixteen settled shoes around the courtyard. | -0.23 |
| This is quite an interesting task. | -0.14 | Statement is quite an interesting task. | -0.04 | It is quite an interesting task. | -0.22 |
| We could use a new faucet. | -0.10 | Teen could use a new faucet. | -0.08 | They could use a new faucet. | -0.20 |

**SI Table 16C.** Example *suppress* sentences generated by the *modify* algorithm from a random *seed* sentence (first column).

#### SI 16B: Evaluating brain responses to the *drive*/*search* sentences derived using *modify*

**SI Figure 16A** shows the average responses for the n=2 *evaluation participants* for the *drive, suppress* (derived using the *modify* method), and *baseline* sentence conditions. The *drive* sentences yielded significantly higher responses than the *suppress* sentences (β=0.12, t=2.77, p<.05). The *drive* sentences also yielded significantly higher responses than the *baseline* sentences (β=0.20, t=5.57, p<.001) with the evoked BOLD signal being 57% higher for the *drive* condition relative to *baseline* (quantified using non-normalized BOLD responses). The *suppress* sentences did not yield significantly lower responses than the *baseline* sentences (β=0.07, t=2.06, p=0.098), in fact, the evoked BOLD response was 19.5% higher for the *suppress* condition relative to *baseline*. **SI Figure 16B** shows the observed versus predicted brain responses in the left hemisphere language network for the *modify*-based sentences along with the *baseline* sentences. Across the full set of 1,500 *baseline, drive,* and *suppress* sentences, we obtained a Pearson correlation of 0.22 (p<.001) between observed and predicted brain responses. For the 1,000 *baseline* sentences, the correlation was 0.32 (p<.001).

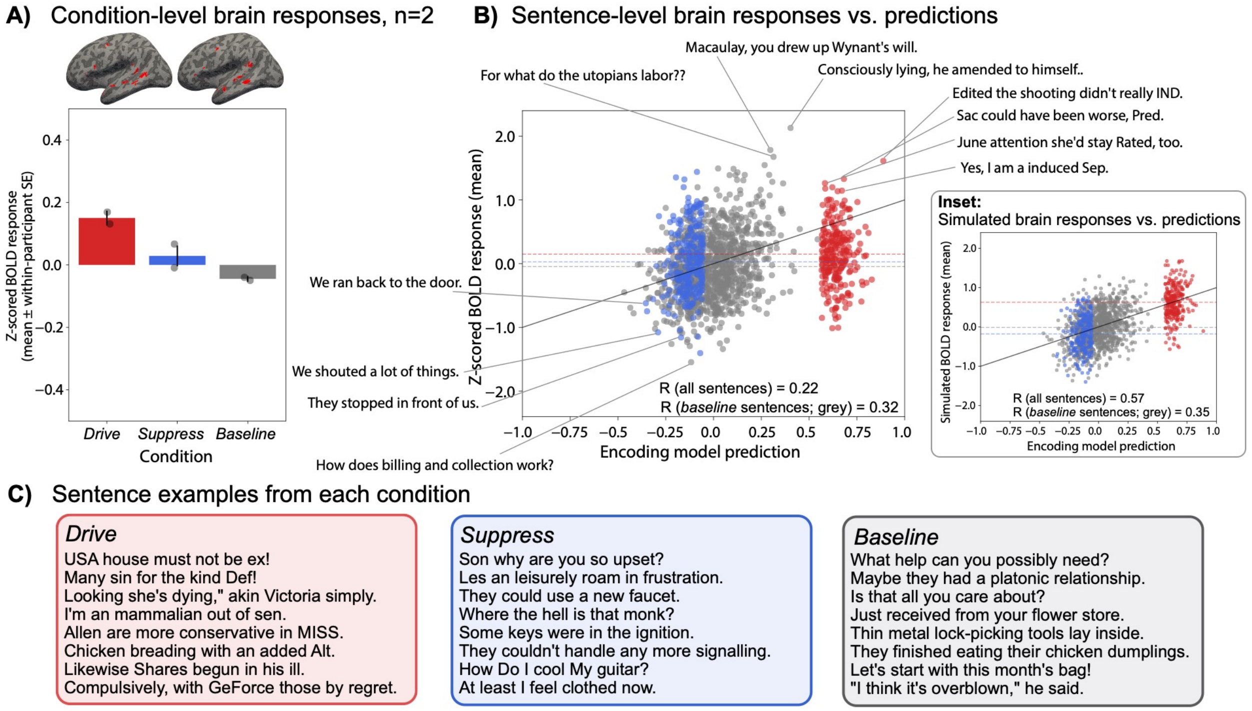

**SI Figure 16. Sentences with word replacements show some ability to drive and suppress the language network but sentence-level predictions are not accurate.**
**A)** The mean LH language network fMRI response across respectively 250 *drive,* 250 *suppress* (derived using the *modify* approach), and 1,000 *baseline* sentences for n=2 *evaluation* participants collected in an event-related, single-trial fMRI design. Individual points show the average of each condition per participant. Error bars show within-participant standard error of the mean. The brain illustrations show the functionally defined language network in the participants of interest on the surface-inflated brain.
**B)** Sentence-level brain responses as a function of the predicted responses along with sentence examples. Predicted brain responses were obtained from the encoding model (x-axis). The true brain responses (y-axis) are the average of n=2 *evaluation* participants (LH language network). The blue points represent the *suppress* sentences, the gray points represent the *baseline* sentences, and the red points

represent the *drive* sentences. Dashed horizontal lines show the mean of each condition.
**Inset:** The simulated brain responses (y-axis) were obtained by sampling from a noise distribution representing the empirical inter-participant variability. This plot illustrates the maximum possible predictive performance, given inter-participant variability and fMRI measurement noise.
**C)** Example sentences from each condition (note that the *baseline set* contains the same materials as in all remaining parts in this study).

To better interpret the accuracy of sentence-level predictions, we quantified the maximal possible prediction performance by treating inter-participant variability as "noise" that cannot be predicted by a computational model (same approach as in Results; Model captures most explainable variance in new participants). According to these calculations, we could have expected to obtain a Pearson correlation of 0.57 across all 1,500 sentences (observed: 0.22, i.e. 38.6% of the theoretically obtainable correlation), and Pearson correlation of 0.35 for the 1,000 *baseline* sentences (observed: 0.32, i.e. 91.4% theoretically obtainable correlation) (**SI Figure 16C)**. Hence, for the *baseline* sentences we evidenced a moderate correlation between observed and predicted brain responses (similar to what we observed for the n=3 *evaluation* participants in the *search* experiment), but the correlation for the full set of sentences, including the *modify*-based *drive*/*suppress* sentences, was lower. Overall, the results from the *modify* approach suggest that novel sentences created using gradient-based word modifications modulated responses in the language network to some degree, but that the encoding model predictions for the individual *drive* and *suppress* sentences were inaccurate. This could be due to (at least) two reasons: i) the resulting strings were often akin to word lists (see SI Table 16B, 16C), which generally elicit a relatively lower response in the language network (e.g., [61,4,74]) (the *modify* sentences were generally rated as ungrammatical and implausible by human participants; SI 19) and ii) word lists were not included as part of the training set for the encoding model.

#### SI 17: Comparison of GPT2-XL encoding model versus surprisal-based encoding models

The encoding model used in this work was based on the hidden states (i.e., unit activations) from the Transformer model GPT2-XL [75] (Methods; Encoding model development). Motivated by the pervasive role of surprisal in accounting for behavioral and neural responses during language processing (e.g., [76–78]; and Figure 5C in the main text), we developed encoding models based on surprisal estimates and evaluated their predictivity relative to our main encoding model. In particular, we obtained surprisal estimates from two surprisal models that predated Transformers: a lexical n-gram model (5-gram; see details in SI 20A) and a probabilistic context-free grammar model (PCFG; see details in SI 20B). Both n-gram and PCFG surprisal have been shown to explain (independent) variance in the language network's neural response during story comprehension [79]. For completeness, we also included surprisal estimates from GPT2-XL (Methods; Sentence properties that modulate brain responses).

First, we evaluated the cross-validated prediction performance of the surprisal models on the *baseline* set (n=1,000 sentences) in a procedure identical to the one reported in SI 6. For reference, the performance of the GPT2-XL hidden states encoding model (i.e., using the representations from the Transformer blocks, specifically block 22, as opposed to the surprisal estimate obtained via a linear layer at the last block) was Pearson r = 0.38 (67.9% of the noise ceiling which was estimated to be r = 0.56, see SI 5). Performance of the n-gram, PCFG, and the GPT2-XL surprisal models was substantially lower: 5-gram surprisal model: r = 0.23 (41.1% of the noise ceiling); PCFG surprisal model: r = 0.15 (26.8% of the noise ceiling), and GPT2-XL surprisal model: r = 0.22 (39.3% of the noise ceiling).

Next, we evaluated how well the surprisal models could account for the data obtained from the n=3 *evaluation* participants on the n=1,500 sentences (the *drive*/*suppress/baseline* sentences). Although this comparison is not completely fair to the surprisal models (given that the stimuli were obtained using the GPT2-XL encoding model), it still provides a proxy for the held-out participant predictivity performance of the surprisal models. **SI Figure 17** shows the observed sentence-level brain responses versus the predictions obtained from the main encoding model (panel A; same as Figure 3 in the main text) vs. the three surprisal models (panels B-D). As evident from the plots and in line with the cross-validated performance evaluation, the surprisal models fall short of the encoding model's performance.

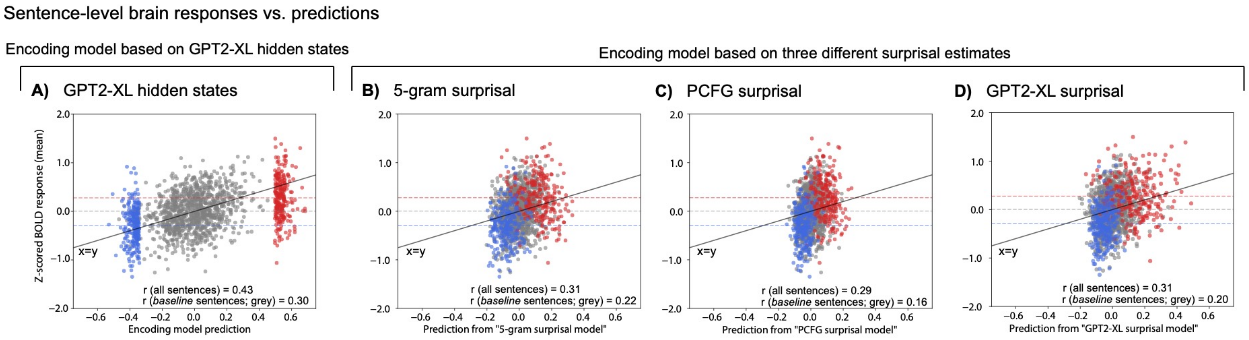

**SI Figure 17. The encoding model based on the GPT2-XL hidden states achieves higher predictivity compared to the encoding models based on univariate sentence surprisal estimates.** We compared the performance of the encoding model (based on GPT2-XL hidden states; **Panel A**) against surprisal estimates from three surprisal models (**panels B-D)** on n=3 *evaluation* participants on the n=1,500 sentences (the *drive*/*suppress/baseline* sentences)**.** The panels show the sentence-level brain responses as a function of the predicted responses and the plots mirror Figure 3 in the main text (same x- and y-axis limits). The only difference is that the predicted brain responses were obtained from the surprisal models in panels B-D.

#### SI 18: Statistical significance of the differences in the BOLD response among sentence conditions

Section SI 18A contains statistics accompanying Results; Model-selected sentences control language network responses” in the main text based on the *search* approach.
Section SI 18B contains statistics accompanying SI 16 based on the *modify* approach.

##### SI 18A: *Search* participants (n=3 participants with 1,500 BOLD responses each)

BOLD response data met assumptions of normality (via Kolmogorov-Smirnov test) and equal variances (via Levene test).

Linear mixed effect formula:

*BOLD response ~ condition + (1 | sentence) + run_within_session + trial_within_run*

| ***Model term*** | ***Coefficient*** | ***SE*** | ***t*** | ***p*** | ***df*** |
|---|---|---|---|---|---|
| *intercept* | 0.073* | 0.029 | 2.569 | 0.01 | 4493 |
| *condition_D* | 0.273*** | 0.028 | 9.734 | <0.001 | 4493 |
| *condition_S* | -0.293*** | 0.028 | -10.448 | <0.001 | 4493 |
| *run_within_session* | -0.003 | 0.003 | -0.941 | 0.347 | 4493 |
| *trial_within_run* | -0.002** | 0.001 | -3.004 | 0.003 | 4493 |
| *SD (Intercept item_id_factor)* | 0.131 | | | | |
| *SD (Observations)* | 0.649 | | | | |
| *Num.Obs.* | 4500 | | | | |
| *R2 Marg.* | 0.060 | | | | |
| *R2 Cond.* | 0.096 | | | | |
| *AIC* | 9112.2 | | | | |
| *BIC* | 9157.1 | | | | |
| *RMSE* | 0.64 | | | | |

*+p < 0.1, *p < 0.05, **p < 0.01, ***p < 0.001*

**SI Table 18Ai**. Effect size estimates, standard error estimates, t-statistics, p-values, degrees of freedom for the LME model as well as model fit statistics (e.g., R2). “condition_D” and “condition_S” denote respectively the *drive* and *suppress* condition (the *baseline* condition was coded in the intercept).

| ***Contrast*** | ***Estimate*** | ***SE*** | ***df*** | ***t.ratio*** | ***p.value*** |
|---|---|---|---|---|---|
| *B - D* | -0.273 | 0.0281 | 1504 | -9.723 | <.0001 |
| *B - S* | 0.293 | 0.0281 | 1504 | 10.435 | <.0001 |
| *D - S* | 0.567 | 0.0356 | 1505 | 15.933 | <.0001 |

**SI Table 18Aii.** Pairwise comparisons of conditions (using estimated marginal means). Degrees-of-freedom method was Kenward-Roger and p-value adjustment was Tukey method for comparing a family of 3 estimates.

#### SI 18B: *Modify* participants (n=2 participants with 1,500 BOLD responses each)

BOLD response data met assumptions of normality (via Kolmogorov-Smirnov test) and there was a marginal effect suggesting unequal variances (via Levene test, p=0.043).

Linear mixed effect formula:

*BOLD response ~ condition + (1 | sentence) + run_within_session + trial_within_run*

| Model term | Coefficient | SE | t | p | df |
|---|---|---|---|---|---|
| *intercept* | 0.003 | 0.035 | 0.086 | 0.932 | 2993 |
| *condition_D* | 0.197*** | 0.035 | 5.575 | <0.001 | 2993 |
| *condition_S* | 0.073* | 0.035 | 2.065 | 0.039 | 2993 |
| *run_within_session* | -0.001 | 0.004 | -0.179 | 0.858 | 2993 |
| *trial_within_run* | -0.002** | 0.001 | -2.072 | 0.038 | 2993 |
| *SD (Intercept item_id_factor)* | 0.246 | | | | |
| *SD (Observations)* | 0.613 | | | | |
| *Num.Obs.* | 3000 | | | | |
| *R2 Marg.* | 0.013 | | | | |
| *R2 Cond.* | 0.150 | | | | |
| *AIC* | 6047.6 | | | | |
| *BIC* | 6089.6 | | | | |
| *RMSE* | 0.57 | | | | |

*+p < 0.1, *p < 0.05, **p < 0.01, ***p < 0.001*

**SI Table 18Bi**. Effect size estimates, standard error estimates, t-statistics, p-values, degrees of freedom for the LME model as well as model fit statistics (e.g., R2). “condition_D” and “condition_S” denote respectively the *drive* and *suppress* condition (the *baseline* condition was coded in the intercept).

| Contrast | Estimate | SE | df | t.ratio | p.value |
|---|---|---|---|---|---|
| *B - D* | -0.1965 | 0.0353 | 1505 | -5.568 | <.0001 |
| *B - S* | -0.0728 | 0.0353 | 1504 | -2.062 | 0.0982 |
| *D - S* | 0.1238 | 0.0447 | 1505 | 2.772 | 0.0156 |

**SI Table 18Bii.** Pairwise comparisons of conditions (using estimated marginal means). Degrees-of-freedom method was Kenward-Roger and p-value adjustment was Tukey method for comparing a family of 3 estimates.

### Sections related to sentence properties that modulate brain responses

#### SI 19: Average values for each sentence property across conditions

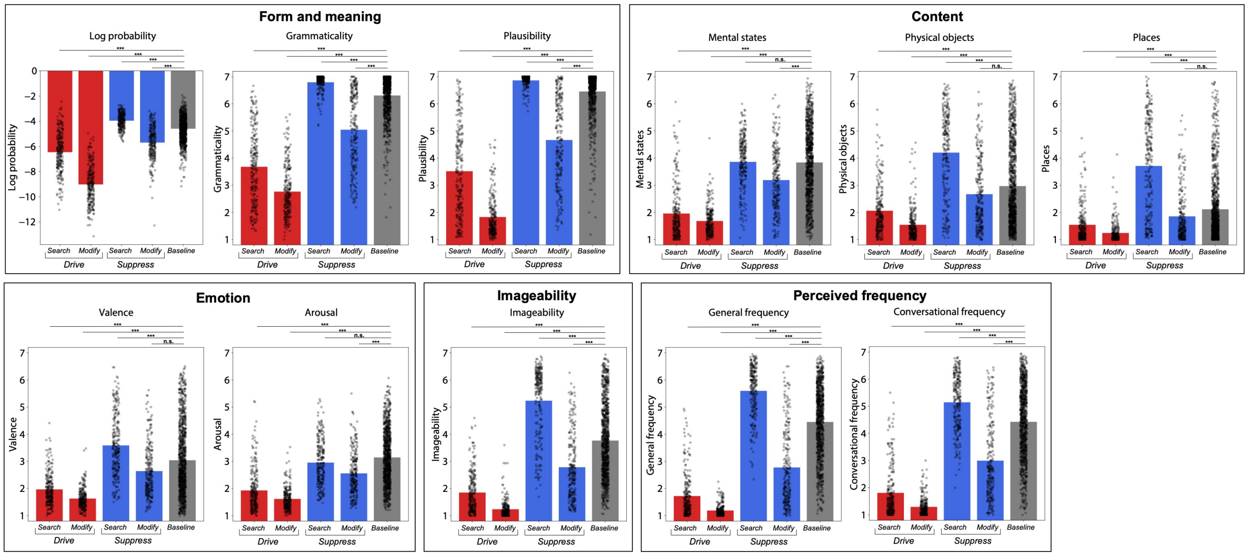

**SI Figure 19. Average values for each sentence property.**
Average values for each sentence property (n=11 sentence properties) are shown across the full set of n=2,000 sentences (1,000 *baseline* sentences, 250/250 *drive/suppress* sentences from the main *search* approach, and finally 250/250 *drive/suppress* sentences from the exploratory *modify* approach). Individual data points show individual sentences. We quantified whether properties for each of the *drive* and *suppress* conditions (from either *search* or *modify*) were significantly different from the *baseline* condition via non-parametric permutation tests. Across 1,000 iterations, we sampled 250 sentences (with replacement) from the *drive*/*suppress* condition and 250 sentences (with replacement) from the *baseline* condition. We generated a null distribution by randomly permuting the condition assignment and measuring the difference between the average of these permuted condition assignment lists. The p-value was obtained by counting the number of times the absolute difference was smaller than the absolute values measured from the permuted data (two-sided test), divided by the number of permutations (n=1,000). P-values were corrected for multiple comparisons (across all 44 comparisons) using the Bonferroni procedure. Significance values are denoted as: p<.001: ***; n.s.: non-significant.

#### SI 20: Brain responses versus surprisal: Additional surprisal estimates

In the main text (**Figure 5**), we estimated surprisal (log probability) across all sentences in the study (n=2,000 sentences) using GPT2-XL. GPT2-XL surprisal is sensitive to all preceding words in the sentence and is trained on massive amounts of diverse text. Here, we included two additional surprisal models to capture distinct information about the predictability of each sentence, specifically: 1) An n-gram model that is sensitive to word co-occurrence patterns of a certain number of preceding words (in our case, four) but limited in its ability to model hierarchical language syntax, 2) A probabilistic context-free grammar (PCFG) model that is sensitive to the syntactic structure of the sentence but does not directly encode word co-occurrence patterns.

##### SI 20A: Lexical n-gram (5-gram) surprisal

Lexical surprisal was computed using a 5-gram language model which estimates surprisal given the preceding four words (i.e., for the first four words there is no such window; the surprisal for the first word of a sentence is the probability of that given word beginning the sentence, for the second word it is the 2-gram probability and so on). The model was trained using the KenLM library ([80]; build on 12/24/2022; https://github.com/kpu/kenlm) with default smoothing parameters (modified Kneser-Ney smoothing) on the training set of Wikitext-103 [81]. The training data were tokenized using the *sent_tokenizer* from the NLTK library ([11]; version 3.8.1) followed by tokenization of sentences using the *wordpunct_tokenizer* from the same library. Punctuation was stripped and all words were lower-cased. The same tokenization and preprocessing procedure was applied to the *drive*/*suppress*/*baseline* materials. Surprisal estimates for each word in the materials was obtained using default parameters of the *full_scores* function in KenLM. Surprisal for each word in the sentence along with an end-of-sentence token (</s>) was computed. 223 unique words from our materials (out of a total of 4,119 unique words, 5.41%) were out of vocabulary for the n-gram model and hence surprisal could not be estimated for these words. We obtained the sentence-level surprisal by taking the mean of the word-level surprisals.

##### SI 20B: Probabilistic context-free grammar (PCFG) surprisal

Lexicalized probabilistic context-free grammar (PCFG) surprisal was computed using the incremental left-corner parser of van Schijndel et al. [82] trained on a generalized categorial grammar [83] reannotation of Wall Street Journal sections 2 through 21 of the Penn Treebank . Each sentence was tokenized using a Penn Treebank Tokenizer and punctuation was retained. We obtained the sentence-level surprisal by taking the mean of the word-level surprisals.

#### SI 20C: Scatterplots of brain responses versus additional surprisal estimates

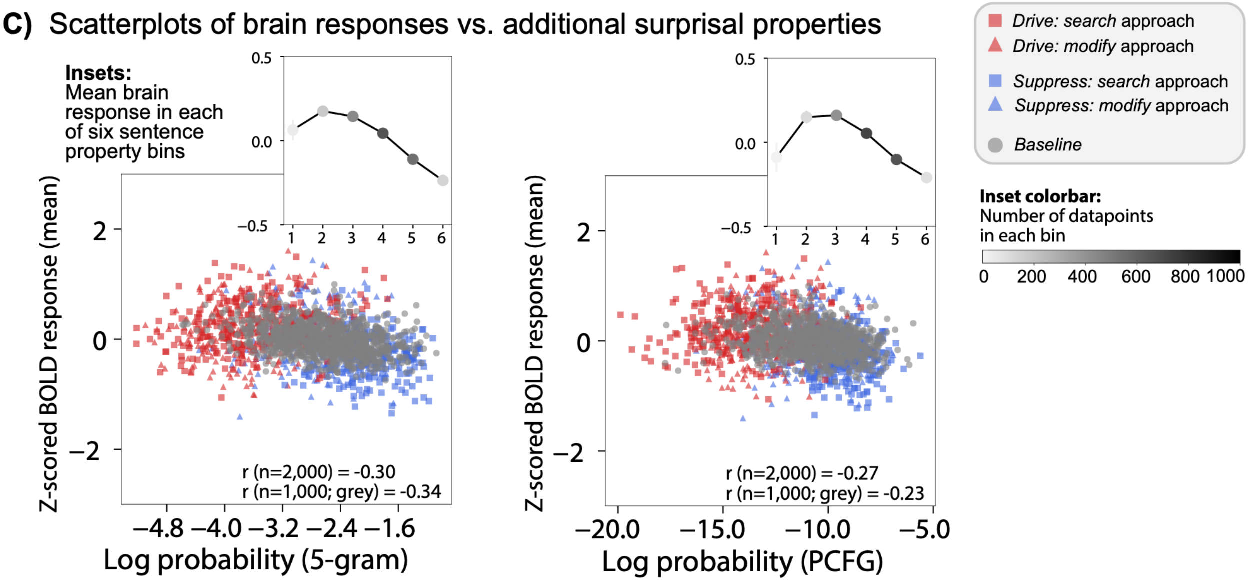

**SI Figure 20C. Sentence-level brain responses as a function of two additional surprisal estimates (supplementary to Figure 5 in the main text).** The brain responses (y-axis) were averaged across n=5 *train* and n=5 *evaluation* participants (250/250 *drive/suppress* sentences from the *search* approach and 250/250 *drive/suppress* from the exploratory *modify* approach as well as 1,000 *baseline* sentences. The insets show the average brain response with each property grouped into six uniformly sized bins. The color of the inset point denotes how many datapoints were available within each bin.

The 5-gram log probability values were correlated with the PCFG log probability values at Pearson r = 0.59. (GPT2-XL log probability was correlated with 5-gram log probability at r = 0.56 and with PCFG log probability at r = 0.42 (on the n=1,000 *baseline* set)).

#### SI 21: Modulation of individual brain regions' responses by sentence properties

To complement the analyses for the whole left hemisphere (LH) language network in the main text (Results; Sentence complexity modulates language network responses), we investigated the modulation of individual brain regions' responses by sentence properties. We here examined i) functionally defined language ROIs (5 in the LH, 5 in the RH; see Methods; Definition of ROIs), and ii) a set of anatomical parcels [36] located in typically language-responsive areas.

First, we quantified the similarity among the n=10 language fROIs in how they are modulated by the different sentence properties for the n=1,000 *baseline* sentences (**SI Figure 21A**). We included the RH fROIs because several lines of work suggest some functional differentiation between the LH and RH language fROIs (e.g., [85,86]). These analyses are similar to those reported in Figure 5A in the main text for the whole LH language network but break down the findings by fROI. As evident from the **SI Figure 21A** (as well as the Figure 4B and 4C in the main text), the LH language fROIs are similar in their sensitivity to different linguistic features (e.g., all fROIs are more sensitive to grammaticality and plausibility than valence). The RH fROIs generally exhibit lower correlations with sentence properties than the LH fROIs. In addition, we observe a clear difference between the LH and RH language fROIs in that the RH language fROIs, but not LH language fROIs, show positive correlations with the "Mental states" and "Arousal" features: sentences with content related to mental states and/or with arousing content elicit higher responses in all RH language fROIs (most pronounced for the RH IFGorb and the RH AntTemp fROIs).

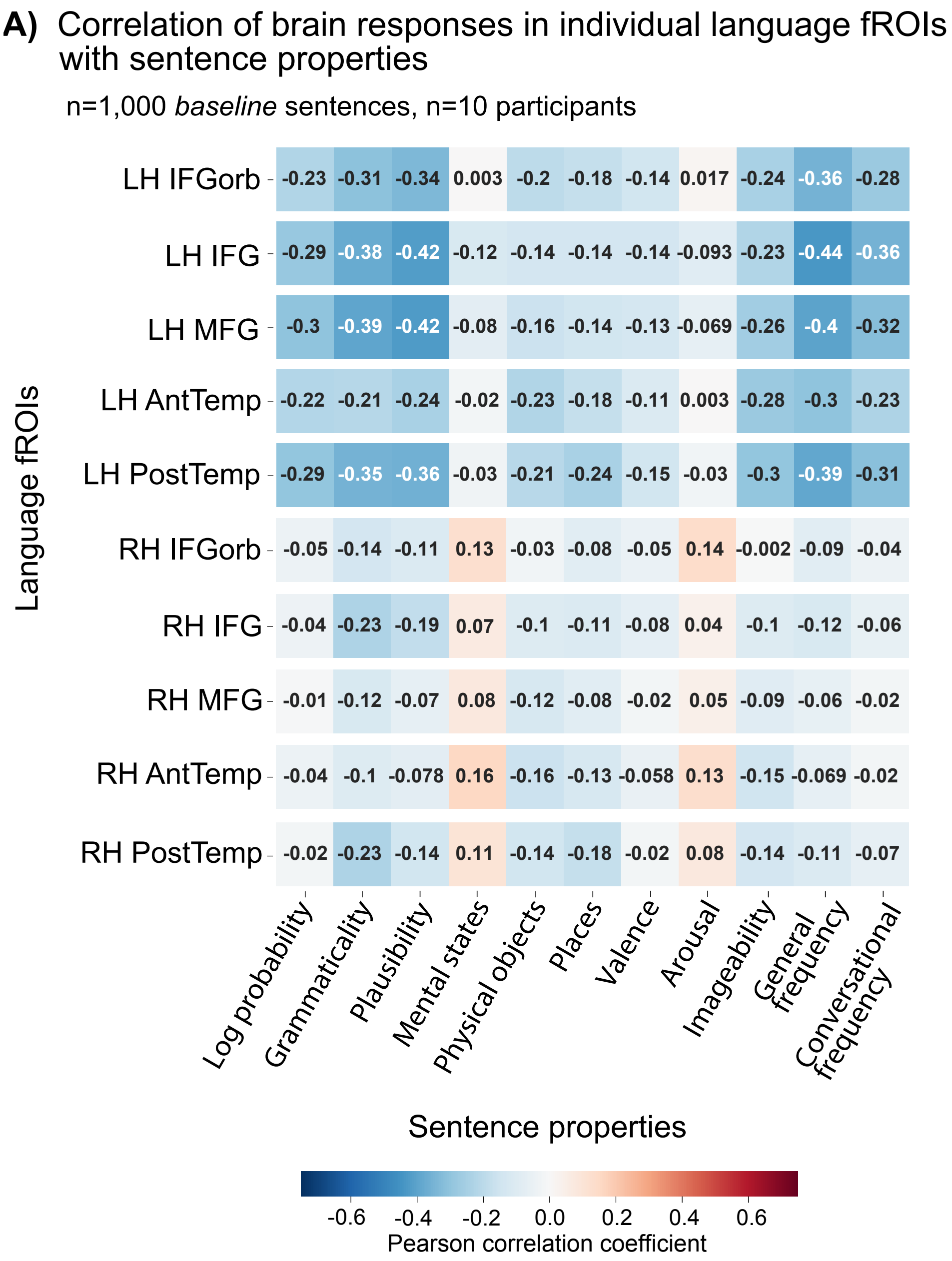

**SI Figure 21A. Modulation of language fROIs' responses by sentence properties.**
Correlation of n=10 language fROIs' (5 in the LH, 5 in the RH) responses (rows) with 11 sentence properties (columns) for n=1,000 *baseline* sentences (averaged across n=5 *train* and n=5 *evaluation* participants).

Next, we quantified the correlation between responses in n=28 anatomical Glasser parcels [36] located in typically language-responsive areas and sentence properties for the n=1,000 *baseline* sentences. **SI Figure 21B** visualizes these correlations on the surface-inflated brain (negative correlations are shown in blue shades, correlations around 0 are not shown, and positive correlations are shown in orange and red shades). First, we note that the correlation magnitudes are substantially lower for the anatomically defined parcels compared to the functional ROIs (**SI Figure 21A**), to be expected given the inter-individual variability in the location of language regions (e.g., [61,87,88,37]). Second, as evident from **SI Figure 21B**, the results for the anatomical parcels mirror the patterns observed for the functional language ROIs reported in Figure 5 in the main text and **SI Figure 21A**: namely, the RH fROIs generally exhibit lower correlations than the LH fROIs, and, in contrast to the LH fROIs, show positive correlations with the “Mental states” and “Arousal” predictors.

**B)** Correlation of BOLD response in Glasser parcels and sentence properties
n=1,000 *baseline* sentences, n=10 participants

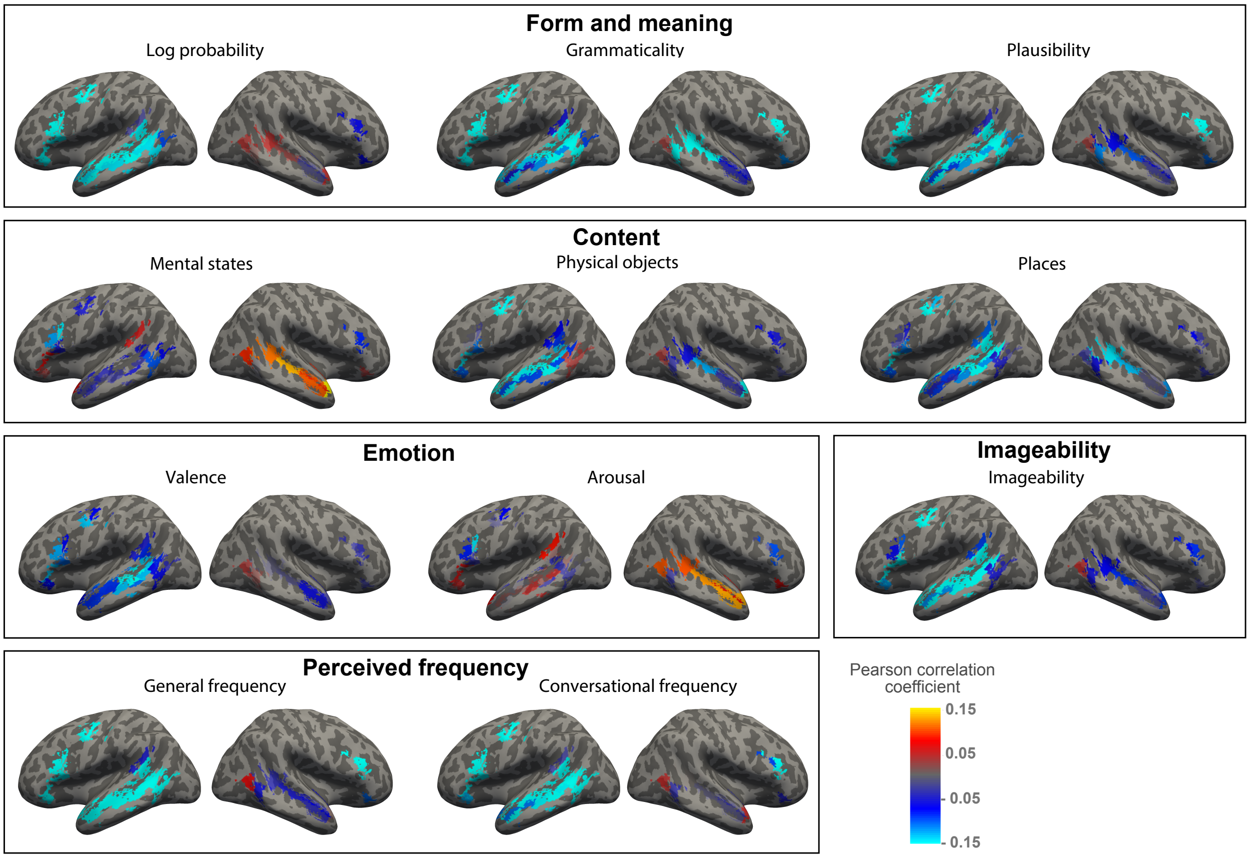

**SI Figure 21B. Modulation of anatomical parcels’ responses by sentence properties.** Correlation of responses in n=28 anatomical Glasser parcels (18 in the LH, 10 in the RH) that are located in language-responsive areas with 11 sentence properties for n=1,000 *baseline* sentences (averaged across n=5 *train* and n=5 *evaluation* participants). To identify parcels in typical language-responsive areas, we identified parcels that overlapped by at least 25% of voxels with one of the five broad, anatomical LH language parcels (Methods; Definition of ROIs), as was done in Lipkin et al. [3], resulting in n=21 parcels in each hemisphere. The names of these Glasser parcels are reported in SI Figure 6D. We excluded 14 of the 42 Glasser parcels (3 in the LH, 11 in the RH) where the noise ceiling split-half reliability (SI 5B) overlapped with zero, which left 28 parcels. The correlation values for each parcel were projected to the surface and visualized on the surface-inflated MNI152 template brain. Negative correlations are shown in blue shades, correlations around 0 are not shown, and positive correlations are shown in orange and red shades.

#### SI 22: Behavioral experiments on sentence properties

##### SI 22A: Participant exclusion criteria

The following criteria were defined prior to the study. Participants were excluded based on:

**1. Native speaker status:** Participants were excluded based on their native speaker status self report as well as the Prolific/mTurk language and location filters.
**2. Sentence completions**: Participants were excluded if their sentence completions were ungrammatical, contained spelling errors (that were not obvious typos) or if the completions were deeply nonsensical.
**3. Average response time:** Participants were excluded if the average response time per question was less than 3 seconds (i.e., for the survey that contained two questions, the threshold was 6 seconds).
**4. Lack of variance in ratings:** Participants were excluded if they only used a total of 2 unique rating values (out of 7) for all items in the survey. In addition, for the 2-question “form and meaning” survey, participants were excluded if they always provided the same rating for two questions across all items.
**5. Correlation with other participants:** Participants were excluded if the average Pearson correlation with the ratings of remaining participants fell below 2 standard deviations below the mean inter-participant correlation. The inter-participant correlated was computed by correlating a vector of responses for a given participant with the vector of responses for each of the remaining participants and taking the average of these pairwise correlation values.

##### SI 22B: Sentence completion prompts

Participants were instructed: “*Please finish the following sentences:*”.

1. The garden that …
2. When I was younger, I would often …
3. The longer the workers protested, ...
4. I could never have imagined that …
5. Because Jane lived by herself, …
6. The most difficult thing about the trip was that ...

##### SI 22C: Experimental instructions and participants

Form and meaning

*Instructions*

In this survey, you will be asked to rate 100 sentences. We would like you to rate each sentence for **two features**.

**First,** we want you to rate the sentence for whether it **makes sense** on a scale from 1 (does not make any sense) to 7 (makes perfect sense). For example, a sentence like *"The father had to buy some new fishing equipment for his trip"* makes sense, so you might rate it as a 6 or 7. In contrast, a sentence like *"The window attacked an inflated red kitten after the long opera surfaced"* does not really make any sense, so you might rate it as a 1 or 2. Some sentences may fall somewhere in between: for example, *"The tired traveller sat down on the book to have a drink of silver*" so you might rate them as a 3 or 4.

**Second,** we want you to rate the sentence for how **grammatical** it is (how well it follows English grammar rules) on a scale from 1 (completely ungrammatical) to 7 (perfectly grammatical). For example, although the sentence *"The window attacked an inflated red kitten after the long opera surfaced"* does not make sense, it obeys grammatical rules, so you might rate this as a 6 or 7. In contrast, a sentence like *"Red the inflated attacked opera window long after kitten surfaced an"* does not obey grammatical rules, so you might rate it as a 1 or 2. Some sentences may fall somewhere in between: for example, *"Father had to bought into some new fishing equipment for his trip*" (here, there are some parts that are grammatical, but also some errors) so you might rate it as a 3 or 4."

Question phrasing for each item:

- How much does the sentence make sense?
- How grammatical is the sentence?

*Participants*

Participants were recruited using the Amazon Mechanical Turk (mTurk) crowd-sourcing platform. The study was restricted to "Mechanical Turk Masters" workers. For lists 8-20, we additionally restricted the study to workers with a HIT approval rate greater than or equal to 98% as well as "Location" set to US using the mTurk qualification filters. 400 participants took part in the experiment. 100 participants were excluded following pre-defined exclusion criteria (SI 21A), leaving 300 participants (75%). In particular, 33 participants were excluded because they listed a language other than English as their native language, 32 participants were excluded based on sentence completions, 3 participants were excluded based on their average response times, 0 participants were excluded based on the lack of rating variance, and 32 participants were excluded based on a low correlation with the remaining participants.

The experiment took 25.01 minutes, on average (SD=3.58). Each item was rated by 10-18 participants (15 participants on average, SD=2.60). The average inter-participant Pearson correlation, computed by correlating a vector of responses for a given participant with the vector of responses for each of the remaining participants and taking the average of these pairwise correlation values, was 0.84 (SD=0.03) for the sense rating and 0.74 (SD=0.04) for the grammaticality rating.

Mental states

*Instructions*

In this survey, you will be asked to read and evaluate 100 sentences. We would like you to rate each sentence according to **how much it made you think of other people's experiences, thoughts, beliefs, desires, and/or emotions** on a scale from 1 (not at all) to 7 (very much).

For example, sentences like *"The woman wondered what made him jealous."* or *"I was so angry that I hit the door."* have meanings that are related to other people's states of mind and emotions so you might rate them as a 6 or a 7. In contrast, sentences like *"A car drove into the parking garage."* or *"Most of earth's surface is covered by water."* do not, so you might rate them as a 1 or a 2.

Question phrasing for each item:

- How much does the sentence make you think of other people's experiences, thoughts, beliefs, desires, and/or emotions?

The Prolific crowdsourcing platform was used to recruit participants for the remaining n=8 surveys.

*Participants*

Participants were asked to rate each sentence (on a scale from 1 to 7) according to how much it made them think of other people's experiences, thoughts, beliefs, desires, and/or emotions (1: not at all, 7: very much).
400 participants took part in the experiment. 83 participants were excluded following pre-defined exclusion criteria, leaving 317 participants (79.25%). In particular, 32 participants were excluded because they listed a language other than English as their native language, 31 participants were excluded based on sentence completions, 1 participant were excluded based on their average response times, 1 participant was excluded based on the lack of rating variance, and 18 participants were excluded based on a low correlation with the remaining participants.
The experiment took 15.73 minutes, on average (SD=2.58). Each item was rated by 14-19 participants (15.85 participants on average, SD=1.57). The average inter-participant Pearson correlation was 0.50 (SD=0.06).

Physical objects

*Instructions*

In this survey, you will be asked to rate 100 sentences. We would like you to rate each sentence according to **how much it made you think of physical objects and/or physical causal interactions** on a scale from 1 (not at all) to 7 (very much).

For example, sentences like *“A few rows of bookshelves came tumbling down.”* or *“There were several large boxes on the floor.”* have meanings that are related to physical objects and/or physical causal interactions, so you might rate them as a 6 or a 7. In contrast, sentences like *“You are very attentive and curious.”* or *“The US justice system consists of three branches of government.”* do not, so you might rate them as a 1 or a 2.

Question phrasing for each item:

- How much does the sentence make you think of physical objects and/or physical causal interactions?

*Participants*

Participants were asked to rate each sentence (on a scale from 1 to 7) according to how much it made them think of physical objects and/or physical causal interactions (1: not at all, 7: very much).
400 participants took part in the experiment. 97 participants were excluded following pre-defined exclusion criteria, leaving 303 participants (75.75%). In particular, 22 participants were excluded because they listed a language other than English as their native language, 52 participants were excluded based on sentence completions, 5 participants were excluded based on their average response times, 1 participant was excluded based on the lack of rating variance, and 17 participants were excluded based on a low correlation with the remaining participants.
The experiment took 13.78 minutes, on average (SD=2.12). Each item was rated by 13-18 participants (15.15 participants on average, SD=1.39). The average inter-participant Pearson correlation was 0.55 (SD=0.10).

Places

*Instructions*

In this survey, you will be asked to rate 100 sentences. We would like you to rate each sentence according to **how much it made you think of places, natural scenes and/or environments** on a scale from 1 (not at all) to 7 (very much).

For example, sentences like *“There are several tall trees in the backyard.”* or *“They walked to the end of the hallway.”* have meanings that are related to places, natural scenes and/or environments, so you might rate them as a 6 or a 7. In contrast, sentences like *“You are very attentive and curious.”* or *“Wool is a type of textile fiber.”* do not, so you might rate them as a 1 or a 2.

Question phrasing for each item:

- How much does the sentence make you think of places, natural scenes and/or environments?

*Participants*

Participants were asked to rate each sentence (on a scale from 1 to 7) according to how much it made them think of places, natural scenes and/or environments (1: not at all, 7: very much).
400 participants took part in the experiment. 95 participants were excluded following pre-defined exclusion criteria, leaving 305 participants (76.25%). In particular, 16 participants were excluded because they listed a language other than English as their native language, 51 participants were excluded based on sentence completions, 5 participants were excluded based on their average response times, 2 participants were excluded based on the lack of rating variance, and 21 participants were excluded based on a low correlation with the remaining participants.
The experiment took 12.74 minutes, on average (SD=2.26). Each item was rated by 12-19 participants (15.25 participants on average, SD=1.94). The average inter-participant Pearson correlation was 0.59 (SD=0.08).

Valence

*Instructions*

In this survey, you will be asked to rate 100 sentences. We would like you to rate each sentence according to **how much it made you feel happy, pleased, content, and/or hopeful** on a scale from 1 (not at all) to 7 (very much).

For example, sentences like *“The puppy snuggles on the couch.”* or *“I celebrated my birthday with my best friends.”* have meanings that are pleasant and positive, so you might rate them as a 6 or a 7. In contrast, sentences like *“The woman was about to vomit.”* or *“The murder took place around midnight.”* do not, so you might rate them as a 1 or a 2.

<u>Question phrasing for each item:</u>

- How much does the sentence make you feel happy, pleased, content, and/or hopeful?

*Participants*

Participants were asked to rate each sentence (on a scale from 1 to 7) according to how much it made them feel happy, pleased, content, and/or hopeful (1: not at all, 7: very much).
400 participants took part in the experiment. 90 participants were excluded following pre-defined exclusion criteria, leaving 310 participants (77.50%). In particular, 18 participants were excluded because they listed a language other than English as their native language, 48 participants were excluded based on sentence completions, 2 participants were excluded based on their average response times, 1 participant was excluded based on the lack of rating variance, and 21 participants were excluded based on a low correlation with the remaining participants.
The experiment took 13.98 minutes, on average (SD=2.09). Each item was rated by 12-19 participants (15.50 participants on average, SD=2.06). The average inter-participant Pearson correlation was 0.54 (SD=0.07).

Arousal

*Instructions*

In this survey, you will be asked to rate 100 sentences. We would like you to rate each sentence according to **how much it made you feel stimulated, excited, frenzied, wide-awake, and/or aroused** on a scale from 1 (not at all) to 7 (very much).

For example, sentences like *"The rollercoaster had tight turns."* or *"His brother slapped him in the face."* have meanings that are arousing, so you might rate them as a 6 or a 7. In contrast, sentences like *"The trees were slowly swaying in the wind."* or *"The woman was sleepy after meditating."* do not, so you might rate them as a 1 or a 2.

Question phrasing for each item:

- How much does the sentence make you feel stimulated, excited, frenzied, wide-awake, and/or aroused?

*Participants*

Participants were asked to rate each sentence (on a scale from 1 to 7) according to how much it made them feel stimulated, excited, frenzied, wide-awake, and/or aroused (1: not at all, 7: very much).
400 participants took part in the experiment. 99 participants were excluded following pre-defined exclusion criteria, leaving 301 participants (75.25%). In particular, 29 participants were excluded because they listed a language other than English as their native language, 46 participants were excluded based on sentence completions, 1 participant was excluded based on their average response times, 6 participants were excluded based on the lack of rating variance, and 17 participants were excluded based on a low correlation with the remaining participants.
The experiment took 15.38 minutes, on average (SD=2.80). Each item was rated by 12-18 participants (15.05 participants on average, SD=1.61). The average inter-participant Pearson correlation was 0.40 (SD=0.05).

Imageability

*Instructions*

In this survey, you will be asked to rate 100 sentences. We would like you to rate each sentence according to **how easy it is to visualize, or to form an image of the sentence's meaning in your mind** on a scale from 1 (not at all) to 7 (very much).

For example, sentences like *"The cup is filled with black coffee."* or *"The girl looked up at the cloud-filled sky."* have meanings that bring to mind relevant images and are easy to visualize, so you might rate them as a 6 or a 7. In contrast, sentences like *"It seems unlikely to happen."* or *"I did not get the gist of the idea."* do not, so you might rate them as a 1 or a 2.

<u>Question phrasing for each item:</u>

- How easy is the sentence to visualize, or to form an image of the sentence's meaning in your mind?

*Participants*

Participants were asked to rate (on a scale from 1 to 7) how easy each sentence is to visualize, or to form an image of the sentence's meaning in their mind (1: not at all, 7: very much).
400 participants took part in the experiment. 105 participants were excluded following pre-defined exclusion criteria, leaving 295 participants (73.75%). In particular, 26 participants were excluded because they listed a language other than English as their native language, 58 participants were excluded based on sentence completions, 2 participants were excluded based on their average response times, 1 participant was excluded based on the lack of rating variance, and 18 participants were excluded based on a low correlation with the remaining participants.
The experiment took 14.64 minutes, on average (SD=2.12). Each item was rated by 12-19 participants (14.75 participants on average, SD=1.59). The average inter-participant Pearson correlation was 0.63 (SD=0.04).

Perceived general frequency

*Instructions*

In this survey, you will be asked to rate 100 sentences. We would like you to rate each sentence according to **how likely you think you are to encounter this sentence** on a scale from 1 (not at all likely) to 7 (very likely).

For example, sentences like *"I want my coffee black, please."* or *"What time is our meeting?"* are quite common, so you might rate them as a 6 or a 7. In contrast, sentences like *"The man measures the height of the tripod." or "Five prophets from Egypt were present."* are not, so you might rate them as a 1 or a 2.

<u>Question phrasing for each item:</u>

- How likely do you think you are to encounter this sentence?

*Participants*

Participants were asked to rate each sentence (on a scale from 1 to 7) according to how likely they think they are to encounter the sentence (1: not at all likely, 7: very likely).
400 participants took part in the experiment. 85 participants were excluded following pre-defined exclusion criteria, leaving 315 participants (78.75%). In particular, 31 participants were excluded because they listed a language other than English as their native language, 31 participants were excluded based on sentence completions, 1 participant was excluded based on their average

response times, 0 participants were excluded based on the lack of rating variance, and 22 participants were excluded based on a low correlation with the remaining participants.
The experiment took 14.27 minutes, on average (SD=2.23). Each item was rated by 13-19 participants (15.75 participants on average, SD=1.55). The average inter-participant Pearson correlation was 0.72 (SD=0.05).

Perceived conversational frequency

*Instructions*

In this survey, you will be asked to rate 100 sentences. We would like you to rate each sentence according to **how likely you think it is to occur in a conversation between people** on a scale from 1 (not at all likely) to 7 (very likely).

For example, sentences like *“I love your shoes!”* or *“What did you guys do last night?.”* are likely to be said during a conversation, so you might rate them as a 6 or a 7. In contrast, sentences like *“Hemoglobin is a protein that carries oxygen.”* or *“Land owners do not obtain any tax treaty.”* are not and instead are more likely to occur in written texts, so you might rate them as a 1 or a 2.

Question phrasing for each item:

- How likely do you think the sentence is to occur in a conversation between people?

*Participants*

Participants were asked to rate each sentence (on a scale from 1 to 7) according to how likely they think the sentence is to occur in a conversation between people (1: not at all, 7: very much).
400 participants took part in the experiment. 105 participants were excluded following pre-defined exclusion criteria, leaving 295 participants (73.75%). In particular, 37 participants were excluded because they listed a language other than English as their native language, 48 participants were excluded based on sentence completions, 3 participants were excluded based on their average response times, 1 participant was excluded based on the lack of rating variance, and 16 participants were excluded based on a low correlation with the remaining participants.
The experiment took 13.47 minutes, on average (SD=1.39). Each item was rated by 12-18 participants (14.75 participants on average, SD=1.68). The average inter-participant Pearson correlation was 0.64 (SD=0.05).

#### SI 23: Statistical significance of the effect of sentence properties on the BOLD response

The following section contains statistics accompanying Results; Sentence complexity modulates language network responses in the main text. Section SI 23A contains full model formulae along with model fit statistics, while section SI 23B contains statistics comparing pairs of models. BOLD response data met assumptions of normality (tested via Kolmogorov-Smirnov test).

##### SI 23A: Individual linear mixed effect models

Log probability

Linear mixed effect formula:
*BOLD response ~ log_probability + (1 | sentence) + run_within_session + trial_within_run*

| ***Model term*** | ***Coefficient*** | ***SE*** | ***t*** | ***p*** | ***df*** |
|---:|---|---|---|---|---|
| *intercept* | -0.333*** | 0.044 | -7.625 | <0.001 | 9982 |
| *log_probability* | -0.085*** | 0.009 | -9.937 | <0.001 | 9982 |
| *run_within_session* | -0.002 | 0.002 | -1.100 | 0.271 | 9982 |
| *trial_within_run* | -0.002*** | 0.000 | -4.809 | <0.001 | 9982 |
| *SD (Intercept item_id_factor)* | 0.168 | | | | |
| *SD (Observations)* | 0.629 | | | | |
| *Num.Obs.* | 9988 | | | | |
| *R2 Marg.* | 0.018 | | | | |
| *R2 Cond.* | 0.083 | | | | |
| *AIC* | 19658.6 | | | | |
| *BIC* | 19701.9 | | | | |
| *RMSE* | 0.62 | | | | |

Log probability and plausibility

Linear mixed effect formula:
*BOLD response ~ log_probability + plausibility + (1 | sentence) + run_within_session + trial_within_run*

| ***Model term*** | ***Coefficient*** | ***SE*** | ***t*** | ***p*** | ***df*** |
|---|---|---|---|---|---|
| *intercept* | 0.553*** | 0.106 | 5.210 | <0.001 | 9981 |
| *log_probability* | -0.044*** | 0.009 | -4.693 | <0.001 | 9981 |
| *plausibility* | -0.108*** | 0.012 | -9.086 | <0.001 | 9981 |
| *run_within_session* | -0.003 | 0.002 | -1.139 | 0.255 | 9981 |
| *trial_within_run* | -0.002*** | 0.000 | -4.711 | <0.001 | 9981 |
| *SD (Intercept item_id_factor)* | 0.151 | | | | |
| *SD (Observations)* | 0.629 | | | | |
| *Num.Obs.* | 9988 | | | | |
| *R2 Marg.* | 0.030 | | | | |
| *R2 Cond.* | 0.083 | | | | |
| *AIC* | 19588.5 | | | | |
| *BIC* | 19638.9 | | | | |
| *RMSE* | 0.62 | | | | |

Log probability, plausibility, and grammaticality

Linear mixed effect formula:
*BOLD response ~ log_probability + plausibility + grammaticality + (1 | sentence) + run_within_session + trial_within_run*

| ***Model term*** | ***Coefficient*** | ***SE*** | ***t*** | ***p*** | ***df*** |
|---|---|---|---|---|---|
| *intercept* | 0.0672*** | 0.111 | 6.081 | <0.001 | 9980 |
| *log_probability* | -0.042*** | 0.009 | -4.476 | <0.001 | 9980 |

| | | | | | |
|---|---|---|---|---|---|
| *plausibility* | -0.068*** | 0.016 | -4.246 | <0.001 | 9980 |
| *grammaticality* | -0.058*** | 0.016 | -3.613 | <0.001 | 9980 |
| *run_within_session* | -0.003 | 0.002 | -1.151 | 0.250 | 9980 |
| *trial_within_run* | -0.002*** | 0.000 | -4.700 | <0.001 | 9980 |
| *SD (Intercept item_id_factor)* | 0.149 | | | | |
| *SD (Observations)* | 0.629 | | | | |
| *Num.Obs.* | 9988 | | | | |
| *R2 Marg.* | 0.032 | | | | |
| *R2 Cond.* | 0.083 | | | | |
| *AIC* | 19584.0 | | | | |
| *BIC* | 19641.6 | | | | |
| *RMSE* | 0.62 | | | | |

Log probability and grammaticality

Linear mixed effect formula:
*BOLD response ~ log_probability + grammaticality + (1 | sentence) + run_within_session + trial_within_run*

| ***Model term*** | ***Coefficient*** | ***SE*** | ***t*** | ***p*** | ***df*** |
|---|---|---|---|---|---|
| *intercept* | 0.466*** | 0.100 | 4.651 | <0.001 | 9981 |
| *log_probability* | -0.054*** | 0.009 | -6.020 | <0.001 | 9981 |
| *grammaticality* | -0.104** | 0.012 | -8.790 | <0.001 | 9981 |
| *run_within_session* | -0.003 | 0.002 | -1.146 | 0.252 | 9981 |
| *trial_within_run* | -0.002*** | 0.000 | -4.727 | <0.001 | 9981 |

| | |
|---|---|
| *SD (Intercept item_id_factor)* | 0.152 |
| *SD (Observations)* | 0.629 |
| *Num.Obs.* | 9988 |
| *R2 Marg.* | 0.029 |
| *R2 Cond.* | 0.083 |
| *AIC* | 19593.4 |
| *BIC* | 19643.8 |
| *RMSE* | 0.62 |

Log probability and mental states

Linear mixed effect formula:

*BOLD response ~ log_probability + mental_states + (1 | sentence) + run_within_session + trial_within_run*

| **Model term** | **Coefficient** | **SE** | **t** | **p** | **df** |
|---|---|---|---|---|---|
| *intercept* | -0.361*** | 0.055 | -6.521 | <0.001 | 9981 |
| *log_probability* | -0.087*** | 0.009 | -9.861 | <0.001 | 9981 |
| *mental_states* | 0.005 | 0.006 | 0.829 | 0.407 | 9981 |
| *run_within_session* | -0.002 | 0.002 | -1.094 | 0.274 | 9981 |
| *trial_within_run* | -0.002*** | 0.000 | -4.813 | <0.001 | 9981 |
| *SD (Intercept item_id_factor)* | 0.167 | | | | |
| *SD (Observations)* | 0.629 | | | | |
| *Num.Obs.* | 9988 | | | | |
| *R2 Marg.* | 0.018 | | | | |
| *R2 Cond.* | 0.083 | | | | |
| *AIC* | 19668.2 | | | | |
| *BIC* | 19718.6 | | | | |
| *RMSE* | 0.62 | | | | |

Log probability and physical objects

Linear mixed effect formula:

*BOLD response ~ log_probability + physical_objects + (1 | sentence) + run_within_session + trial_within_run*

| ***Model term*** | ***Coefficient*** | ***SE*** | ***t*** | ***p*** | ***df*** |
|---|---|---|---|---|---|
| *intercept* | -0.239*** | 0.044 | -5.496 | <0.001 | 9981 |
| *log_probability* | -0.095*** | 0.008 | -11.341 | <0.001 | 9981 |
| *physical_objects* | -0.047*** | 0.005 | -8.780 | <0.001 | 9981 |
| *run_within_session* | -0.002 | 0.002 | -1.007 | 0.314 | 9981 |
| *trial_within_run* | -0.002*** | 0.000 | -4.708 | <0.001 | 9981 |
| *SD (Intercept item_id_factor)* | 0.152 | | | | |
| *SD (Observations)* | 0.629 | | | | |
| *Num.Obs.* | 9988 | | | | |
| *R2 Marg.* | 0.029 | | | | |
| *R2 Cond.* | 0.083 | | | | |
| *AIC* | 19595.1 | | | | |
| *BIC* | 19645.6 | | | | |
| *RMSE* | 0.62 | | | | |

Log probability and places

Linear mixed effect formula:

*BOLD response ~ log_probability + places + (1 | sentence) + run_within_session + trial_within_run*

| ***Model term*** | ***Coefficient*** | ***SE*** | ***t*** | ***p*** | ***df*** |
|---|---|---|---|---|---|
| *intercept* | -0.240*** | 0.044 | -5.473 | <0.001 | 9981 |
| *log_probability* | -0.090*** | 0.008 | -10.742 | <0.001 | 9981 |

| | | | | | |
|---|---|---|---|---|---|
| *places* | -0.053*** | 0.007 | -8.095 | <0.001 | 9981 |
| *run_within_session* | -0.003 | 0.002 | -1.143 | 0.253 | 9981 |
| *trial_within_run* | -0.002*** | 0.000 | -4.673 | <0.001 | 9981 |
| *SD (Intercept item_id_factor)* | 0.155 | | | | |
| *SD (Observations)* | 0.629 | | | | |
| *Num.Obs.* | 9988 | | | | |
| *R2 Marg.* | 0.027 | | | | |
| *R2 Cond.* | 0.083 | | | | |
| *AIC* | 19605.5 | | | | |
| *BIC* | 19655.9 | | | | |
| *RMSE* | 0.62 | | | | |

Log probability and valence

Linear mixed effect formula:
*BOLD response ~ log_probability + valence + (1 | sentence) + run_within_session + trial_within_run*

| ***Model term*** | ***Coefficient*** | ***SE*** | ***t*** | ***p*** | ***df*** |
|---|---|---|---|---|---|
| *intercept* | -0.239*** | 0.049 | -4.872 | <0.001 | 9981 |
| *log_probability* | -0.082*** | 0.009 | -9.627 | <0.001 | 9981 |
| *valence* | -0.026*** | 0.006 | -4.083 | <0.001 | 9981 |
| *run_within_session* | -0.003 | 0.002 | -1.121 | 0.263 | 9981 |
| *trial_within_run* | -0.002*** | 0.000 | -4.829 | <0.001 | 9981 |
| *SD (Intercept item_id_factor)* | 0.164 | | | | |

| | |
|---|---|
| *SD (Observations)* | 0.629 |
| *Num.Obs.* | 9988 |
| *R2 Marg.* | 0.020 |
| *R2 Cond.* | 0.083 |
| *AIC* | 19652.4 |
| *BIC* | 19702.8 |
| *RMSE* | 0.62 |

Log probability and arousal

Linear mixed effect formula:

*BOLD response ~ log_probability + arousal +  (1 | sentence) + run_within_session + trial_within_run*

| Model term | Coefficient | SE | t | p | df |
|---|---|---|---|---|---|
| *intercept* | -0.347*** | 0.054 | -6.369 | <0.001 | 9981 |
| *log_probability* | -0.086*** | 0.009 | -9.896 | <0.001 | 9981 |
| *arousal* | 0.004 | 0.009 | 0.429 | 0.668 | 9981 |
| *run_within_session* | -0.002 | 0.002 | -1.097 | 0.273 | 9981 |
| *trial_within_run* | -0.002*** | 0.000 | -4.811 | <0.001 | 9981 |
| *SD (Intercept item_id_factor)* | 0.168 | | | | |
| *SD (Observations)* | 0.629 | | | | |
| *Num.Obs.* | 9988 | | | | |
| *R2 Marg.* | 0.018 | | | | |
| *R2 Cond.* | 0.083 | | | | |
| *AIC* | 19668.1 | | | | |
| *BIC* | 19718.6 | | | | |
| *RMSE* | 0.62 | | | | |

+p < 0.1, *p < 0.05, **p < 0.01, ***p < 0.001

Log probability and imageability

Linear mixed effect formula:

*BOLD response ~ log_probability + imageability + (1 | sentence) + run_within_session + trial_within_run*

| Model term | Coefficient | SE | t | p | df |
|---|---|---|---|---|---|
| intercept | -0.094+ | 0.048 | -1.941 | 0.052 | 9981 |
| log_probability | -0.081*** | 0.008 | -9.798 | <0.001 | 9981 |
| imageability | -0.058*** | 0.006 | -9.874 | <0.001 | 9981 |
| run_within_session | -0.002 | 0.002 | -1.034 | 0.301 | 9981 |
| trial_within_run | -0.002*** | 0.000 | -4.744 | <0.001 | 9981 |
| SD (Intercept item_id_factor) | 0.149 | | | | |
| SD (Observations) | 0.629 | | | | |
| Num.Obs. | 9988 | | | | |
| R2 Marg. | 0.032 | | | | |
| R2 Cond. | 0.083 | | | | |
| AIC | 19576.2 | | | | |
| BIC | 19626.7 | | | | |
| RMSE | 0.62 | | | | |

Log probability and perceived frequency

Linear mixed effect formula:

*BOLD response ~ log_probability + general_frequency + (1 | sentence) + run_within_session + trial_within_run*

| Model term | Coefficient | SE | t | p | df |
|---|---|---|---|---|---|
| intercept | 0.264*** | 0.073 | 3.642 | <0.001 | 9981 |
| log_probability | -0.028*** | 0.010 | -2.844 | <0.004 | 9981 |
| general_frequency | -0.075*** | 0.007 | -10.071 | <0.001 | 9981 |
| run_within_session | -0.003 | 0.002 | -1.128 | 0.259 | 9981 |
| trial_within_run | -0.002*** | 0.000 | -4.706 | <0.001 | 9981 |
| SD (Intercept item_id_factor) | 0.148 | | | | |

| | |
|---|---|
| *SD (Observations)* | 0.629 |
| *Num.Obs.* | 9988 |
| *R2 Marg.* | 0.032 |
| *R2 Cond.* | 0.083 |
| *AIC* | 19572.1 |
| *BIC* | 19622.6 |
| *RMSE* | 0.62 |

Log probability and perceived conversational frequency

Linear mixed effect formula:
*BOLD response ~ log_probability + conversational_frequency + (1 | sentence) + run_within_session + trial_within_run*

| **Model term** | ***Coefficient*** | ***SE*** | ***t*** | ***p*** | ***df*** |
|---|---|---|---|---|---|
| *intercept* | 0.039 | 0.070 | 0.557 | 0.578 | 9981 |
| *log_probability* | -0.051*** | 0.010 | -5.150 | <0.001 | 9981 |
| *conversational_frequency* | -0.048*** | 0.007 | -6.742 | <0.001 | 9981 |
| *run_within_session* | -0.003 | 0.002 | -1.120 | 0.263 | 9981 |
| *trial_within_run* | -0.002*** | 0.000 | -4.765 | <0.001 | 9981 |
| *SD (Intercept item_id_factor)* | 0.159 | | | | |
| *SD (Observations)* | 0.629 | | | | |
| *Num.Obs.* | 9988 | | | | |
| *R2 Marg.* | 0.025 | | | | |
| *R2 Cond.* | 0.083 | | | | |
| *AIC* | 19624.3 | | | | |
| *BIC* | 19764.8 | | | | |
| *RMSE* | 0.62 | | | | |

#### SI 23B: Comparison of pairs of linear mixed effect models

Grammaticality versus plausibility and grammaticality

Note that log probability was included in both of these models as a base predictor.

Model 1: *BOLD response ~ log_probability + grammaticality + (1 | sentence) + run_within_session + trial_within_run*

Model 2: *BOLD response ~ log_probability + plausibility + grammaticality + (1 | sentence) + run_within_session + trial_within_run*

| npar | AIC | BIC | logLik | deviance | Chisq | Df | Pr(>Chisq) |
|---|---|---|---|---|---|---|---|
| 7 | 19 546.78 | 19 597.35 | -9 766.391 | 19 532.78 | na | na | na |
| 8 | 19 530.92 | 19 588.59 | -9 757.459 | 19 514.92 | 17.86499 | 1 | 0.0000237 |

Plausibility versus plausibility and grammaticality

Note that log probability was included in both of these models as a base predictor.

Model 1: *BOLD response ~ log_probability + plausibility + (1 | sentence) + run_within_session + trial_within_run*

Model 2: *BOLD response ~ log_probability + plausibility + grammaticality + (1 | sentence) + run_within_session + trial_within_run*

| npar | AIC | BIC | logLik | deviance | Chisq | Df | Pr(>Chisq) |
|---|---|---|---|---|---|---|---|
| 7 | 19 541.89 | 19 592.35 | -9 763.944 | 19 527.89 | na | na | na |
| 8 | 19 530.92 | 19 588.59 | -9 757.459 | 19 514.92 | 12.97034 | 1 | 0.00031647 |

Log probability versus mental states

Model 1: *BOLD response ~ log_probability + (1 | sentence) + run_within_session + trial_within_run*

Model 2: *BOLD response ~ log_probability + mental_states + (1 | sentence) + run_within_session + trial_within_run*

| npar | AIC | BIC | logLik | deviance | Chisq | Df | Pr(>Chisq) |
|---|---|---|---|---|---|---|---|
| 6 | 19 619.22 | 19 662.47 | -9 803.609 | 19 607.22 | na | na | na |
| 7 | 19 620.53 | 19 671.00 | -9 803.266 | 19 606.53 | 0.6870964 | 1 | 0.4071538 |

Log probability versus physical objects

Model 1: *BOLD response ~ log_probability + (1 | sentence) + run_within_session + trial_within_run*

Model 2: *BOLD response ~ log_probability + physical_objects + (1 | sentence) + run_within_session + trial_within_run*

| npar | AIC | BIC | logLik | deviance | Chisq | Df | Pr(>Chisq) |
|---|---|---|---|---|---|---|---|
| 6 | 19 619.22 | 19 662.47 | -9 803.609 | 19 607.22 | na | na | na |
| 7 | 19 546.96 | 19 597.42 | -9 766.479 | 19 532.96 | 74.26132 | 1 | 0 |

Log probability versus places

Model 1: *BOLD response ~ log_probability + (1 | sentence) + run_within_session + trial_within_run*

Model 2: *BOLD response ~ log_probability + places + (1 | sentence) + run_within_session + trial_within_run*

| npar | AIC | BIC | logLik | deviance | Chisq | Df | Pr(>Chisq) |
|---|---|---|---|---|---|---|---|
| 6 | 19 619.22 | 19 662.47 | -9 803.609 | 19 607.22 | na | na | na |
| 7 | 19 557.75 | 19 608.21 | -9 771.873 | 19 543.75 | 63.47272 | 1 | 0 |

Log probability versus valence

Model 1: *BOLD response ~ log_probability + (1 | sentence) + run_within_session + trial_within_run*

Model 2: *BOLD response ~ log_probability + places + (1 | sentence) + run_within_session + trial_within_run*

| npar | AIC | BIC | logLik | deviance | Chisq | Df | Pr(>Chisq) |
|---|---|---|---|---|---|---|---|
| 6 | 19 619.22 | 19 662.47 | -9 803.609 | 19 607.22 | na | na | na |
| 7 | 19 604.69 | 19 655.15 | -9 795.343 | 19 590.69 | 16.53291 | 1 | 0.000047813 |

Log probability versus arousal

Model 1: *BOLD response ~ log_probability + (1 | sentence) + run_within_session + trial_within_run*

Model 2: *BOLD response ~ log_probability + arousal + (1 | sentence) + run_within_session + trial_within_run*

| npar | AIC | BIC | logLik | deviance | Chisq | Df | Pr(>Chisq) |
|---|---|---|---|---|---|---|---|
| 6 | 19 619.22 | 19 662.47 | -9 803.609 | 19 607.22 | na | na | na |
| 7 | 19 621.03 | 19 671.50 | -9 803.517 | 19 607.03 | 0.1842934 | 1 | 0.6677092 |

Log probability versus imageability

Model 1: *BOLD response ~ log_probability + (1 | sentence) + run_within_session + trial_within_run*

Model 2: *BOLD response ~ log_probability + imageability + (1 | sentence) + run_within_session + trial_within_run*

| *npar* | *AIC* | *BIC* | *logLik* | *deviance* | *Chisq* | *Df* | *Pr(>Chisq)* |
|---|---|---|---|---|---|---|---|
| 6 | 19 619.22 | 19 662.47 | -9 803.609 | 19 607.22 | na | na | na |
| 7 | 19 528.19 | 19 578.65 | -9 757.094 | 19 514.19 | 93.03145 | 1 | 0 |

Log probability versus perceived frequency

Model 1: *BOLD response ~ log_probability + (1 | sentence) + run_within_session + trial_within_run*

Model 2: *BOLD response ~ log_probability + general_frequency + (1 | sentence) + run_within_session + trial_within_run*

| *npar* | *AIC* | *BIC* | *logLik* | *deviance* | *Chisq* | *Df* | *Pr(>Chisq)* |
|---|---|---|---|---|---|---|---|
| 6 | 19 619.22 | 19 662.47 | -9 803.609 | 19 607.22 | na | na | na |
| 7 | 19 524.59 | 19 575.05 | -9 755.295 | 19 510.59 | 96.62785 | 1 | 0 |

Log probability versus perceived conversational frequency

Model 1: *BOLD response ~ log_probability + (1 | sentence) + run_within_session + trial_within_run*

Model 2: *BOLD response ~ log_probability + conversational_frequency + (1 | sentence) + run_within_session + trial_within_run*

| *npar* | *AIC* | *BIC* | *logLik* | *deviance* | *Chisq* | *Df* | *Pr(>Chisq)* |
|---|---|---|---|---|---|---|---|
| 6 | 19 619.22 | 19 662.47 | -9 803.609 | 19 607.22 | na | na | na |
| 7 | 19 576.76 | 19 627.22 | -9 781.380 | 19 562.76 | 44.45788 | 1 | 0 |

#### SI 24: Statistical differences between the BOLD response for pairs of sentence property bins

The following section contains statistics accompanying the inset plots in Figure 5C in the main text results section Sentence complexity modulates language network responses (the inset plots in Figure 5C show the average brain response for each sentence property grouped into six uniformly sized bins). **SI Figure 24** shows the p-values from independent samples t-tests between each pair of bins, for each sentence property.

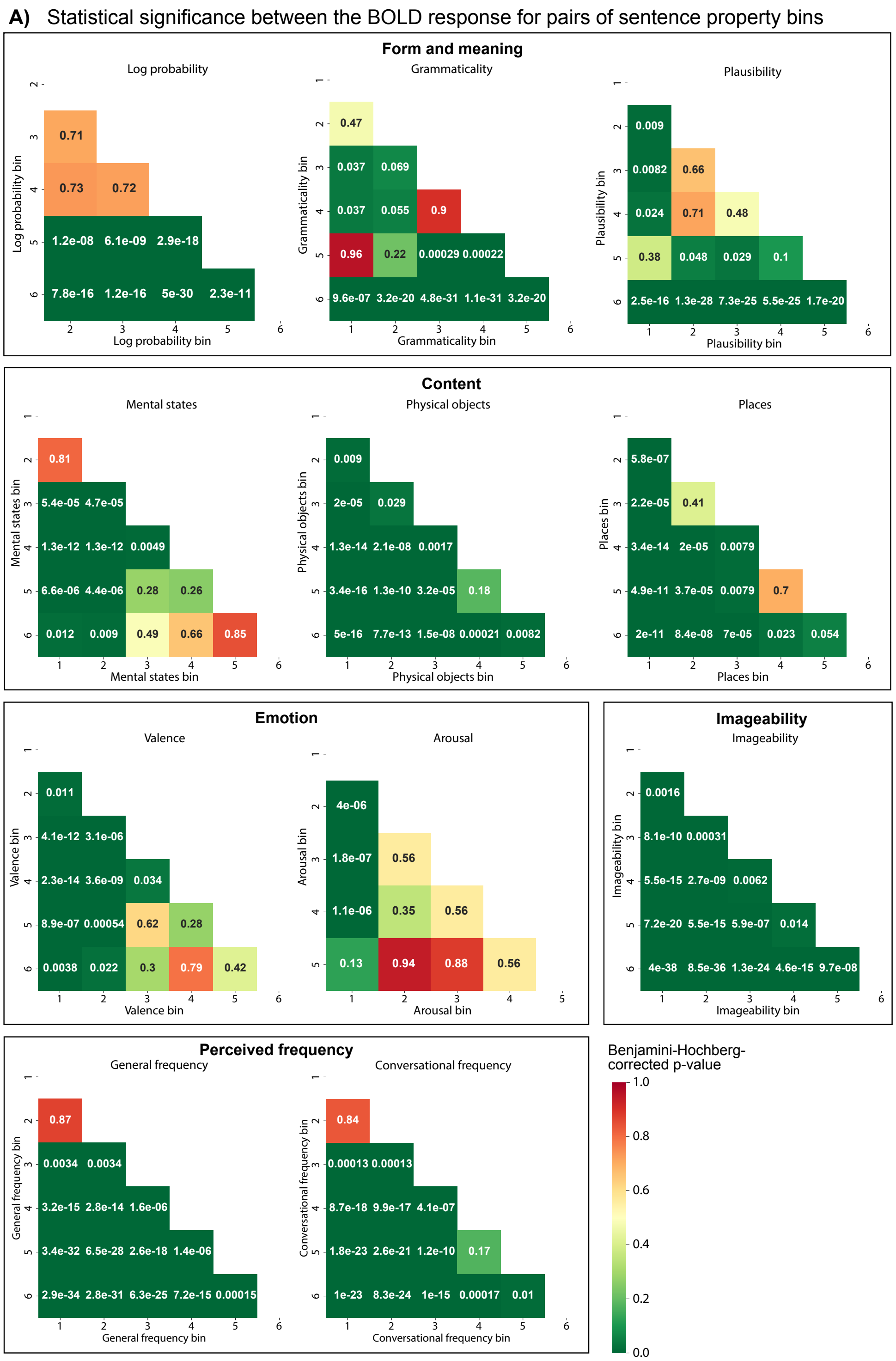

**SI Figure 24. Statistical differences between the BOLD response for pairs of sentence property bins for each of 11 sentence properties.**

Plots show the p-value from independent samples t-tests (two-sided) between each pair of sentence property bins. Mirroring the bin inset plots (Figure 5C, main text), bins containing less than 1% of the data, i.e., 20 responses, were omitted from the analyses. We performed false discovery rate correction via the Benjamini-Hochberg procedure for the comparisons within each sentence property. We show these corrected p-values as a grid containing the unique pairwise comparisons.
